# Six new reference-quality bat genomes illuminate the molecular basis and evolution of bat adaptations

**DOI:** 10.1101/836874

**Authors:** David Jebb, Zixia Huang, Martin Pippel, Graham M. Hughes, Ksenia Lavrichenko, Paolo Devanna, Sylke Winkler, Lars S. Jermiin, Emilia C. Skirmuntt, Aris Katzourakis, Lucy Burkitt-Gray, David A. Ray, Kevin A. M. Sullivan, Juliana G. Roscito, Bogdan M. Kirilenko, Liliana M. Dávalos, Angelique P. Corthals, Megan L. Power, Gareth Jones, Roger D. Ransome, Dina Dechmann, Andrea G. Locatelli, Sebastien J. Puechmaille, Olivier Fedrigo, Erich D. Jarvis, Mark S. Springer, Michael Hiller, Sonja C. Vernes, Eugene W. Myers, Emma C. Teeling

## Abstract

Bats account for ~20% of all extant mammal species and are considered exceptional given their extraordinary adaptations, including biosonar, true flight, extreme longevity, and unparalleled immune systems. To understand these adaptations, we generated reference-quality genomes of six species representing the key divergent lineages. We assembled these genomes with a novel pipeline incorporating state-of-the-art long-read and long-range sequencing and assembly techniques. The genomes were annotated using a maximal evidence approach, *de novo* predictions, protein/mRNA alignments, Iso-seq long read and RNA-seq short read transcripts, and gene projections from our new TOGA pipeline, retrieving virtually all (>99%) mammalian BUSCO genes. Phylogenetic analyses of 12,931 protein coding-genes and 10,857 conserved non-coding elements identified across 48 mammalian genomes helped to resolve bats’ closest extant relatives within Laurasiatheria, supporting a basal position for bats within Scrotifera. Genome-wide screens along the bat ancestral branch revealed (a) selection on hearing-involved genes (e.g *LRP2, SERPINB6, TJP2)*, which suggest that laryngeal echolocation is a shared ancestral trait of bats; (b) selection (e.g *INAVA, CXCL13, NPSR1*) and loss of immunity related proteins (e.g. *LRRC70, IL36G*), including pro-inflammatory NF-kB signalling; and (c) expansion of the *APOBEC* family, associated with restricting viral infection, transposon activity and interferon signalling. We also identified unique integrated viruses, indicating that bats have a history of tolerating viral pathogens, lethal to other mammal species. Non-coding RNA analyses identified variant and novel microRNAs, revealing regulatory relationships that may contribute to phenotypic diversity in bats. Together, our reference-quality genomes, high-quality annotations, genome-wide screens and *in-vitro* tests revealed previously unknown genomic adaptations in bats that may explain their extraordinary traits.

## Introduction

With more than ~1400 species identified to date^1^, bats (Chiroptera) account for ~20% of all currently recognised, extant, mammal species, are found around the globe, and successfully occupy diverse ecological niches^1,2^. Their global success is attributed to their extraordinary suite of adaptations including: powered flight, laryngeal echolocation for orientation and hunting in complete darkness, exceptional longevity, and a unique immune system that enables bats to tolerate viruses that are typically lethal in other mammals (e.g., rabies, SARS, MERS)^2^. It has been proposed that the evolution of extended longevity and immunity in bats was driven by the acquisition of flight, which has a high metabolic cost^3–5^, but the mechanisms underlying these adaptations are unknown and their potential connection to flight is still debated^6,7^. Given bats’ distinctive adaptations, they represent important model systems to uncover the molecular basis and evolution of extended healthspan^7,8^, enhanced disease tolerance^9^ and sensory perception^10,11^. To understand the evolution of such traits, one needs to understand bats’ evolutionary history. However, key aspects of that evolutionary history such as monophyly of echolocating bats and the single origin of laryngeal echolocation^10,12^ remain debated, partially stemming from a poor fossil record^13^, incongruent phylogenetic analyses^14^, and importantly the limited quality of available genome assemblies.

Here, we generated the first reference-quality genomes of six bats as part of the Bat1K global genome consortium^2^ (http://bat1k.com) in coordination with the Vertebrate Genome Project (https://vertebrategenomesproject.org/). Species were chosen to enable capture of the major ecological trait space and life histories observed in bats while representing deep phylogenetic divergences. These six bat species belong to five families that represent key evolutionary clades, unique adaptations and span both major lineages in Chiroptera estimated to have diverged ~64 MYA^15^. In the suborder Yinpterochiroptera we sequenced *Rhinolophus ferrumequinum* (Greater horseshoe bat; family Rhinolophidae) and *Rousettus aegyptiacus* (Egyptian fruit bat; Pteropodidae), and in Yangochiroptera we sequenced *Phyllostomus discolor* (Pale spear-nose bat; Phyllostomidae), *Myotis myotis* (Greater mouse-eared bat; Vespertilionidae), *Pipistrellus kuhlii* (Kuhl’s pipistrelle; Vespertilionidae) and *Molossus molossus* (Velvety free-tailed bat; Molossidae) (Table S1). These bat genera represent the extremes in known bat longevity^16^. They also represent major adaptations in bat sensory perception and ecological diversity^2^, and include species considered key viral reservoirs and asymptomatic hosts^9,17^.

To obtain genome assemblies of high contiguity and completeness, we developed novel pipelines incorporating state-of-the-art sequencing and assembly. To ascertain the position of Chiroptera within Laurasiatheria and thus resolve a long-standing phylogenetic debate^14^, we mined these near complete genomes to produce a comprehensive orthologous gene data set (12,931), including data from 42 other representative mammalian genomes (TableS1), and applied a suite of diverse phylogenetic approaches. To identify molecular changes in regions of the genome - both coding and non-coding - that underlie bat adaptations, we carried out selection tests, analysed gains and losses of genes, and experimentally validated novel bat microRNAs. We focussed on assessing the shared commonalities between the bat species enabling us to infer the ancestral selection driving key bat adaptations. We elucidated the diversity of endogenous viruses contained within the bat genomes, exploring bats’ putative history with these viruses. Herein, we present the first six reference-quality bat genomes, which we make available in the open access Bat1K browser (also available on NCBI and GenomeArk) and demonstrate both the value of highly contiguous and highly complete genomes and the utility of bats as model organisms to address fundamental questions in biology.

### Genome Sequencing and Assembly

For each of the six bats, we generated: (i) PacBio long reads (52-70X in reads ≥4 kb; N50 read length 14.9-24.5 kb), (ii) 10x Genomics Illumina read clouds (43-104X), (iii) Bionano optical maps (coverage of molecules ≥150 kb 89-288X), and (iv) Hi-C Illumina read pairs (15-95X). PacBio reads were assembled into contigs with a customized assembler called DAmar, a hybrid of our earlier Marvel^18^, Dazzler (https://dazzlerblog.wordpress.com/), and Daccord^19,20^ systems (Fig. 1a). Next, we used PacBio reads and 10x read cloud Illumina data to remove base errors, which was followed by identifying and phasing all regions of the contigs that had a sufficient rate of haplotype heterogeneity. We retained one haplotype for each region, yielding primary contigs. These primary contigs were then scaffolded using a Bionano optical map and the Hi-C data (see supplementary methods section 2).

**Figure 1:**
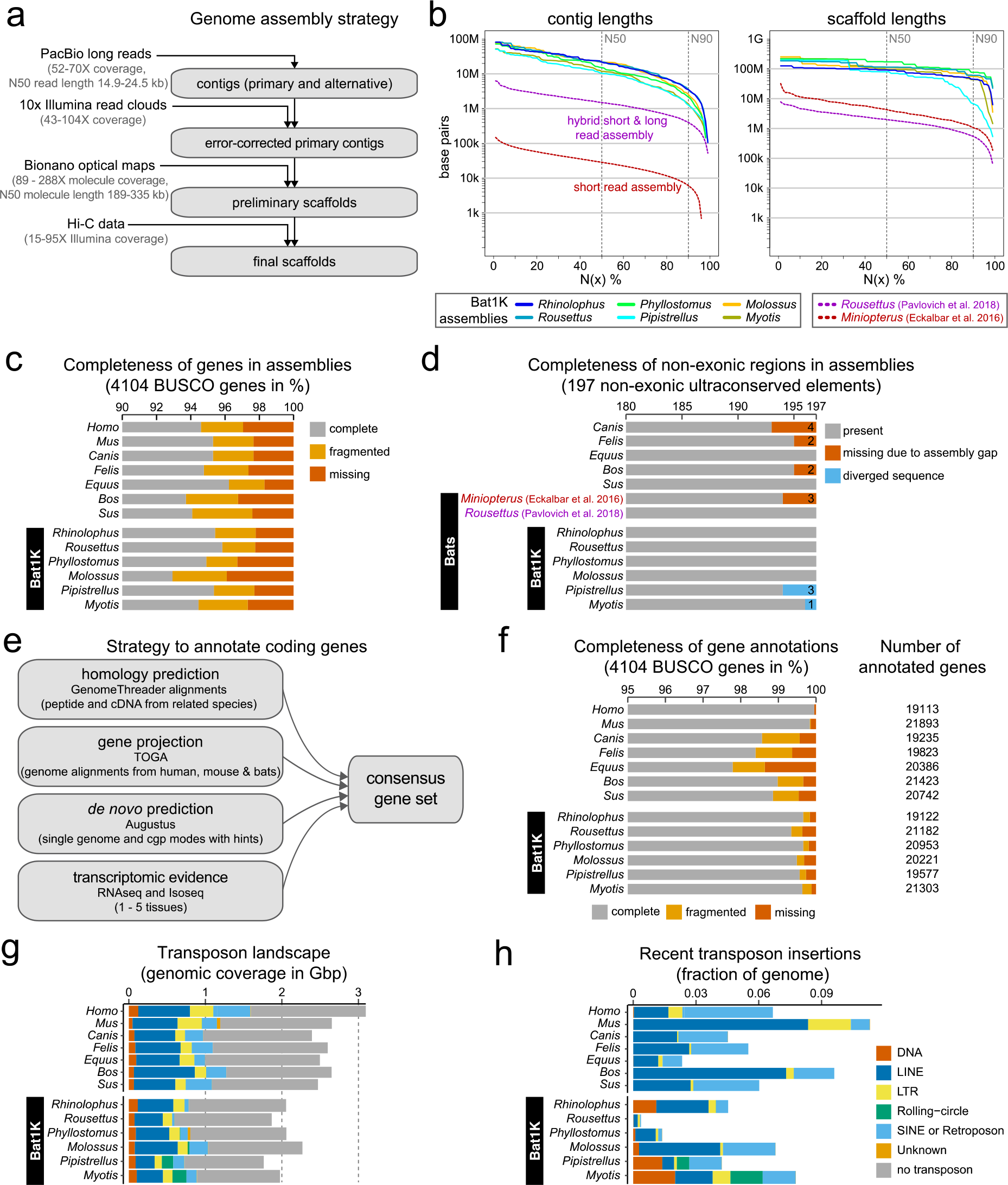
Assembly and annotation of the genomes of six bats. (a) Genome assembly strategy and the amount of data produced for assembling contigs and scaffolds. (b) Comparison of assembly contiguity. N(x)% graphs show the contig (left) and scaffold (right) sizes (y-axis), where x% of the assembly consists of contigs and scaffolds of at least that size. Dashed lines show contiguities of two recent bat assemblies, *Miniopterus* generated from short read data^21^, and *Rousettus* generated from a hybrid of short and long read data^22^. (c) Comparison of coding gene completeness. Bar charts show the percent of 4104 highly-conserved mammalian BUSCO genes that are completely present, fragmented or missing in the assembly. (d) Comparison of completeness in non-exonic regions. Bar charts show the number of detected ultraconserved elements that align at stringent parameters. Ultraconserved elements not detected are separated into those that are missing due to assembly incompleteness and those that exhibit real sequence divergence. Note that human and mouse are not shown here because both genomes were used to define ultraconserved elements^26^. (e) Our strategy to annotate coding genes combining various types of gene evidences. (f) Comparison of the completeness of gene annotations, using 4101 BUSCO genes, and the number of annotated genes. (f) Bar charts compare genome sizes and the proportion that consist of major transposon classes. (g) Fraction of the genome that consists of recent transposon insertions, defined as transposons that diverged less than 6.6% from their consensus sequence.

For all six bats, this sequencing and assembly strategy produced assemblies with contig N50 values ranging from 10.6 to 22.2 Mb (Fig. 1b, Table S2). Thus, our contigs are ≥355 times more contiguous than the recent *Miniopterus* assembly generated from short read data^21^, and ≥7 times more contiguous than a previous *Rousettus* assembly generated from a hybrid of short and long read data^22^ (Fig. 1b). Our scaffold N50 values ranged from 80.2 to 171.1 Mb and were often limited by the size of chromosomes (Fig. 1b, Table S2). We estimated that 87 to 99% of each assembly is in chromosome-level scaffolds (Table S3). Consensus base accuracies across the entire assembly range from QV 40.8 to 46.2 (Table S2) for the six bats (where QV 40 represents 1 error in 10,000 bp). Since the algorithms for assembling, scaffolding, and haplotyping are an active area of research^23^, we expect that in the future even more complete genome reconstructions can be produced with the data we collected. Even so, our current strategy and algorithms generated chromosome-level assemblies of the six bats with unprecedented contiguity, which are comparable to the best reference-quality genomes currently generated for any eukaryotic species with a complex, multi-gigabyte genome^24^. Importantly, they meet the Vertebrate Genome Project (VGP) minimum standard of 3.4.2QV40 and have been added to the VGP collection.

To assess genome completeness, we first evaluated the presence of 4,104 genes that are highly conserved among mammals (BUSCO, Benchmarking Universal Single-Copy Orthologs^25^). Between 92.9 and 95.8% of these genes were completely present in our assemblies, which is comparable to the assemblies of human, mouse, and other Laurasiatheria (Fig. 1c, Table S4). Second, to assess completeness in non-exonic genomic regions, we determined how many of 197 non-exonic ultraconserved elements (UCEs)^26^ align at ≥85% identity to the human sequence. As expected, the vast majority of UCEs were detected in all assemblies (Fig. 1d). Two to four UCEs were not detected in *Miniopterus*, dog, cat, and cow due to assembly incompleteness (i.e. assembly gaps; Table S5, Fig. S1). In the bat genomes reported herein, no UCEs were missing due to assembly incompleteness. Instead, one to three UCEs were not detected in our *Myotis* and *Pipistrellus* assemblies because the UCE sequences are more than 85% diverged (Table S5), a striking result given that UCE’s are highly conserved across other more divergent mammals (e.g. human-mouse-rat comparison). To determine if this sequence divergence was caused by base errors in the assemblies, we aligned raw PacBio and Illumina reads and sequencing data of related bats, which confirmed that these UCEs are truly diverged (Figs. S1-S5). In summary, our six bat assemblies are highly complete and revealed the first examples of highly diverged UCEs.

### Genome Annotation

To comprehensively annotate genes, we integrated a variety of evidence (Fig. 1e). First, we aligned protein and cDNA sequences of a related bat species to each of our six genomes (Table S6). Second, we projected genes annotated in human, mouse^27^ and two bat assemblies (*Myotis lucifugus* (Ensembl) and *Myotis myotis* (Bat1K)) to our genomes via whole-genome alignments^28^. Third, we generated *de novo* gene predictions by applying Augustus^29^ with a trained bat-specific gene model in single-genome mode to individual genomes, and in comparative mode to a multiple genome alignment including our bat assemblies. Fourth, we integrated transcriptomic data from both publicly available data sources and our own Illumina short read RNA-seq data (Table S7). Additionally, we generated PacBio long read RNA sequences (Iso-seq) from all six species to capture full-length isoforms and accurately annotate untranslated regions (UTRs) (Table S8). Iso-seq data were processed using the TAMA pipeline^30^ which allowed capturing a substantially greater diversity of transcripts and isoforms than the default pipeline (https://github.com/PacificBiosciences/IsoSeq3). All transcriptomic, homology-based and *ab initio* evidence were integrated into a consensus gene annotation that we further enriched for high-confidence transcript variants and filtered for strong coding potential.

For the six bats, we annotated between 19,122 and 21,303 coding genes (Fig. 1f). These annotations completely contain between 99.3 and 99.7% of the 4,104 highly conserved mammalian BUSCO genes (Fig. 1f, Table S4), showing that our six bat assemblies are highly complete in coding sequences. Since every annotated gene is by definition present in the assembly, one would expect that BUSCO applied to the protein sequences of annotated genes and BUSCO applied to the genome assembly should yield highly similar statistics. However, the latter finds only 92.9 to 95.8% of the exact same gene set as completely present, showing that BUSCO applied to an assembly only, underestimates the number of completely contained genes. Importantly, this gene annotation completeness of our bats is higher than the Ensembl gene annotations of dog, cat, horse, cow and pig, and is only surpassed by the gene annotations of human and mouse, which have received extensive manual curation of gene models (Fig. 1f, Table S4). This suggests reference-quality genome assemblies and the integration of various gene evidence as detailed above, can be used to generate high-quality and near-complete gene annotations of bats as well as other species too. All individual evidence and the final gene set can be visualized and obtained from the Bat1K genome browser (https://genome-public.pks.mpg.de).

### Genome Sizes and Transposable Elements

At ~2 Gb, bat genomes are generally smaller than genomes of other placental mammals that are typically between 2.5 and 3.5 Gb^2^. Nevertheless, our assemblies revealed noticeable genome size differences within bats, with assembly sizes ranging from 1.78 Gb for *Pipistrellus* to 2.32 Gb for *Molossus* (Fig. 1g). As genome size is often correlated with transposable element (TE) content and activity, we focused on the genomes of the six bats and seven other representative Boreoeutherian mammals (Laurasiatheria + Euarchontoglires), selected for the highest genome contiguity, and used a previously-described workflow and manual curation to annotate TEs^31^. This showed that TE content generally correlates with genome size (Fig. 1g). Next, we compared TE copies to their consensus sequence to obtain a relative age from each TE family. This revealed an extremely variable repertoire of TE families with evidence of recent accumulation (defined as consisting of insertions with divergences<6.6% from the relevant consensus sequence). For example, while the 1.89 Gb *Rousettus* genome exhibits few recent TE accumulations, ~0.38%, while ~4.2% of the similarly sized 1.78 Gb *Pipistrellus* genome is derived from recent TE insertions (Fig. 1g-h). The types of TE that underwent recent expansions also differ substantially in bats compared to other mammals, particularly in regards to the evidence of recent accumulation by rolling-circle and DNA transposons in the vespertilionid bats (Fig. 1g-h). These two TE classes have been largely dormant in most mammals for the past ~40 million years and recent insertions are essentially absent from other Boreoeutherian genomes^32^. These results add to previous findings revealing a substantial diversity in TE content within bats, with some species exhibiting recent and ongoing accumulation from TE classes that are extinct in most other mammals while other species show negligible evidence of TE activity^33^.

### The Origin of Chiroptera within Laurasiatheria

Identifying the evolutionary origin of bats within Laurasiatheria is a key prerequisite for comparative analyses aimed at revealing the genomic basis of traits shared by bats. However, the phylogeny of Laurasiatheria and, in particular, the position of bats has been a long-standing, unresolved phylogenetic question^14,34^. This is perhaps the most challenging interordinal problem in placental mammal phylogenetics, as multiple phylogenetic and systematic investigations using large nucleotide and genomic scale datasets or transposable element insertions support alternative topologies^35^. These incongruent results have been attributed to the challenge of identifying two, presumably short, internal branches linking four clades (Chiroptera, Cetartiodactyla, Perissodactyla, Carnivora + Pholidota) that diverged in the Late Cretaceous^35^.

We revisited this question leveraging the high completeness of our gene annotation. First, we extracted a comprehensive set of 12,931 orthologous protein-coding genes from the 48 mammalian genomes, resulting in a dataset comprising 21,471,921 aligned nucleotides in length, which contained 7,913,054 parsimony-informative sites. The best-fit model of sequence evolution for each of the 12,931 gene alignments was inferred using ModelFinder^36^ (Table S9). The species tree was then estimated by maximum likelihood using the model-partitioned dataset with IQTREE^37^ and rooted on Atlantogenata^38^. Branch-support values were obtained by UFBoot^39^ with 1000 bootstrap pseudoreplicates. These analyses led to 100% bootstrap support across the entire tree (Fig. 2a) and seemingly identified the origin of bats within Laurasiatheria. The basal split is between Eulipotyphla and other laurasiatherians (i.e., Scrotifera). Within Scrotifera, Chiroptera is the sister clade to Fereuungulata (Cetartiodactyla + Perissodactyla + Carnivora + Pholidota). This tree disagrees with the Pegasoferae hypothesis^40^, which groups bats with Perissodactyla, Carnivora and Pholidota, but agrees with concatenation analyses of phylogenomic data^41^. Evolutionary studies based on 102 retroposons, including ILS-aware analyses, also support a sister-group relationship between Chiroptera and Fereuungulata, but differ from the present analyses in supporting a sister-group relationship between Carnivora and Cetartiodactyla^34,35^. However, as the number of homologous sites increases in phylogenomic datasets, so too does bootstrap support^42^, even sometimes for an incorrect phylogeny^43^, and as non-coding sequences can produce a different topology than coding sequences ^44^, we further explored the phylogenomic signal within our genomes.

**Figure 2:**
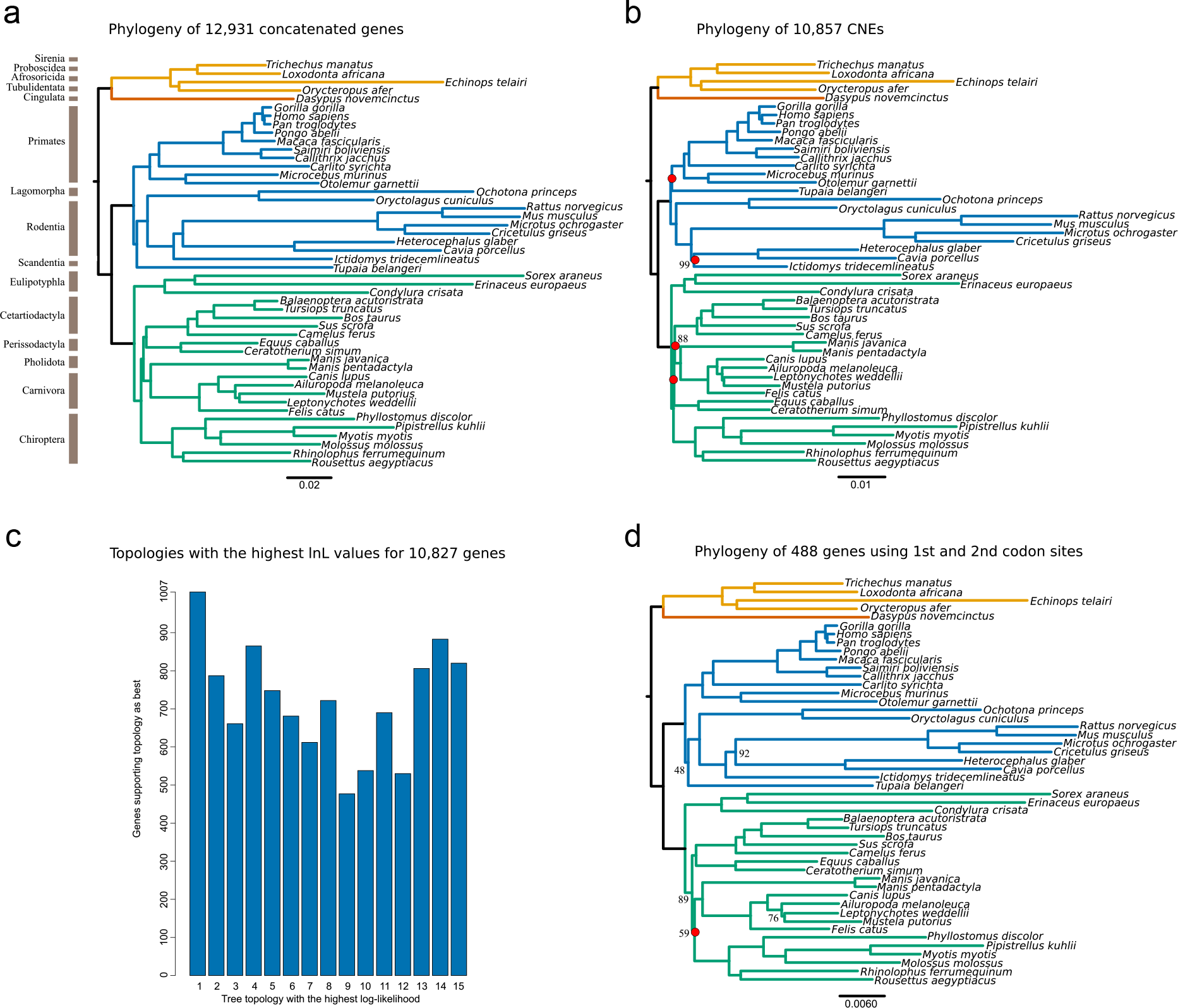
Phylogenetic analysis of Laurasiatheria. (a) We inferred a mammalian phylogram using a supermatrix of 12,931 concatenated genes and the maximum likelihood method of tree reconstruction (topology 1, Fig. S6). (b) A total of 10,857 conserved non-coding elements (CNEs) were used to determine a mammalian phylogeny using non-coding regions (topology 2, Fig. S6). Bootstrap support values less than 100 are displayed, with internal nodes that differ to the protein-coding supermatrix highlighted in red. (c) All gene alignments were fit to the 15 laurasiatherian topologies (Fig. S6) explored to determine which tree had the highest likelihood score for each gene. The number of genes supporting each topology are displayed. (d) A supermatrix consisting of 1^st^ and 2^nd^ codon sites from 448 genes that are evolving under homogenous conditions, thus considered optimal ‘fit’ for phylogenetic analysis, was used to infer a phylogeny using maximum likelihood (topology13 Feig. S6). Bootstrap support values less than 100 are displayed, with internal nodes that differ to the protein-coding supermatrix phylogeny highlighted in red.

To assess whether the tree inferred from the concatenated dataset (Fig. 2a) is also supported by the non-coding part of the genome, we estimated a phylogeny using the models of best fit (Table S9) for a dataset comprising 10,857 orthologous conserved non-coding elements (CNEs), which contained 5,234,049 nucleotides and 1,225,098 parsimony-informative sites (Table S10), using methods as described above. The result of this analysis (Fig. 2b) supports a tree similar but not identical to that inferred from the protein-coding sequences (Fig. 2a), including a sister-group relationship between Chiroptera and Fereuungulata, but with Perissodactyla more closely related to Carnivora + Pholidota than to Cetartiodactyla. The CNE tree also recovered a different position for *Tupaia* (Scandentia) within Euarchontoglires.

Given that two very short branches at the base of Scrotifera define relationships between its four major clades (Carnivora + Pholidota, Cetartiodactyla, Chiroptera, Perissodactyla), this region of the placental tree may be in the “anomaly zone”, defined as a region of tree space where the most common gene tree(s) differs from the species tree topology^45^. In the case of four taxa and a rooted pectinate species tree, anomalous gene trees should be symmetric rather than pectinate. To explore this, we estimated the maximum-likelihood support of each protein-coding gene (n=12,931) for the 15 possible bifurcating topologies involving four clades, in our case with Eulipotyphla as the outgroup (Fig. S6), and with the sub-trees for the relevant clades identical to those in Fig. 2a. Based on the log-likelihood scores, 2,104 gene alignments supported more than one tree, so these genes were excluded from further analysis. The remaining 10,827 genes supported one fixed tree topology over the other 14 (Table S11), with the number of genes supporting each topology highlighted in Fig. 2c. The best-supported topology was that of our concatenated dataset for protein-coding genes (Fig. 2a; Tree1 with 1007/10827 genes), showing a sister group relationship between Chiroptera and Fereuungulata, which is also supported by the CNEs (Fig. 2b). This suggests that the majority of the genome supports a sister relationship between Chiroptera and the other Scrotifera. That said, there were four other topologies that had support from >800 genes (Tree14 883/10827; Tree04 865/10827; Tree15 820/10827; Tree13 806/10827) (Fig. 2c). However, even with similar support levels for several topologies, the phylogenetic position for Chiroptera is pectinate on the most common gene tree and does not qualify as anomalous. If the base of Scrotifera is in the anomaly zone, as suggested by coalescence analyses of retroposon insertions^35^, then we may expect the most common gene tree(s) to be symmetric rather than pectinate. We may also expect the species tree based on concatenation to be symmetric instead of pectinate^45^. One explanation for the absence of anomalous gene trees, and for a pectinate species tree based on concatenation, is that both protein-coding genes and CNEs are generally under purifying selection, which reduces both coalescence times and incomplete lineage sorting relative to neutrally evolving loci^46,47^.

Bias in phylogenetic estimates can also be due to model misspecification, which is an inadequate fit between phylogenetic data and the model of sequence evolution used^48^. Misleading support for incorrect phylogenies can also be due to gene tree error arising from a lack of phylogenetic informativeness amongst data partitions^49^. To overcome these biases, we performed a series of compatibility analyses on each gene partition and across the supermatrix at 1^st^, 2^nd^ and 3^rd^ codon sites; 1^st^ + 2^nd^ codon sites; 1^st^ + 2^nd^ + 3^rd^ codon sites; amino acids, assuming a 4-state alphabet for nucleotides and a 20 state-alphabet for amino acids (see supplementary methods section 4.2). We excluded all alignments for which evidence of saturation of substitutions and thus decay of the historical signal was detected by SatuRation 1.0 (https://github.com/lsjermiin/SatuRationSatuRation). Furthermore, we excluded all alignments for which model mis-specification due to evolution under non-homogeneous conditions was detected by the matched-pairs test of symmetry^50^ implemented in Homo 2.0 (https://github.com/lsjermiin/Homo2.0).

A total of 488 gene alignments, consisting of 1st and 2nd codon positions and containing all 48 taxa, were considered optimal for phylogenetic analysis (Table S12). We concatenated these data into a supermatrix of 241,098 nucleotides in length with 37,588 informative positions and completed all phylogenetic analyses using methods as described above. However, this reduced data set did not provide an unambiguous phylogenetic estimate. Specifically, while the best-supported topology differed from the best trees inferred using all protein-coding genes and CNEs in its position of Chiroptera, which is now sister to Carnivora + Pholidota (Fig. 2d), this node has low bootstrap support (58%; topology 13; Fig. 2d) and Approximately Unbiased (AU) tests could not reject the topologies depicted in Fig. 2a and 2b. Furthermore, the phylogeny inferred from the subset of 488 genes is also symmetric for the four major lineages of Scrotifera, as may be expected if this node is in the anomaly zone and concatenation is misleading. We further analysed these data using a single-site coalescence-based method, SVDquartets^51,52^, which provides an alternative to concatenation. The resulting optimal topology also supported Chiroptera as sister taxa to Fereuungulata (Fig. S7, topology 1), which is the most supported position from all of our analyses and data partitions.

Taken together, multiple lines of evidence suggest that the majority of the genome supports Chiroptera as sister to all other scrotiferans. However, different regions of the genome can and do reflect alternative evolutionary scenarios. This highlights the importance of generating phylogenetic inferences from multiple genomic regions and the importance of screening these regions for violations of phylogenetic assumptions and incongruent signals, especially when dealing with short internal branches.

### Genome-wide screens for gene selection, losses and gains

To study the genomic basis of exceptional traits shared by bats, we first performed three unbiased genome-wide screens for gene changes that occurred in the six bats. First, we screened 12,931 genes classified as 1:1 orthologs for signatures of positive selection on the ancestral bat (stem Chiroptera) branch under the aBSREL model^53^ using HyPhy^54^ and the best-supported phylogeny (Fig. 2a). For genes with significant evidence for selection after multiple test correction (FDR<0.05), we manually inspected the underlying alignment to ensure homology (supplementary methods section 4.3.1), and additionally required that the branch-site test implemented in PAML codeml^55^ independently verified positive selection (P<0.05). This revealed 9 genes with a robust signal of positive selection at the bat ancestor (Table S13). While these 9 genes have diverse functions, they included two genes with hearing-related functions, which may relate to the evolution of echolocation. These genes, *LRP2* (low-density lipoprotein receptor-related protein 2, also called megalin) and *SERPINB6* (serpin family B member 6) are expressed in the cochlea and associated with human disorders involving deafness. *LRP2* encodes a multi-ligand receptor involved in endocytosis that is expressed in the kidney, forebrain and, importantly, is also expressed in the cochlear duct^56^. Mutations in this gene are associated with Donnai-Barrow Syndrome, an autosomal recessive disease with symptoms including sensorineural deafness^57^, and progressive hearing loss has also been observed in *Lrp2* knockout mice^58^. Similarly, *SERPINB6* is associated with non-syndromic hearing loss and this serine protease inhibitor is expressed in cochlear hair cells^59,60^. Sites identified as having experienced positive selection at the bat ancestor showed bat-specific substitutions in both genes. Interestingly, the echolocating bats showed a specific asparagine to methionine substitution in *LRP2*. In *Rousettus,* the only non-laryngeal echolocator in our six bats, this site has been substituted for a threonine. Combined with analysis of 6 other publicly available bat genomes (n=6), we confirmed the presence of a methionine in all laryngeal echolocating bats (n=9) and a threonine residue in all non-echolocating pteropodids (n=3) (Fig. S8).

We also initially identified positive selection in the bat ancestor in a third hearing-related gene, *TJP2* (tight junction protein 2), that is expressed in cochlear hair cells and associated with hearing loss^61,62^. However, manual inspection revealed a putative alignment ambiguity and the manually-corrected alignment had a reduced significance (aBSREL raw P=0.009, not significant after multiple test correction considering 12,931 genes). Interestingly, the corrected alignment revealed a four amino acid microduplication found only in echolocating bats (n=9) (Fig. S9), which may be explained by incomplete lineage sorting or convergence. It should be noted that insertions and deletions may also affect protein function but are not considered by tests for positive selections, however a phylogenetic interpretation of these events may uncover functional adaptations. In general, experimental studies are required to test whether the pattern of positive selection and bat-specific mutations on the stem Chiroptera branch affect hearing-related functions of these three genes. If so, this would provide molecular support for laryngeal echolocation as a shared ancestral trait of bats and subsequent loss in pteropodids, informing a long-standing debate in bat biology of whether ancestral bats had the ability to echolocate^12^.

In addition to hearing-related genes, our genome-wide screen revealed selection on immunity-related genes, *CXCL13* (C-X-C motif chemokine ligand 13), *NPSR1* (neuropeptide S receptor 1) and *INAVA* (innate immunity activator), which may underlie bats’ unique tolerance of pathogens^9^. The CXCL13 (previously B-lymphocyte chemoattractant) protein is a B-cell specific chemokine, which attracts B-cells to secondary lymphoid organs, such as lymph nodes and spleen^63^. *NPSR1* expresses a receptor activated by neuropeptide S. Activation of NSPR1 induces an inflammatory immune response in macrophages and *NSPR1* polymorphisms have been associated with asthma in humans^64^. *INAVA* encodes an immunity-related protein with a dual role in innate immunity. In intestinal epithelial cells, this gene is required for intestinal barrier integrity and the repair of epithelial junctions after injury^65,66^. Consistent with these functions, mutations in human *INAVA* are associated with inflammatory bowel disease^67^, a disorder characterized by chronic inflammation of the gastrointestinal tract and an increased susceptibility to microbial pathogens. In macrophages, *INAVA* amplifies an IL-1β-induced pro-inflammatory response by enhancing NF-kB signalling^66^.

While a genome-wide screen for significant signatures of positive selection is comprehensive, considering 12,931 orthologous genes may reduce sensitivity due to the necessity to correct for 12,931 statistical tests. To increase the sensitivity in detecting positive selection in genes relevant for prominent bat traits (i.e. longevity, immunity, metabolism^2^) we further performed a screen considering 2,453 candidate genes (Table S14) associated with these terms according to Gene Ontology (GO), AmiGO^68^ and GenAge^69^ annotations. This reduced gene set permitted a screen for signatures of positive selection using both the aBSREL model and the branch-site test implemented in codeml (supplementary methods section 4.3.1). Requiring significance by both aBSREL and codeml (FDR<0.05), we found 10 additional genes with robust evidence of positive selection in the ancestral bat lineage (Table S15, Fig. S10). These genes include *IL17D* and *IL-1β*, which are involved in immune system regulation ^70^ and NF-kB activation (*IL-1β*)^66,71^, and *GP2* and *LCN2*, which are involved in the response to pathogens^72,73^. Interestingly, selection was also inferred for *PURB*, a gene that plays a role in cell proliferation and regulates the oncogene *MYC*^74^, which was previously shown to be under divergent selection in bats^16^ and which exhibits a unique anti-ageing transcriptomic profile in long lived *Myotis* bats^8^. Overall, combining genome-wide and candidate gene screens revealed robust patterns of selection in stem Chiroptera on several genes involved in immunity and infection, which suggests that ancestral bats evolved immunomodulatory mechanisms that enabled a higher tolerance to pathogens.

Second, we used a previously developed approach^75^ to systematically screen for gene loss. This revealed 10 genes that are inactivated in our six bats but present in the majority of Laurasiatheria (Table S16). Two of these genes again point to changes in immune function in bats, having immune-stimulating and pro-inflammatory functions; *LRRC70* (leucine rich repeat containing 70, also called synleurin) and *IL36G* (interleukin 36 gamma) (Fig. 3a). *LRRC70* is expressed in a broad range of tissues and potentiates cellular responses to multiple cytokines^76^ and is well conserved among Laurasiatheria. Importantly, *LRRC70* strongly amplifies bacterial lipopolysaccharide-mediated NF-kB activation^76^. Our finding of *LRRC70* loss in bats makes this poorly characterized gene an interesting target for future mechanistic studies. *IL36G*, encodes a pro-inflammatory interleukin belonging to the interleukin-1 family. Increased expression of *IL36G* was detected in psoriasis and inflammatory bowel disease patients, and *IL36G* is likely involved in the pathophysiology of these diseases by inducing the canonical NF-kB pathway and other proinflammatory cytokines^77–79^. Further analysis of common mutations between our assembled genomes and previously published bat genomes (n=9), revealed these genes were in fact lost multiple times within Chiroptera (Fig. S11 and S12), suggesting these genes came under relaxed selection in bats followed by with subsequent gene losses. Together, genome-wide screens for gene loss and positive selection revealed several genes involved in NF-kB signalling (Fig. 3b), suggesting that altered NF-kB signalling may contribute to immune related adaptations in bats.

**Figure 3:**
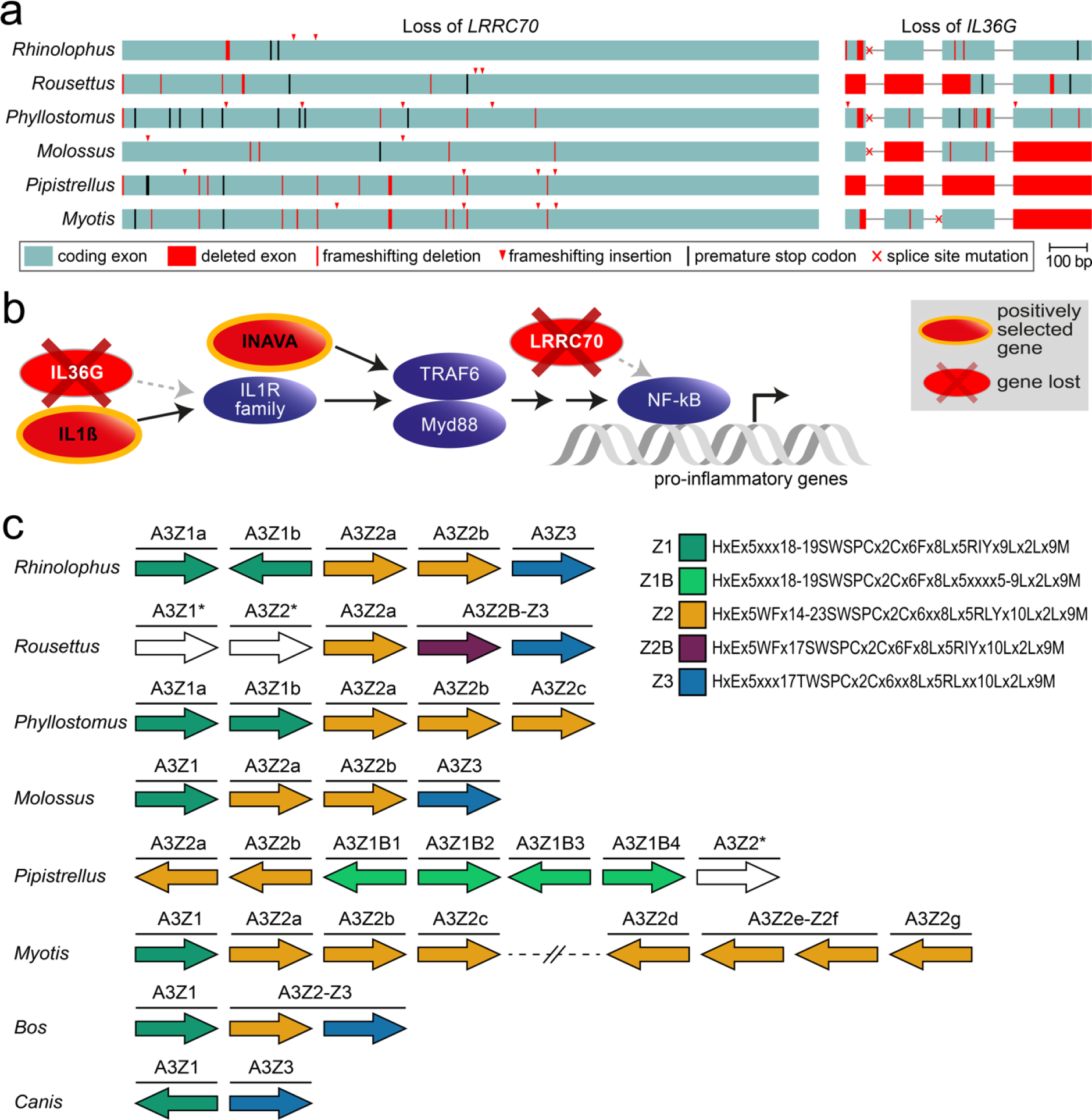
Genome-wide screens highlight changes in genes potentially involved in bat’s unique immunity. (a) Inactivation of the immune genes *LRRC70* and *IL36G*. Boxes represent coding exons proportional to their size, overlaid with gene-inactivating mutations present in the six bats. (b) Diagram showing the canonical NF-kB signalling pathway (purple) and interacting proteins which have experienced positive selection or have been lost in bats. (c) Expansion of the *APOBEC3* gene locus in bats. Each arrow represents a cytidine deaminase domain, coloured by domain subtypes as defined by given motifs, with likely pseudogenes are in white. Genes containing multiple deaminase domains are shown as a single bar over more than one domain. A transposition event in *Myotis* has created two *APOBEC3* loci on different chromosomes, indicated by the broken line in this species. Cow and dog are shown as two Laurasiatheria outgroups, where cow also represents the likely, mammalian ancestral state.

Third, we investigated changes in gene family size, which revealed 35 cases of significant gene family expansions and contractions at the bat ancestor (Table S17). Among these, we inferred an expansion of the *APOBEC* gene family. Expansion involved *APOBEC3*-type genes (Fig. 3c) and supported a small expansion in the ancestral bat lineage, followed by up to 14 duplication events within Chiroptera. The *APOBEC3* locus is highly-dynamic, with a complex history of duplication, loss and fusion in Mammalia^80^. Our analysis of this locus in Chiroptera adds to previous evidence of a genus specific expansion in the flying foxes (genus *Pteropus*)^81^, showing this locus has undergone many independent expansions in bats. *APOBEC* genes are DNA and RNA editing enzymes with roles in lipoprotein regulation and somatic hypermutation^82^. *APOBEC3*-type genes have been previously associated with restricting viral infection, transposon activity^82^ and may also be stimulated by interferon signalling^83^. Expansion of *APOBEC3* genes in multiple bat lineages suggests these duplications may contribute to viral tolerance in these lineages.

### Integrated Viruses in Bat Genomes

There is mounting evidence to suggest that bats are major zoonotic reservoir hosts, as they can tolerate and survive viral infections (e.g. Ebola and MERs), potentially due to adaptations in their immune response^84^, consistent with our findings of selection in immune-related genes (e.g. *INAVA*) and expansions of the viral-restricting *APOBEC3* gene cluster. We screened our high-quality genomes to ascertain the number and diversity of endogenous viral elements (EVEs), considered as ‘molecular fossil’ evidence of ancient infections. Given their retroviral life cycle endogenous retroviruses (ERVs) are the largest group found among all EVEs in vertebrate genomes^85,86^ (making up ~10% of the mouse^87^ and 8% of the human genome^88^), while non-retroviral EVEs are far less numerous in animal genomes^86^.

Using reciprocal BLAST searches and a custom comprehensive library of viral protein sequences we first screened our six bat genomes and seven mammalian outgroups (supplementary methods section 3.4) for the presence of EVEs, including ERVs and non-retroviral EVEs. We identified three predominant non-retroviral EVE families: *Parvoviridae*, *Adenoviridae* and *Bornaviridae* (Fig. 4a). Parvovirus and bornavirus integrations were found in all bats except for *Rousettus* and *M. molossus* respectively. A partial filovirus EVE was found to be present in the Vespertilionidae (*Pipistrellus & Myotis*), but absent in the other bat species, suggesting that vespertilionid bats have been exposed in the past to and can survive filoviral infections, corroborating a previous study^89^.

**Figure 4:**
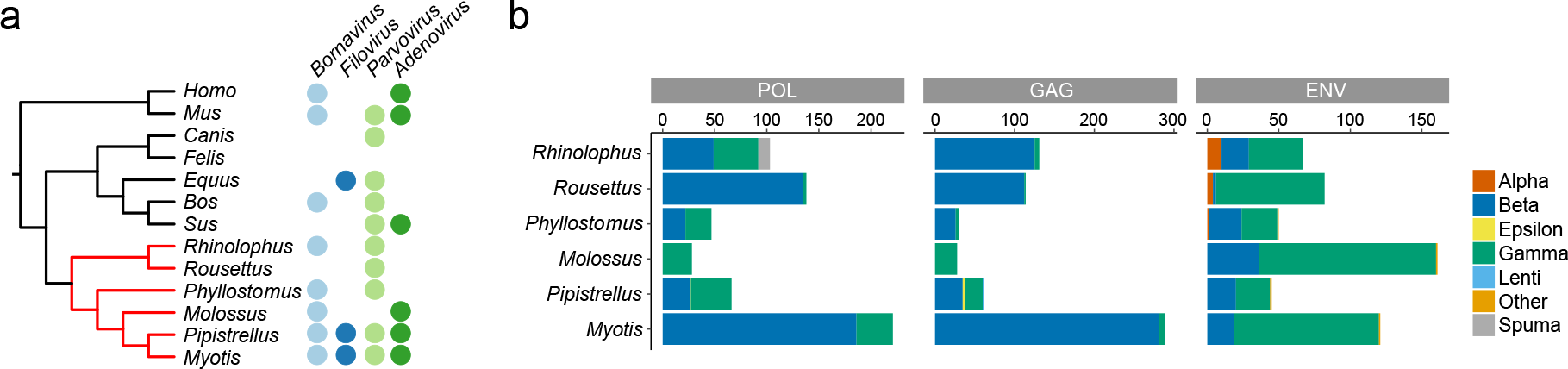
Endogenous Viruses in Bat Genomes. (a) Viral families identified in more than one genus mapped to phylogenetic tree of six bat species and seven additional mammals. Endogenous sequences identified as *Adenoviridae*, *Parvoviridae*, *Filoviridae* and *Bornaviridae* were represented across several mammalian genera. (b) Bar plot showing numbers of sequences found for each of the viral proteins in six species of bat and the representation in all seven *Retroviridae* genera.

Next, we identified retroviral proteins from all ERV classes within the bat genomes. Consistent with other mammals, the highest number of integrations came from beta- and gamma-like retroviruses^90,91^, with beta-like integrations most common for *pol* and *gag* proteins and gamma-like integrations most common for *env* proteins in most of the bats (Fig. 4b & Fig. S13). Overall, the highest number of integrations was observed in *M. myotis* (n=630), followed by *Rousettus* (n=334) with *Phyllostomus* containing the lowest (n=126; Fig. 4b, Table S18). Additionally, we detected ERV sequences with hits for alpha- and lenti-retroviruses in reciprocal BLAST searches. Until now, alpharetroviruses were considered as exclusively endogenous avian viruses^92^. Thus, our discovery of endogenous alpharetroviral-like elements in bats is the first record of these sequences in mammalian genomes, widening the known biodiversity of potential hosts for retrovirus transmission. We detected several alpha-like *env* regions in *Phyllostomus, Rhinolophus*, and *Rousettus* (Fig. 4b), showing that multiple and diverse bat species have been and possibly are being infected by alpharetroviruses. We also detected lentivirus *gag*-like fragments in *Pipistrellus*, which are rarely observed in endogenized form^93^.

To identify historical ancestral transmission events, we reconstructed a phylogenetic tree from our recovered ERVs with the known viral protein ‘probe’ sequences for all six bat genomes and seven mammalian outgroups (Fig. S14). The majority of sequences group as single bat-species clusters, suggesting that relatively recent integration events, more than ancestral transmission (Fig. S14) govern the ERV diversity. While, most ERVs are simple retroviruses, consisting of *gag*, *pol* and *env* genes, we found an unusual diversity of complex retroviruses in bats, which are generally rare in endogenous form^93–95^ (Fig. S14). We detected a clade of 5 *Rhinolophus pol* sequences clustered together with reference foamy retroviruses – Feline Foamy Virus (FFV) and Bovine Foamy Virus (BFV). Foamy retroviruses in bats were detected before from metagenomic data from *Rhinolophus affinis*^96^, however, until now it was unclear whether these sequences represented exogenous or endogenous viruses^97^. With the detection of these sequences, we can now confirm the presence of endogenous spumaretroviruses in the *R. ferrumequinum* genome, which furthers our understanding of the historical transmission dynamics of this pathogen. We also detected *pol* sequences in the *Molossus* genome clustering closely with reference delta sequences (Bovine Leukemia virus – BLV, Human T-lymphotropic Virus – HTLV). *Pol* regions for delta retroviruses in bats have not been detected before, with only partial gag and a single LTR identified previously in *Miniopterus* and *Rhinolophus* species^94,98^.

Overall these results show that bat genomes contain a surprising diversity of ERVs, with some sequences never previously recorded in mammalian genomes, confirming interactions between bats and complex retroviruses, which endogenize exceptionally rarely. These integrations are indicative of past viral infections, highlighting which viruses bat species have co-evolved with and tolerated, and thus, can help us better predict potential zoonotic spillover events and direct routine viral monitoring in key species and populations. In addition, bats, as one of the largest orders of mammals, are an excellent model to observe how co-evolution with viruses can shape the mammalian genome over evolutionary timescales. For example, the expansion of the *APOBEC3* genes in bats shown herein, could be a result of a co-evolutionary arms race shaped by ancient retroviral invasions, and could contribute to the restriction in copy number of endogenous viruses in some bat species. Given that these findings were generated from only six bat genomes we can be confident that further cross-species comparison with similar quality bat genomes will bring even greater insight.

### Changes in Non-Coding RNAs

In addition to coding genes, changes in non-coding (nc)RNAs can be associated with interspecific phenotypic variation and can drive adaptation^99,100^. We used our reference-quality genomes to comprehensively annotate non-coding RNAs and search for ncRNA changes between bat species and other mammals. To annotate different classes of conserved non-coding RNA genes, we used computational methods that capture characteristic sequence and structure features of ncRNAs (Fig. 5a; supplementary methods section 5.1). We found that a large proportion of non-coding RNA genes were shared across all six bats (Fig. S15), and between bats and other mammals (e.g. 95.8% ~ 97.4% shared between bats and human).

**Figure 5:**
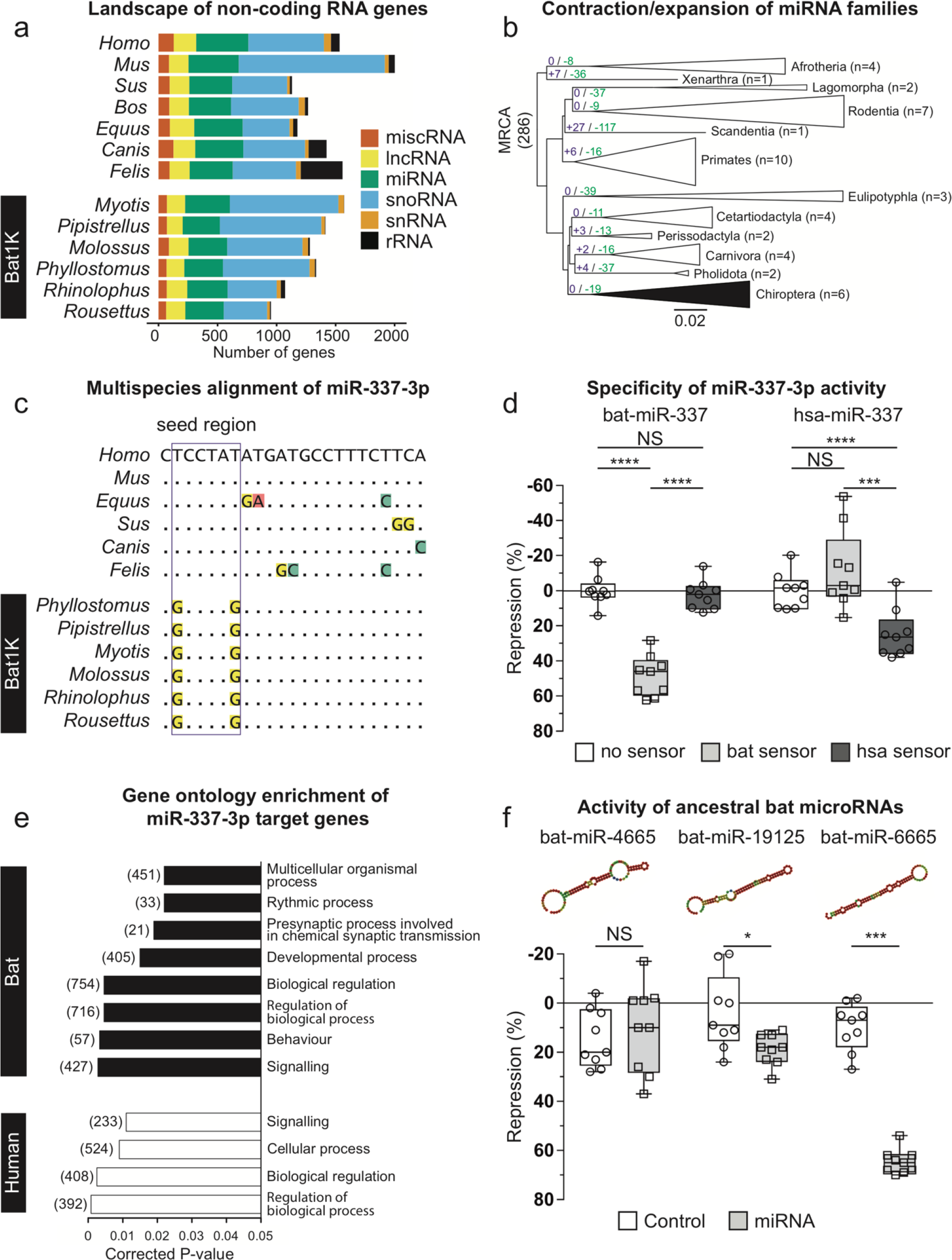
The evolution of non-coding RNA genes in bats. (a) The number of non-coding RNA genes annotated in six bat genomes and 7 reference mammalian genomes. (b) miRNA family expansion and contraction analyses in 48 mammalian genomes. The numbers highlighted on the branches designate the number of miRNA families expanded (purple, +) and contracted (green, −) at the order level. n indicates the number of species in each order used in the analysis. (c) The alignment of the mature miR-337-3p sequences across six bats and six reference species (miR-337-3p could not be found in *Bos taurus* genome). The box indicates the seed region of mature miR-337-3p, which is conserved across mammals, but divergent in bats. (d) Specificity of human (hsa) and bat miR-337-3p activity was shown using species specific sensors in luciferase reporter assays (n=9 per experiment; see supplementary methods section 5.4). Significance was calculated using two-way ANOVA test, followed by post-hoc Tukey calculation. Statistical significance is indicated as: ***p<0.001; ****p<0.0001. (e) Gene ontology enrichment (via DAVID) of targets predicted for human and bat miR-337-3p (f) Validation of the activity of ancestral bat miRNAs, absent in the other mammalian genomes. The predicted secondary structures for each novel miRNA are displayed. For each miRNA, the sensor was tested against a control unrelated miRNA that was not predicted to bind to the sensor (left) and the cognate miRNA (right) in luciferase reporter assays (n=9 per experiment; see supplementary methods section 5.4). Significance for each independent control-miRNA pair was calculated using pairwise t tests. Statistical significance is indicated as: *p<0.05; ***p<0.001. Box plots extend from the 25th to 75th percentiles, the central line represents the median value, and whiskers are drawn using the function “min to max” in GraphPad Prism7 (GraphPad Software, La Jolla California USA, http://www.graphpad.com) and go down to the smallest value and up to the largest.

Within ncRNAs, we next investigated microRNAs (miRNA), which can serve as developmental and evolutionary drivers of change^101^. We employed a strict pipeline to annotate known miRNAs in our six bat genomes and in the 42 outgroup mammal taxa (Table S19, supplementary methods section 5.1) and investigated how the size of miRNA families evolved using CAFÉ^102^. We identified 286 miRNA families present in at least one mammal and observed massive contractions of these miRNA families (Fig. S16) with an estimated overall rate of ‘death’ 1.43 times faster than the rate of ‘birth’ (see supplementary methods section 5.1). There were 19 families that significantly (FDR<0.05) contracted in the ancestral bat branch, with no evidence of expansions, and between 4 and 35 miRNA families were contracted across bats (Fig. 5b, Fig. S16). We also inferred the miRNA families lost in each bat lineage using a Dollo parsimony approach, which revealed 16 miRNA families that were lost in the bat ancestor (Fig. S17 and S18). Interestingly, the oncogenic miR-374 was lost in all bat species but was found in the other examined orders (Table S19). Since miR-374 promotes tumour progression and metastasis in diverse human cancers^103^, this bat specific loss may contribute to low cancer rates in bats^16^.

Next, we investigated the evolution of single-copy miRNA genes to determine if sequence variation in these miRNAs may be driving biological change. Alignments of 98 highly conserved, single-copy miRNA genes identified across all 48 mammal genomes revealed that one miRNA, miR-337-3p, had unique variation in the seed region in bats compared to all other 42 mammals (Fig. 5c). miR-337-3p was pervasively expressed in brain, liver, and kidney across all six bat species (Fig. S19). Given that the seed sequences of microRNAs represent the strongest determinant of target specificity, these changes are expected to alter the repertoire of sequences targeted by miR-337-3p in bats.

To test this hypothesis, we used reporter assays^104,105^ to determine if the bat and human versions of miR-337-3p were functionally active and if they showed species-specific regulation of an “ideal” predicted target sequence (Table S20). While bat miR-337-3p strongly repressed the expression of its cognate bat target sequence, it had no effect on the human site, and *vice versa* (Fig. 5d). This result demonstrated that the miR-337-3p seed changes found in bats alter its binding specificity. To explore whether this difference in binding specificity changes the set of target genes regulated by bat miR-337-3p, we used our raw Iso-seq data to identify 3’UTRs of coding genes in bats (n=6,891-16,115) and determined possible target genes of miR-337-3p using a custom *in silico* pipeline (Table S21; supplementary methods section 5.3.5). We also obtained the equivalent human 3’UTRs for all predicted bat 3’UTRs and identified the human miR-337-3p gene targets (supplementary methods section 5.3.5). In bats, miR-337-3p was predicted to regulate a distinct spectrum of gene targets compared to humans (Table S22). GO enrichment analysis of these target gene sets suggests a shift towards regulation of developmental, rhythmic, synaptic and behavioural gene pathways by miR-337-3p in bats (Fig. 5e), pointing to a dramatic change in processes regulated by miR-337-3p in bats compared to other mammals.

In addition to losses and changes in miRNAs, continuous miRNA innovation is observed in eukaryotes, which is suggested as a key player in the emergence of increasing organismal complexity^99^. To identify any novel miRNAs that evolved in bats, we performed deep sequencing of small RNA libraries from brain, liver and kidney for all six bats (Table S23), analysed these data using a comprehensive custom analysis pipeline (see supplementary methods section 5.3.3), and identified those miRNAs that possess seed regions not found in miRBase (release 22). This screen revealed between 122 and 261 novel miRNAs across the six bat genomes. Only a small number of these novel miRNAs were shared across the six bats, supporting rapid birth of miRNAs on bat lineages (Fig. S20). We identified 12 novel miRNAs that were found in all six bats but did not have apparent homologs in other mammals (Table S24). Prediction of miRNAs from genomic sequences alone may result in false positives due to the occurrence of short hairpin-forming sequences that are predicted to form hairpins but are not processed or functionally active, emphasizing the need for experimental testing of these miRNAs. Therefore, to test whether these candidates indeed function as miRNAs we selected the top 3 candidates (bat-miR-4665, bat-miR-19125, bat-miR-6665) (Table S24) based on their expression and secondary structures, and experimentally tested their ability to regulate an ideal target sequence in reporter assays, as above (Table S20). Two of the three miRNAs (miR-19125 and miR-6665) were able to regulate their targets, showing that they are actively processed by the endogenous miRNA machinery, and able to be loaded onto the RISC complex to repress target mRNAs (Fig. 5f). Thus, miR-19125 and miR-6665 represent true miRNAs that are novel to bats. Taken together, these data demonstrate innovation in the bat lineage with regard to miRNAs both in seed sequence variation as well as novel miRNA emergence.

In summary, our genomic screens and experiments revealed losses of ancestral miRNAs, gains of novel functional miRNA and a striking case of miRNA seed change that alters the target specificity. Changes in these miRNAs and their target genes point to a regulatory role in cancer, development and behaviour in bats. Further detailed mechanistic studies will be crucial to determine the role of these miRNAs in bat physiology and evolution.

## Discussion

We have used a combination of state-of-the-art methods including long-read, short-read, and scaffolding technologies to generate chromosome level, near-complete assemblies of six bats that represent diversity within Chiroptera. These reference-quality genomes improve on all published bat genomes and are on par with the best reference-quality genomes currently generated for any eukaryotic species with a complex, multi-gigabyte genome. Compared to the contiguity and completeness of previous bat genomes assembled with short reads, our reference-quality genomes offer significant advances. First, while fragmented and incomplete assemblies hamper gene annotation, reference-quality genomes allow comprehensive annotations by integrating a variety of methods and evidence. In particular, reference-quality genomes facilitate genome alignment, which provides a powerful way of transferring gene annotations of related species to new assemblies and ensures that transcriptomic data can be comprehensively mapped. Second, while fragmented and incomplete assemblies resulted in countless efforts by individual labs to laboriously clone and re-sequence genomic loci containing genes of interest, such efforts are not necessary with comprehensively annotated, reference-quality assemblies. Third, reference-quality assemblies are a resource for studying gene regulation by non-coding RNAs and cis-regulatory elements. The high completeness enables a comprehensive mapping of functional genomics data such as miRNA read data and epigenomic data (e.g. ChIP-seq, ATAC-Seq), and the high contiguity is crucial for assigning regulatory regions to putative target genes and linking genotype to phenotype.

The six reference-quality assemblies coupled with methodological advances enabled us to address the long-standing question of the phylogenetic position of bats within Laurasiatheria. We used our comprehensive gene annotations to obtain the largest set of orthologous genes and homologous regions to date, which enabled us to explore the phylogenetic signal across different genomic partitions. Consistently, a variety of phylogenetic methods and data sets estimate that bats are a sister clade to Fereuungulata and highlight the importance of maximising the genetic coverage and ensuring that the appropriate models and data are used when reconstructing difficult nodes.

Our comprehensive and conservative genome-wide screens investigating gene gain, loss and selection provide candidates that are likely related to the unique immunity of bats. Furthermore, our screens reveal selection in hearing genes in stem Chiroptera, which is consistent with the hypothesis that echolocation evolved once in bats and was secondarily lost in Pteropodidae, but inconsistent with the alternative hypothesis that echolocation evolved twice independently within bats. As such, our analysis provides molecular evidence informing a long-standing question of when echolocation evolved. We further show that bats have a long coevolutionary history with viruses and identified unique mammalian viral integrations. Finally, we explored the non-coding genome in bats, where we found miRNAs that were novel to bats, lost in bats, or carried bat specific changes in their seed sequence. These important regulators of gene expression point to ancestral changes in the bat genome that may have contributed to adaptations related to the low incidence of cancer in bats, as well as developmental and behavioural processes.

While the six bat genomes presented here are an excellent starting point to understand the evolution of exceptional traits in bats, questions remain to be addressed in future studies, particularly because bats as a group exhibit such an incredible diversity. To resolve the phylogeny of the 21 currently-recognized bat families and to further understand the evolution and molecular mechanisms of traits that vary among bat families, such as longevity, mode of echolocation or diet, the Bat1K project aims at producing, in its next phase, reference-quality assemblies for at least one member of each of the 21 bat families. To enable efficient use of our reference-quality genomes, we provide all genomic and transcriptomic data together with all annotation and genome alignment in an open access genome browser (https://genome-public.pks.mpg.de) for download and visualization. These, and future bat genomes are expected to provide a rich resource by which to address the evolution of the extraordinary adaptations in bats and contribute to our understanding of key phenotypes including those relevant for human health and disease.

## Data availability statement

All data generated or analysed during this study are included in this published article and its supplementary information files. All genomic and transcriptomic data are publicly available for visualization and download via the open-access Bat1K genome browser (https://genome-public.pks.mpg.de). In addition, the assemblies have been deposited in the NCBI database and GenomeArk (https://vgp.github.io/genomeark/). Accession numbers for all data deposits can be found in the supplementary information files of this article.

## Code availability statement

All code has been made available on github. Details of the location can be found in the supplementary information files of this article. Other custom software is available upon request.

## Author contributions

MH, SCV, EWM and ECT conceived and supervised the project. MH, SCV, EWM and ECT provided funding. MLP, SJP, DD, GJ, RDR, AGL, ECT and SCV provided tissue samples for sequencing. ZH, JGR, OF and SW were responsible for nucleic acid extraction and sequencing. MP assembled and curated all genomes. DJ provided coding gene annotation and was responsible for coding gene evolutionary analysis. DJ provided multiple sequence and genome alignments. MH and DJ analysed UCE and genome completeness. DJ and MH established the Bat1K genome browser. ZH provided non-coding gene annotation and was responsible for non-coding gene evolutionary analysis. KL processed Iso-seq data and provided UTR annotation. ZH, KL, PD and SCV conducted miRNA target prediction and gene ontology enrichment. PD conducted miRNA functional experiments. GMH, LSJ, MS and ECT provided phylogenomic analyses. GMH and LMD were responsible for codeml analysis. DJ, MH, ZH, GMH, ECT, LMD and AP interpreted evolutionary analyses. BMK and MH developed the TOGA gene projection tool and BMK provided projections for non-bat mammals. ECS, LBG and AK provided EVE annotation and analysis. DR and KAMS provided TE annotation and analysis. EDJ provided support for sequencing of *Phyllostomus* and *Rhinolophus* genomes. DJ, ZH, MP, GH, MH, SCV, EWM and ECT wrote the manuscript. All authors provided edit and comment.

## Acknowledgements

This work was supported by the Max Planck Society, the German Research Foundation (HI 1423/3-1), and by European Research Council Research Grant (ERC-2012-StG311000). SCV was funded by a Max Planck Research Group Award, and a Human Frontiers Science Program (HFSP) Research grant (RGP0058/2016). GJ/ECT – funding from Royal Society/Royal Irish Academy cost share programme. LMD was supported, in part, by NSF-DEB 1442142 and 1838273, and NSF-DGE 1633299. DAR was supported, in part, by NSF-DEB 1838283. The authors would like to thank Stony Brook Research Computing and Cyberinfrastructure, and the Institute for Advanced Computational Science at Stony Brook University for access to the high-performance SeaWulf computing system, which was made possible by a National Science Foundation grant (#1531492).ECT was funded by a European Research Council Research Grant (ERC-2012-StG311000), UCD Wellcome Institutional Strategic Support Fund, financed jointly by University College Dublin and SFI-HRB-Wellcome Biomedical Research Partnership (ref 204844/Z/16/Z) and Irish Research Council Consolidator Laureate Award. EDJ and OF were funded by the Rockefeller University and the Howard Hughes Medical Institute. We thank the Long Read Team of the DRESDEN-concept Genome Center, DFG NGS Competence Center, c/o Center for Molecular and Cellular Bioengineering (CMCB), Technische Universität Dresden, Dresden, Germany, Sven Kuenzel and his team of the Max-Planck Institute of Evolutionary Biology in Ploen, Germany, the members of the Vertebrate Genomes Laboratory at The Rockefeller University, New York, US for their support. Special thanks to Lutz Wiegrebe, Uwe Firzlaff and Michael Yartsev who gave us access to captive colonies of *Phyllostomus* and *Rousettus* bats and aided with tissue sample collection.

