## Supplementary materials for "Six new reference-quality bat genomes illuminate the molecular basis and evolution of bat adaptations"

### Supplementary Information

**This text file includes:**

**Supplementary Methods**

**Supplementary Figure 1-13, 15-30**

**Supplementary Table 1-8, 16, 20-21, 23, 25-37, 40-41**

### 1. Samples, DNA extraction and genome sequencing

#### 1.1 Ethical statements and sample storage

The ethical statements of collecting and processing tissue samples for each species are listed as follows:

***Myotis myotis***: All procedures were carried out in accordance with the ethical guidelines and permits (AREC-13-38-Teeling) delivered by the University College Dublin and the Préfet du Morbihan, awarded to Emma Teeling and Sébastien Puechmaille respectively. A single *M. myotis* individual was humanely sacrificed given that she had lethal injuries, and dissected. ***Rhinolophus ferrumequinum***: All the procedures were conducted under the license (Natural England 2016-25216-SCI-SCI) issued to Gareth Jones. The individual bat died unexpectedly and suddenly during sampling and was dissected immediately. ***Pipistrellus kuhlii***: The sampling procedure was carried out following all the applicable national guidelines for the care and use of animals. Sampling was done in accordance with all the relevant wildlife legislation and approved by the Ministry of Environment (Ministero della Tutela del Territorio e del Mare, Aut.Prot. N°: 13040, 26/03/2014). ***Molossus molossus***: All sampling methods were approved by the Ministerio de Ambiente de Panamá (SE/A-29-18) and by the Institutional Animal Care and Use Committee of the Smithsonian Tropical Research Institute (2017-0815-2020). ***Phyllostomus discolor***: *P. discolor* bats originated from a breeding colony in the Department Biology II of the Ludwig-Maximilians-University in Munich. Approval to keep and breed the bats was issued by the Munich district veterinary office. Under German Law on Animal Protection, a special ethical approval is not needed for this procedure, but the sacrificed animal was reported to the district veterinary office. ***Rousettus aegyptiacus***: Egyptian fruit bats originated from a breeding colony at University of California (UC), Berkeley. All experimental and breeding procedures were approved by the UC Berkeley Institutional care and use committee (IACUC).

Sampled tissues were snap-frozen in liquid nitrogen immediately after dissection and were kept at -80ºC until further processed. Detailed information of samples is available in Table S25.

#### 1.2 Genomic DNA isolation and library preparation (PacBio, Illumina, HiC, 10X Genomics and Bionano)

##### 1.2.1 Phenol-chloroform extraction of genomic DNA

Snap-frozen tissues of all bat species were pulverized into a fine powder in liquid nitrogen. Powdered muscle tissue was lysed overnight at 55ºC in high-salt tissue lysis buffer (400 mM NaCl, 20 mM Tris base pH 8.0. 30 mM EDTA pH 8.0, 0,5% SDS, 100 µg/ml Proteinase K), and powdered lung tissue was lysed overnight in Qiagen G2 lysis buffer (Cat. No. 1014636, Qiagen, Hilden, Germany) containing 100 µg/ml Proteinase K at 55ºC. RNA was removed by incubating in 50 µg/ml RNase A for 1 hour at 37ºC. High molecular weight genomic DNA (HMW gDNA) was purified with two washes of Phenol-Chloroform-IAA equilibrated to pH 8.0, followed by two washes of Chloroform-IAA, and precipitated in ice-cold 100% Ethanol. Filamentous HMW gDNA was either spooled with shepherds hooks or collected by centrifugation. HMW gDNA was washed twice with 70% Ethanol, dried for 20 minutes at room temperature and eluted in TE. DNA molecule length was between 50 and 300 kb as shown by pulse field gel electrophoresis (PFGE) (Pippin Pulse, SAGE Science, Beverly, MA).

##### 1.2.2 Bionano agarose plug based isolation of megabase-size gDNA

Megabase-size gDNA was extracted according to the Bionano Prep™ Animal tissue DNA isolation soft tissue protocol (Document number 30077, Bionano, San Diego, CA) for liver tissue and according to the Bionano Prep™ Animal tissue DNA isolation fibrous tissue protocol (Document number 30071) for lung, muscle, and heart tissues. Fibrous tissues were mildly fixed in 2% formaldehyde and homogenized. Nuclei were enriched by centrifugation. Soft tissues were homogenized in a tissue grinder directly followed by a mild ethanol fixation. Nuclei or homogenized tissues were embedded into agarose plugs and treated with Proteinase K and RNase A. Genomic DNA has been extracted from agarose plugs and purified by drop dialysis against 1x TE. PFGE revealed mega-size DNA molecule length of 100 kb up to 500 kb. For *P. discolor*, additionally we extracted DNA using the Qiagen MagAttract HMW DNA kit (according to manufacturer guidelines) using 25-30 mg of tissue. The information regarding gDNA extraction is detailed in Table S26.

##### 1.2.3 PacBio long insert library preparation

Long insert libraries were prepared as recommended by Pacific Biosciences (PacBio, Menlo Park, CA) according to the guidelines for preparing size-selected 20 kb SMRTbell™ templates. The Megaruptor™ device (Diagenode, Liege, Belgium) was used for shearing 10-20 µg genomic DNA following the manufacturer's instructions. PacBio SMRTbell™ libraries were size-selected for large fragments using the SAGE BluePippin™ devise. SMRT sequencing was done on the SEQUEL system using sequencing chemistries 1.0 to 2.0. Movie time was 10 hours for all SMRT cells. The detailed information regarding PacBio sequencing statistics is available in Table S27.

##### 1.2.4 Bionano optical mapping of megabase-size gDNA

Megabase-size gDNA of *P. discolor* and *R. ferrumequinum* was labelled as described in the Bionano Prep™ Labeling NLRS protocol (Document Number 30024). DNA was tagged with two different enzymes each (BSPQI and BSSSI) to achieve the maximum labelling information. Labelled gDNA of these species was run on the Saphyr platform at the Vertebrate Genome Lab at the Rockefeller University. Megabase-size gDNA of the other four species (*M. molossus*, *M. myotis*, *P. kuhlii*, and *R. aegyptiacus*) was labelled as described in the Bionano Prep direct label and stain (DLS) protocol (Document number 30206). These DNAs were tagged with the nicking-free DLE enzyme. One flow cell of *M. molossus*, *M. myotis*, and *P. kuhlii* labelled gDNA was run on the Bionano Saphyr instrument at the MPI for Evolutionary Biology in Ploen, Germany. One flow cell of labelled *R. aegyptiacus* gDNA was run on the Bionano Saphyr instrument at the DRESDEN concept Genome Center (DcGC), Dresden, Germany. For all six species, at least 100X raw genome coverage was achieved.

##### 1.2.5 10x linked Illumina reads

Linked Illumina reads were generated using the 10x Genomics Chromium™ genome application following the Genome Reagent Kit Protocol v2 (Document CG00043, Rev B, 10x Genomics, Pleasonton, CA). In brief, 1 ng of long or megabase-size genomic DNA was partitioned across 1 Million Gel bead-in-emulsions (GEMS) using the Chromium™ devise. Individual gDNA molecules were amplified in these individual GEMS in an isothermal incubation using primers that contain a specific 16 bp 10x barcode and the Illumima® R1 sequence. After breaking the emulsions, pooled amplified barcoded fragments were purified, enriched and went into Illumina sequencing library preparation as described in the protocol. Pooled Illumina libraries were sequenced to at least 40X genome coverage on an Illumina HiSEQ4000 or an Illumina NovaSeq instrument at the MPI of Molecular Genetics in Berlin, Germany, using the 2x 150 cycles paired-end regime plus 8 cycles of i7 index. The 16 bp 10x barcodes allow the reconstitution of long DNA molecules by linking reads that carry the identical barcode. The detailed information regarding 10x Genomics sequencing is available in Table S28.

##### 1.2.6 Hi-C confirmation capture

Hi-C confirmation capture of *M. myotis*, *P. kuhlii*, and *R. ferrumequinum* was outsourced to Phase Genomics in Seattle, WA. Hi-C confirmation capture of *P. discolor* was done by Arima Genomics in San Diego, CA. For *M. molossus* and *R. aegyptiacus*, Hi-C confirmation capture and Illumina sequencing was done at the DcGC by applying the Arima Genomics Hi-C kit and sequencing on the Illumina Nextseq device.

##### 1.3 Pacific Biosciences long read transcriptome sequencing (Iso-seq)

###### 1.3.1 Total RNA extraction

The overview of tissues and RNA samples used for Iso-seq is available in Table S8. All tissues were lysed in TRIzol reagent (No. 15596-018, Carlsbad, CA). Total RNA extraction and purification was conducted either with a standard chloroform-isopropanol extraction protocol, using either the QIAGEN RNAeasy kit (Cat. No. 74104) or the ReliaPrep™ RNA cell miniprep kit (Cat. No. Z6110, Promega Madison, WI). The quality and quantity of all RNAs were measured using a Bioanalyzer 2100 or an Agilent 2200 Tapestation (Aligent Technologies, Santa Clara, CA). RIN values are given in Table S8.

###### 1.3.2 Library preparation

PacBio Iso-seq libraries were prepared according to the 'Procedure & Checklist - Iso-Seq™ Template Preparation for Sequel® Systems' (PN 101-070-200 version 05) without Blue Pippin size selection. Briefly, cDNA was reversely transcribed using the SMRTer PCR cDNA synthesis kit (Clontech, Mountain View, CA) from 1 µg total RNA and amplified in a large-scale PCR. Two fractions of amplified cDNA were isolated using either 1x AMPure beads or 0,4x AMPure beads. Both fractions were pooled equimolar and went into the Pacbio SMRTbell template preparation v1.0 protocol following the manufacturer's instruction.

###### 1.3.3 Sequencing

PacBio Iso-seq libraries were sequenced on the SEQUEL device with PacBio sequencing chemistry 3.0 and with 20 hours movie time. One SMRT cell was sequenced per Iso-seq library. Raw sequence yield (polymerase yield) for all Iso-seq libraries was between 18 and 32 Gb per SMRT with 624,989 to 732,879 reads per library. The *P. discolor* testes sample was sequenced on one SMRTcell using a 10-hour movie and chemistry 2.1, which resulted in 487,808 reads.

#### 2. Genome assembly

##### 2.1 Data sets and assembly inputs

The original data collection design was to produce 60X coverage in PacBio long reads, 50X in 10x Illumina read clouds, and 10X in Hi-C read pairs[1]. The idea was that the latter two technologies would be used for scaffolding contigs produced by an initial assembly of the PacBio reads into contigs. However, early on, it became clear that the yield of long read clouds with the 10x technology was very low. Even after switching to a plug-based DNA extraction method at a later timepoint, the yield of long clouds, while better, was still not cost efficient. Therefore, we abandoned the idea of using 10x read cloud data for scaffolding, albeit this data was very useful for base error correction and haplotype phasing, as each read cloud was itself phased. To compensate, and also in part based on our experience with the VGP project[2], we decided to generate a higher coverage in Hi-C read pairs for *P. discolor*, *R. aegyptiacus* and *M. molossus*. Furthermore, we decided to collect Bionano restriction mapped molecules and generate optical maps for all six bats since in the year after our initial proposal[1]. Bionano's optical map technology improved greatly in molecule length and was proved very powerful for scaffolding. In addition, the increased Hi-C coverage also gave us more scaffolding power, to the extent that the largest scaffolds were effectively chromosomes. We describe in the subsections 2.1.1 – 2.1.4 each of the four data sets for each of the six bats.

###### 2.1.1 PacBio reads

The target coverage for long read sequencing was 60X. In Table S29, we report statistics on all the raw data that we collected for each species, and all the data used for assembly which is the raw data except all those reads that were <4 kb in length. In the statistics for the raw data we did not count multiple reads of an insert in a given well, but only the longest read from each well. A gradual improvement is observed in yield per cell over the runtime of the project and the estimated coverage of the trimmed data is above or very near the 60X target for 5 of the 6 bats. The only exception is *M. molossus* with a trimmed data coverage of 52X; however, this species turned out to have an unexpectedly larger genome size of 2.3 Gb (versus ~2 Gb or less for all the others). Despite slightly lower read coverage, the PacBio reads for *M. molossus* are the longest (Fig. S21). The expected coverage reported is the total base pairs collected by the *post hoc* genome size of the resulting assemblies. The data for *P. discolor* was created at Rockefeller and Duke University and the other five bats were sequenced in Dresden, Germany.

##### 2.1.2 10x Illumina read counts

We collected about 50X Illumina reads organized into read clouds with the 10x Genomics technology[3] for *M. myotis*, *P. kuhlii*, and *R. ferrumequinum*. In this technology, a small number (*e.g.* 2-20) of ideally long molecules were isolated in an oil-immersion micro-well with a reagent payload that produces roughly 0.2-0.3X amplicons with the same barcode. The resulting library was then Illumina pair-read sequenced, resulting in "clouds" of reads with the same barcode. The reads were phased as the template was single stranded and it is noteworthy that a given cloud should map to a small number of regions whose size and number correspond to the molecules in the well. Locality information is thus rather indirect, but sufficient with large numbers of clouds to achieve moderately good assemblies[4].

We were expecting a large fraction of the clouds to be 100 kb or longer. The size distribution of the molecule lengths from which each cloud was derived cannot be measured directly. However, cloud reads can be mapped to the contigs produced by an initial assembly of the PacBio data in order to get a *post hoc* estimate of this distribution. This revealed that only 1% of the molecules were 100 kb or longer and most were much shorter. This can be seen clearly in Fig. S22, which plots the Nx values for the putative estimates of molecule length. Therefore, the coverage in long molecules was less than 1X and consequently this data provided very little scaffolding information. Given that the reads in a cloud must be inferred, they also tended to have a very high scaffolding error rate.

One could argue that the short molecule distribution was a protocol/lab error, but the data set produced for *P. discolor* by the VGL at Rockefeller University had the same characteristics (see Table S30 and Fig. S22). Later in the project, when we started using DNA extracted with the Bionano plug-based method (Document number 30077, Bionano, San Diego, CA), the molecule length distribution improved significantly with a tail that put about 50% of the data in molecules above 100 kb. However, this is still a relatively low yield of long molecules compared to the Bionano data which will be described section 2.1.3. In summary, while we produced ≥40X of 10x read clouds for all six bats, we only used this data for error polishing and phasing in our assembly pipeline described in the Section 2.2.

##### 2.1.3 Bionano restriction mapped molecules

Since Bionano proved to be producing very long restriction-mapped molecules (100-300 kb on average), we begun to produce this data for all six bats. Table S31 summarizes the gross statistics for each data set.

While one could use each molecule directly to scaffold contigs, we chose to first assemble each Bionano map using the company's restriction map assembler Solve (Document number 30205, Revision E, Bionano, San Diego, CA). This then gave us optical maps that we used in the sequence assembly pipeline. Table S32 summarizes the aggregate statistics for the assemblies. The data sets of *P. discolor* and *R. ferrumequinum* were performed with two enzymes and therefore had a distinct optical map assembly for each enzyme. One should note that while the coverage in molecule was the lowest for *R. aegyptiacus*, the average and N50 map lengths were the highest. This indicates that coverage alone does not determine the degree of assembly and may be less important than the distribution of read lengths, which was the best for *R. aegyptiacus*.

##### 2.1.4 Hi-C Illumina read pairs

Initially we contracted with Phase Genomics to produce 15X Hi-C data sets for *M. myotis*, *P. kuhlii* and *R. ferrumequinum*. Later in the project it became clear that Hi-C data is extremely well suited to give one the overall chromosomal view of a genome. Therefore, we increased the coverage of this data to >60X for the remaining three genomes, contracting one data set to Arima (*P. discolor*) and using the Arima kits in-house for the other two bats (*R. aegyptiacus*, *M. molossus*). Table S33 shows the Hi-C sequencing statistics.

#### 2.2 Assembly pipeline

*De novo* genome assembly was performed with DAmar (https://github.com/MartinPippel/DAmar). This assembler is based on an improved MARVEL assembler (https://github.com/schloi/MARVEL, commit ID: 5e17326)[5,6] and the integration of parts from the DAZZLER (DALIGNER commit ID: 233274a; DAMASKER commit ID: bc7e49c; DASCRUBBER commit ID: 3491b14; DAZZ_DB commit ID: 340fd89; DEXTRACTOR commit ID: 2f51ccb) [7]and the DACCORD code base (version: 0.0.14-release-20180525105343)[8,9].

To assemble the bat genomes, we performed the following steps: setup, PacBio read patching, assembly, error polishing, haplotype phasing, scaffolding and manual curation. Fig. S23 shows a schematic overview of the assembly pipeline.

##### 2.2.1 Setup phase

In the setup phase, PacBio reads were filtered by choosing only the longest read of each zero-mode waveguide (ZMW) and requiring subsequently a minimum read length of 4 kb. The resulting 6.7-11.4 million reads (52X - 70X coverage) for all 6 bats were stored in a DAZZLER database.

##### 2.2.2 Read patching

The patch phase detects and corrects read artefacts including missed adapters, polymerase strand jumps, chimeric reads and long low-quality read segments that are the primary impediments to long contiguous assemblies. We first computed local alignments of all raw reads. Since local alignment computation is the most time- and storage-consuming part of the pipeline, we reduced runtime and storage by masking repeats in the reads as follows. First, low complexity intervals, such as micro satellites or homopolymers, were masked with DBdust (all tools relate to the corresponding git repositories that are specified above). Second, tandem repeats were masked by using datander and TANmask. Third, as described in[10], we split all reads into groups representing 1X read coverage. For each group, we then aligned all reads against all others with daligner and masked all local regions in each read where at least 10 other reads aligned. The repeat masks were subsequently used to prevent k-mer seeding in repetitive regions when computing all local alignments between all reads.

Repeat masking can sometimes be disadvantageous, especially in highly repetitive regions of the genome. Low quality or noisy regions occur randomly in PacBio reads. In case such bad regions are spread into repetitive regions, they induce premature alignment breaks and the repeat mask prohibits further computation of local alignments within the repeat. This can result in alignment piles, where the alignment patterns for chimeric reads, strand jumps and noisy regions cannot be detected anymore. In the worst case the repetitive region is trimmed back in all PacBio reads, which creates dead ends in the following assembly step.

To overcome this problem, we used LAseparate to find proper alignment chains that prematurely end in repeat regions. For those alignment chains, we recomputed local alignments with the repcomp tool without using the repeat mask. Then we applied LAfix, which we further improved in the ability to detect chimeric breaks within repeat regions. Usually, the detection of chimeric reads is based on the alignment pattern that is caused by the chimeric break point, *i.e.* the set of reads that are aligned to the left of the chimer break point is disjoint with the set of reads that are aligned to the right. Furthermore, a chimeric break induces a clear wall of alignment ends and starts at both sides. In repetitive regions, especially in microsatellites, this is not necessarily the case and an interleaved alignment pattern may occur, which complicates the detection of exact break points. To resolve those issues for the bat assemblies, repetitive regions up to a length of 8 kb were analysed for chimers. Any subread which included a repetitive region that could not be spanned by at least three valid alignment chains was marked as chimeric read. This method identified between 0.51% (*P. kuhlii*) and 1.96% (*P. discolor*) chimeric reads. Due to sufficient read coverage, all of them were discarded.

##### 2.2.3 *De novo* assembly

In the assembly phase, we first calculated all overlaps between the patched reads using the same masking and alignment strategy of the patch phase. In addition, we applied an overlap chain rescue step. This step handles cases where a bad quality region was located at the tip of a subread, *i.e.* the interval from the tip to the minimum overlap length of the local alignment step (default 1.5 kb). In these cases, the bad quality region was not patched and therefore no proper overlaps were found. In order to avoid this behaviour, all alignment chains that prematurely ended due to a bad quality interval at subread tips were analysed with the Daccord tool forcealign. Forcealign tries to extend alignments by applying an increased error rate. For the bat assemblies this value was set to 35%. Only those alignments, which reached either a valid end in the A-read or in the B-read, were kept.

The subsequent steps were based on the generated overlaps and the original Marvel assembly pipeline[5,6]. First, the initial repeat annotation that only accounted for frequent repeats was updated by running LArepeat. Repeat regions were determined based on the coverage of the overlaps. If a potential repeat region had a coverage of more than twice the expected coverage of the genome, the region was annotated as a repeat. All following bases that had a coverage of at least 1.5 times the expected coverage were marked as repeat region, in order to compensate for coverage fluctuations in repeat regions. The end of a repeat region was defined as the point where the coverage fell below 1.5 times the expected coverage. The expected coverage was calculated from the overlaps itself and was not given as an argument.

The minimum overlap length of 1.5 kb can result in missing repeat annotations if the ends of reads are repetitive, but do not reach far enough into a repeat. To avoid this problem, we used TKhomogenize to transitively transfer the existing repeat annotations between reads.

The remaining gaps shorter than 100 bp within the pairwise alignments were stitched with LAstitch. Quality scores for all reads were then recalculated and trim tracks were generated by LAq. Next, we used LAgap to rescan the reads for remaining gaps (points which were not spanned by any overlap). Gaps at this stage usually exist due to left-overs of the “weak” regions in the reads that are not detected in previous stages. In order to resolve a gap, the overlaps from the shorter side were discarded. Gap resolution was followed by a round of trimming with LAq.

Based on the remaining overlaps and the updated trim track a final overlap filtering was performed with LAfilter, which discarded local alignments and repeat induced overlaps. For the six bats, we required that proper overlaps were at least 4 kb long and had at least 1000 anchor bases.

Based on the final set of overlaps, an overlap graph was built using OGbuild. Touring the overlap graph was performed by OGtour. The look-ahead for finding all potential paths was set to 10. Afterwards the touring paths were used to create raw-sequence contigs with tour2fasta. To correct base errors of the raw sequence contigs, we used the Marvel correction module, which is also part of DAmar.

In this step, only alignment piles from reads, which were used in the touring, were used to produce a consensus for the corrected contigs. This approach was very fast and reduced the error rate down to 1-2%.

The resulting corrected contigs were analysed and classified with CTanalyze, which separated the contigs into three different sets: primary, alternate and discarded. To this end, the contigs were aligned against each other and these alignments were used to derive a repeat mask. Further information, such as touring relation, patched-read mapping position, coverage, and repeat tracks, was integrated without realigning all reads against the assembly. The main task of CTanalyze is the haplotype separation into a primary contig set and an alternative contig set. For a reliable classification, different contig relations were combined into a multi relation matrix and a consensus classification was derived.

a) Graph touring relation: alternative contigs usually contain large structural variations that differ from the corresponding primary contigs. The graph touring also reports alternative contigs as bubbles or spurs.

b) Contig alignment relation: Contig overlap chains that allow for large structural variation are analysed for containment, bridging and forking relations.

c) Patched read intersection relation: If no reliable contig alignment chain could be found or the size of structural variation is larger than the alignment between two contigs, a) and b) may provide ambiguous or even no information. In that case, the original patched read overlap piles are analysed and if a major fraction of the PacBio reads is shared between two contigs then the smaller contig is assigned as a containment relation.

Afterwards putative primary contigs were further filtered and contigs that had an average coverage below 5, were more than 80% repetitive and were smaller than 20Kb were discarded. In addition to the contig classification, CTanalyze also reported potential issues, such as putative false joins, low coverage drops within contigs, and putative bridges between contigs. For the six bats, the potential issues (between 2-10 per species) were manually inspected and corrected if necessary.

##### 2.2.4 Error polishing

The primary and alternate contigs were further polished by using the raw PacBio reads and applying two rounds of Arrow (https://github.com/PacificBiosciences/GenomicConsensus.git) polishing. Arrow decodes polished sequence in capitals, whereas unpolished sequence was represented in lower case bases. DAmar contigs tend to end within large repeats, which could not always be fully polished. To facilitate the later scaffolding process, uncorrected contig ends that remained after the second polishing round were trimmed back.

To further correct base errors and reduce remaining length errors in homopolymer regions, 10x read clouds were used. To map 10x read clouds to the Arrow-polished contigs, the 10x Genomics Longranger align pipeline (https://github.com/10XGenomics/longranger, version 2.2.0) was applied, which uses the barcode-aware mapping tool Lariat. Afterwards the variant detector FreeBayes (version 1.2.0)[11] detected polymorphic positions and fixed erroneous non-polymorphic sites in the reference sequence using bcftools consensus (version 1.9) (https://github.com/samtools/bcftools). 10x read cloud polishing was iteratively applied in two rounds.

##### 2.2.5 Haplotype phasing

So far, the assembly pipeline did not account for heterozygous events at the base level and the contigs did contain a mixture of both alleles. To address this problem, the 10x Genomics Longranger wgs pipeline with FreeBayes (version 1.2.0)[11] as the variant caller was used. Based on the phased VCF output file, bcftools consensus was used to produce locally-phased primary contigs. Depending on the

10x molecule lengths, the phased N50 of the bats ranged from 0.9 Mb (*P. discolor*) to 6 Mb (*M. molossus*).

##### 2.2.6 Bionano scaffolding

###### 2.2.6.1 *De novo* assembly

The Bionano raw molecules were assembled with Bionano Solve (Version 3.3) that offered command line tools for analysing Bionano data. An additional signal to noise filtering (filter_SNR_dynamic.pl) was required for two bat species (*R. ferrumequinum*, *P. discolor*) for which data from BSSSI and BSPQI nicking enzymes of the Saphyr system was available. The other four bat species, for which the newer DLE-1 direct labelling technique was used, did not require a SNR filtering step.

To assemble the optical maps of the six bats, we used all molecules ⩾150 kb that additionally have at least 9 sites. The number of extension and search operations was set from the default 5 to 10, but after the 7th iteration most optical map assemblies converged, and no major changes were recognized. For each bat, we generated two maps using two different assembly option argument files: nonhaplotype_noES_saphyr.xml (noES) and nonhaplotype_saphyr.xml (ES). The noES option file resulted in more contiguous assemblies with higher N50 values. The nonhaplotype_saphyr.xml option file resulted in assemblies that were larger due to uncollapsed heterozygous maps. Both assembly versions were created for all six bats and evaluated in the following hybrid scaffolding step.

###### 2.2.6.2 Hybrid scaffolding

The input to the Bionano hybrid scaffolding were the locally phased primary contigs, which were *in silico* digested by using the corresponding restriction sites (DLE-1: *M. myotis*, *P. kuhlii*, *R. aegyptiacus*, *M. molossus*; BSPQI and BSSSI: *R. ferrumequinum*, *P. discolor*) and the previously created Bionano assemblies. For *R. ferrumequinum* and *P. discolor*, the two-enzyme hybrid scaffolding procedure was performed using the wrapper script runTGH.R of Bionano Solve. The other four bat assemblies were scaffolded with hybridScaffold.pl, which is also part of the Bionano Solve command line tools. The conflict filter level for Bionano cmaps and contig cmaps were set to 2, *i.e.* if the genome map does not have long molecule support at the conflict junction, then the map is cut. Otherwise the sequence fragment is cut.

Scaffolds that were based on the noES-Bionano assembly had more contigs integrated and therefore had a higher scaffold N50 compared to scaffolds that were based on the ES-Bionano assembly. The correctness of the scaffolds was validated with Bionano Access and manual inspection of the raw molecule coverage that supported each contig integration. Furthermore, the Hi-C reads were mapped to the Bionano scaffolds, and HiGlass[12] was used to explore the genomic contact matrix. With the exception of *R. aegyptiacus*, the noES-Bionano assembly outperformed the ES version for the other five bats. For *R. aegyptiacus*, the noES-based scaffolds included a 110 Mb scaffold, which contained a 60 Mb gap. When inspecting the ES-based scaffolds the gap was filled with a 60 Mb contig, which could not be integrated when using the noES-Bionano assembly. This could be explained by the fact that *R. aegyptiacus* was the only bat for which we could not generate the Bionano data from the same individual. As the Hi-C data indicated that all other noES-Bionano scaffolds were valid, the missing 60 Mb contig was manually integrated into the gap location.

##### 2.2.7 Hi-C scaffolding

To map the Hi-C Illumina read pairs to the previously created Bionano scaffolds the program bwa (version 0.7.17-r1194)[13] was used. The alignments were filtered according to the Arima filtering protocol (https://github.com/ArimaGenomics/mapping_pipeline). The resulting alignments were scaffolded with the Hi-C scaffolder Salsa2 (version 2.2)[14]. The clean option that detects misassemblies in the input assembly was enabled.

##### 2.2.8 Manual curation

To visually inspect and validate the final scaffolds, we used the web-application HiGlass. To this end, Hi-C reads were mapped with bwa (version 0.7.17-r1194) to the Salsa2 scaffolds and the alignments were filtered und successively converted into multi-resolution cooler files.

Our inspection revealed that the overall scaffolding quality was already quite high (Fig. S24). However, visualization revealed a few false joins and unique off-diagonal interaction patterns that suggested joining scaffolds. Scaffolds were split if the Hi-C read mapping density around the diagonal was not supported (Fig. S24 – highlighted with ellipse 1). Scaffolds were joined if the read mapping density in the off-diagonal was increased and the map resolution allowed a unique placement (Fig. S24 - highlighted with ellipse 2). For each bat, up to 10 splits and 10 joins (*P. discolor*) were manually performed and the curated scaffolds were validated again by HiGlass (version 0.6.3) (Fig. S24).

#### 2.3 Assembly results

After applying the DAmar assembler to the PacBio reads, two rounds of Arrow and FreeBayes polishing, and haplotype phasing in combination with the 10x read clouds, the output of this process (described above) is a collection of contigs that are either considered primary, alternate, or contigs that we discarded due to their small size. For all 6 bats, we obtained assemblies comprising just several hundred primary contigs. To determine N50 values, we used the Perl script assemblathon_stats.pl that is part of the Assemblathon 2 analysis pipeline[15] (https://github.com/ucdavis-bioinformatics/assemblathon2-analysis), defining assembly gaps as runs of ⩾10 N's. The N50 of the contigs was > 10 Mbp in every case and correlated with the average read length of the data set. The number of contigs also roughly inversely correlated with read length adjusting for overall genome size (*e.g. M. molossus* had the highest average read length but 396 contigs versus only 260 for *R. aegyptiacus* because the latter has a 1.9 Gbp genome whereas *M. molossus* has a 2.3 Gbp genome). Table S34 gives the number of contigs, total base pairs in these contigs, and N50 length for each species and each contig class. The number of alternate contigs was largely on the same order as the number of primary contigs but varied significantly, presumably reflecting the level of structural heterogeneity between the haplotypes of the individual's genome.

The results of scaffolding the locally-phased primary contigs with the assembled Bionano optical maps are shown in Table S35. In all cases, we obtained large scaffolds by just using Bionano optical maps. The assemblies of *M. myotis* and *P. kuhlii* were more fragmented, which reflects the less contiguous assembled maps for these two genomes. Table S35 also shows that few contig breaks were introduced by Bionano scaffolding.

After Bionano scaffolding, we generated final scaffolds using the Hi-C data. This scaffolding step again joined and broke scaffolds, but scaffold breaks typically occurred only at the tips, as shown by the small "delta" values in Table S36. For all six bats, Hi-C scaffolding substantially increased scaffold N50 sizes by 25-100%. We found that the bats for which we generated only 15X of Hi-C data (*M. myotis*, *P. kuhlii*, and *R. aegyptiacus*) had slightly smaller scaffold N50 values than the bats for which 60X was generated (Table S36). This suggests that a higher coverage of Hi-C read pairs is desirable for scaffolding and we aim at generating 60X or more in future projects.

The results of manually splitting or joining scaffolds based on the plots of the Hi-C maps are shown in Table S3. It is noteworthy that manual curation detected very few conflicts leading to scaffold breaks.

To assess whether our scaffolds often represent chromosomes, we used available karyotypes for each species to estimate the length of each chromosome (https://git.mpi-cbg.de/dibrov/chromosome_size). It should be noted that the length of small chromosomes (less than 20 Mbp) is hard to estimate as these chromosomes are just represented by a small blob in the karyotype.

We plotted the estimated lengths of the chromosomes against the length of our final scaffolds. The karyotype estimate was always larger than the next largest scaffold, presumably because the scaffolds did not have accurate gap lengths for the large centromere between chromosome arms. Nevertheless, as shown in Fig. S25 for three bats, we found a good agreement between estimated chromosome lengths and the lengths of our scaffolds. Specifically, the correlation coefficient between the N karyotypes of a species, and the N largest scaffolds is given in Table S37 and was always above 0.98, except for *P. kuhlii* for which manual curation was not performed. We further examined the remaining scaffolds and characterized this residual into three informal categories: Cliff(x) = no scaffolds over 2 Mb remain and x karyotypes have no corresponding scaffold, Incline(x) = x scaffolds over 2 Mb remain but all were significantly smaller (25% or less) of the smallest karyotype, Tail(x) = x scaffolds over 2 Mb remain and the size distribution gradually declined from the last assigned to a karyotype. We found that, with the exception of *M. myotis* and *P. kuhlii*, all other assemblies had Cliff or Incline endings, suggesting that almost all the scaffolds corresponded to chromosomes. Furthermore, with the exception of *P. kuhlii* (87%), more than 95% of the assembly was in the largest N chromosomes (Table S37).

Table S2 shows the final assembly statistics for our locally-phased primary contigs and our scaffolds of these. QV metrics were computed by mapping all the 10x read data to the final contigs and analysing discrepancies with the Illumina reads. This showed that our assemblies achieved the desired QV40 metric. Together, our contig and scaffold N50 values and our estimation that ≥90% of the scaffolds correspond to chromosomes for five of the six bats, the genome assemblies of five bats exceed the VGP standard of 3.4.2.QV40 and approach in fact 4.5.2.QV40. For *P. kuhlii*, the achieved metric is 4.5.1.

### 3. Genome annotation

##### 3.1 Protein-coding gene annotation

###### 3.1.1 Overview

To annotate coding genes, we used a variety of approaches and data to obtain evidence of coding genes in the bat genomes. These evidences comprise (i) projecting genes annotated in another mammal to our bat genomes via whole genome alignments, (ii) aligning protein and cDNA sequences of related mammals, (iii) mapping RNA-seq and Iso-seq data obtained for the six bats, and (iv) *de novo* gene predictions using a bat-specific gene model. These evidence were integrated into a consensus gene set, which was further enriched for high-quality isoforms. All individual evidence and the final gene set can be visualized and obtained from the genome browser. Below, we detail how each of the evidence was obtained and how they were integrated.

###### 3.1.2 TOGA projections

As the first evidence, we projected annotations of coding genes from multiple reference genomes to our bat genomes using TOGA (Tool to find Orthologs from Genome Alignments, last commit: 02/05/2019). Briefly, TOGA takes as input pairwise genome alignment chains between a designated reference and query genome[16], coding transcript annotations for the reference species and a file linking gene and transcripts isoforms. For each gene, TOGA identifies the chain(s) that aligns the putative ortholog in the query using synteny and the amount of aligning exonic and intronic sequence. To obtain the locations of coding exons of this gene, TOGA extracts the genomic region corresponding to the gene on this chain from the query assembly and uses CESAR 2.0 (Coding Exon Structure Aware Realigner)[17] in multi-exon mode.

We applied TOGA to the genome alignments (see above) to project the Ensembl (version 96, last accessed: 26/04/2016) gene annotation for human (hg38) and mouse (mm10) to our six bats. Furthermore, the *M. lucifugus* (myoLuc2 assembly) Ensembl (v96) annotation was projected to our *M.*

*myotis* assembly and the final gene annotation of *M. myotis* produced for this project was projected to the other 5 bat species. The number of projected genes for each of the six bats is listed in Table S37.

##### 3.1.3 Alignments of protein and cDNA sequences of related bat species

As the second evidence for coding genes, we aligned protein and cDNA sequences of related species to our six bat assemblies. For each of the six bats, we downloaded protein and RNA transcript sequences from NCBI or Ensembl for one other close-related bat species that has annotated genes (Table S6). Protein and transcript sequences were filtered to retain only those with matching peptide and mRNA sequence. Then, we used GenomeThreader (v1.7.0)[18] to simultaneously align protein and mRNA sequences to the respective target genome. GenomeThreader was run using the Bayesian Splice Site Model (BSSM) trained for human and default parameters aside from those detailed below. For protein alignments, we used a seed and minimum match length of 20 amino acids (prseedlength 20, prminmatchlen 20) and allowed a Hamming distance of 2 (prhdist 2). For the transcript alignments, we used a seed length and minimum match length of 32 nucleotides (seedlength 32, minmatchlen 32). At least 80% of the protein or mRNA sequence was required to be covered by the alignment (-gcmincoverage 80), and potential paralogous genes were also computed (-paralogs). For *M. molossus*, these stringent parameters produced much fewer gene predictions compared to the other five bats, likely due to the increased phylogenetic distance between *Molossus* and *Miniopterus* (from which we used annotated genes) compared to the other species pairings. Therefore, we performed an additional GenomeThreader run for *M. molossus* using less stringent parameters (default parameters for the seed, minimum match lengths and Hamming distance). The stringent alignments were provided as hints to Augustus (below), while the less stringent gene predictions were used for consensus gene prediction. For the other 5 genomes, the stringent alignments provided hints for Augustus and were used for consensus gene prediction. The number of filtered gene alignments for each of the six bats is listed in Table S37.

##### 3.1.4 Transcriptome data

As a third evidence for genes, we used RNA-seq and Iso-seq transcriptomic data that were mostly newly-generated for each of the six bats in this project. Table S7 provides details of the tissues used to generate transcriptome data and lists Sequence Read Archive (SRA) accession numbers used to download previously generated data.

For RNA-seq, reads were stringently mapped to the respective genome using HISAT2 (v2.0.0)[19], removing reads with greater than 5% ambiguous characters (-n-ceil L,0,0.05), disallowing discordant and mixed alignments (--no-discordant --no-mixed), and using the --dta (downstream transcriptome assembly) flag. The resulting SAM file was sorted and converted to BAM format using Samtools (v1.9)[20]. Transcripts were assembled using StringTie (v1.3.4d)[21] with default settings.

Since RNA-seq data also contains non-coding transcripts, we next filtered for transcripts that contain an open reading frame (ORF) and are not potential nonsense-mediated decay (NMD) targets. To this end, we used the Transcriptome Annotation by Modular Algorithms (TAMA) package (https://github.com/GenomeRIK/tama.git; accessed 21/5/2019; commit 58f9d98), which predicts ORFs for all assembled transcripts. Putative peptide sequences were queried against the Swissprot database (downloaded 20/05/2019) using blastp from the BLAST+ suite (v2.6.0) with default parameters[22]. BLAST results were parsed, designating a coding sequence (CDS) and mapping this to the corresponding exon structure of each transcript. Transcripts identified as full length by TAMA were retained and used as input for consensus gene models. The number of transcripts obtained from RNA-seq for gene annotation is reported in Table S37.

We used our Iso-seq data to produce high quality ORF predictions. To this end, raw reads were first processed using the IsoSeq3 pipeline (version 3.1.0) (https://github.com/PacificBiosciences/IsoSeq3) with the arrow polish flagged on. The resulting high-quality transcripts (HQ) (full-length and supported by more than one read) and FLNC reads (full-length non-chimeric reads before the clustering step) were further processed in parallel. The FLNC and HQ PacBio BAM files were converted into FASTA format using Bamtools (version 2.4.1) and aligned to the reference genome with Minimap2 (-t 16 -ax splice -uf --secondary=no -C5, version 2.10-r784-dirty). The resulting BAM files were filtered to retain only primary alignments using Samtools (version 1.9). TAMA collapse (https://github.com/GenomeRIK/tama.git) was applied to both HQ and FLNC primary alignments to predict non-redundant transcript set.

The resulting transcript coordinates were used to extract corresponding genomic sequences with Bedtools (getfasta –split –name -s, version v2.27.1) for both the HQ and FLNC set. The ORF prediction was run in two steps. First, the TAMA-GO package was run on HQ transcript sequences (extracted in an earlier step, see above) resulting in the annotation of putative ORF in each transcript. The putative CDS coordinates were used to determine and filter out the potential targets of nonsense-mediated decay pathways[23] by removing all transcripts that have more than one intron in the 3'UTR or transcripts in which an intron is located more than 50 bp from the stop codon. The resulting set (HQ.nonnmd) was used to train an ANGEL ORF prediction model (https://github.com/PacificBiosciences/ANGEL). The FLNC.nonnmd set was produced using TAMA-GO in the same way as described above and was used as input for ANGEL in prediction mode (output_mode=best --min_angel_aa_length 100 --min_dumb_aa_length 100) with the model trained in the previous step. The resulting annotations were used to split the FLNC.nonmd transcript set into three groups: (i) ANGEL positive (with evidence of an ORF predicted by ANGEL); (ii) ANGEL negative but with blastp hits; (iii) ANGEL negative and no blastp hits. ANGEL positive, full length transcripts were provided as gene predictions for consensus gene prediction. The number of putatively coding transcripts obtained with Iso-seq for each of the six bats is listed in Table S37.

##### 3.1.5 *De novo* gene prediction:

As a fourth piece of evidence, we generated *de novo* gene predictions using Augustus (v3.3.1)[24]. To this end, we first trained a bat-specific Augustus model using *M. myotis* as a representative species and the BRAKER pipeline (v2.1)[25]. BRAKER uses extrinsic evidence (RNA sequencing and/or proteins from a close-related species) as training data and performs iterative gene prediction to train model parameters. We used an earlier contig assembly of *M. myotis* and provided GenomeThreader alignments of *M. lucifugus* proteins (downloaded from Ensembl, date: 8/8/2018) and a BAM file of mapped *M. myotis* RNA-seq data from several tissues (kidney, liver, heart and brain) as input to BRAKER. The resulting "bat" model was used in subsequent Augustus runs.

Augustus is able to use extrinsic evidence as hints when predicting genes in a newly-sequenced genome. We compiled the following data as Augustus hints. RNA-seq was used to produce intron hints using the Augustus bam2hints module with the introns-only flag. RNA-seq derived hints was given a 'priority' of 4. High quality ORFs predicted from Iso-seq transcripts and classified as positive using ANGEL (described above) were converted to BAM format, and bam2hints was used to produce intron, exon and exonpart hints. Iso-seq derived hints were given a priority of 6. GenomeThreader alignments were converted to hints using the align2hints.pl script provided in the BRAKER distribution. This produced CDSpart, intron, start and stop hints that were given a priority of 4. Identical hints were merged using the join_mult_hints.pl script from Augustus. Further, human (Gencode version 27) and mouse (Gencode version 16) gene annotations were provided as high weight CDS and intron "manual" hints when running Augustus in comparative mode.

Augustus was run in two modes, in single genome mode for each of our six assemblies and once in comparative mode using a multiple genome alignment. For single genome mode, human TOGA projections were used to divide each genome into approximately 2.5 Mb regions with 250 kb overlap, avoiding splits inside putative genes. Augustus was run with the trained "bat" model and a custom extrinsic config file containing the bonus and malus parameter for each hint type. Alternative splice forms were predicted from evidence (alternatives-from-evidence), and AT/AC splice sites were allowed if supported by hints (allow_hinted_splicesites=atac). The resulting GTF files of gene predictions for each region were merged using the Augustus joingenes module.

For Augustus in comparative mode, we used the multiple genome alignment (MAF format) produced by MultiZ (v11.2) with *M. myotis* as the reference species as input. We used the split regions determined for single genome mode for *M. myotis* to split the MAF file into non-overlapping 2.5 Mb regions. A database containing the genomes for the 6 bat species, human and mouse, and the hint data was constructed, and a custom extrinsic config file was provided. All genomes were provided as soft-masked (repetitive sequence indicated as lower case letters). The phylogeny with branch lengths as estimated using IQ-Tree (see section 4.2 "Phylogenetic inference and divergence time estimation" below) was trimmed to contain only the species in the MAF file, and also provided to Augustus in Newick format. GTF files of gene predictions for each species were merged using the Augustus joingenes module. The number of predicted transcripts for each of the six bats is listed in Table S37.

##### 3.1.6 Integrating all gene evidence into a final gene annotation

We used EVidenceModeler (v1.1.1)[26] to integrate the gene evidences from TOGA projections, Genome Threader alignments, Augustus gene models and transcript ORF predictions from Iso-seq and assembled RNA-seq reads into a consensus gene set. Augustus gene predictions from the single and comparative mode were designated as *ab initio* predictions and given weights 2 and 1 respectively. GenomeThreader alignments were designated protein alignments with weight 2. The TOGA projections were given as "other" predictions all with weight 8. Transcript ORF predictions from assembled RNA-seq were filtered for those labelled as full length by TAMA and were provided as "other" predictions with a weight of 10. ANGEL positive ORF predictions from Iso-seq data that were also labelled as full length by TAMA were provided as "other" predictions with a weight of 12. Genomes were partitioned using EVidenceModeler into 1 Mb chunks with 150 kb overlap. Consensus gene models were called for each partition. EVidenceModeler output was converted into GTF format using in-house Perl scripts. We used the joingenes function from Augustus to combine all outputs into a consensus gene set.

EVidenceModeler does not, by default, produce consensus gene models for genes that are nested in an intron of another gene. Although this behaviour can be enabled via a parameter, it also produces a high number of likely false positive gene models. Therefore, in order to rescue these intronic genes, we incorporated TOGA projections from human and mouse with no CDS overlap to any already-detected consensus gene model. TOGA projections were only considered for incorporation if the gene began and ended with canonical start and stop codons and contained no internal stop codons. Further, only APPRIS "Principal" isoforms[27] were considered. Transcripts were added first from human and then from mouse.

As we used the *M. myotis* gene annotation as input for TOGA projections to the other five bats, we visualised the gene annotation in a genome browser and screened for obvious annotation errors such as potential genes lacking a consensus model, fused or split genes. Manual refinement and correction of a few loci was performed where necessary.

EVidenceModeler produces a single consensus gene model for each locus, and therefore will not annotate exons or splice sites that only occur in alternative isoforms of the same gene. We therefore used evidence sources of high confidence to incorporate isoforms to already-detected gene loci if an isoform provided novel splice information relative to the annotated consensus isoform. We did not incorporate isoforms that are potential NMD targets, defined as transcripts having more than two introns in the 3'UTR or transcripts in which an intron is more than 50 bp downstream of the stop codon[23]. Isoforms predicted from Iso-seq data were added as priority, followed by RNA-seq derived transcripts and finally TOGA projections. RNA-seq transcripts were filtered to remove those which may represent 5' degraded transcripts, identified as a transcript with no novel splice sites and a start codon nested within a previously annotated exon or having more than two 5' non-coding exons. A TOGA gene projection was only considered if it was an APPRIS Principal isoform, had canonical start and stop codons, no internal stop codons and that all coding exons from the reference were projected.

##### 3.1.7 Prediction of 3'UTR sequences from Iso-seq transcripts

3'UTR sequences were predicted using FLNC.nonmd Iso-seq transcripts set as follows. First-pass 3'UTR coordinates were created using CDS predictions, from the stop codon to the end of the transcript. Then, a custom script was run to cluster all 3'UTR coordinates per gene locus that shared the stop codon coordinate but varied in the 3' most (end of 3'UTR) coordinate. For these cases, we chose the longest 3'UTR per cluster and assigned it a weight, defined as the number of Iso-seq transcripts that shared this stop codon coordinate. Next, if more than one clustered 3'UTR per gene locus was found, the one with the highest weight was selected. Finally, the set of the candidate 3'UTRs was compared to gene annotations of our bats and only the sequences with a stop codon within a 100 bp window from the end of the annotated CDS of a gene were retained.

##### 3.1.8 Filtering transcripts for coding potential and assigning gene symbols

Manual inspection showed that integrating transcripts from a variety of evidence also included a number of genes that are unlikely to code for a protein and may represent non-coding or erroneous genes. In particular, many Iso-seq transcripts only had short and non-conserved predicted ORFs, indicative of non-coding genes, but were included by EVidenceModeler because of the high weight we gave this high-confidence transcript evidence. To remove putative erroneous or non-coding genes, all putative peptide sequences were queried against the Swissprot database using blastp with a minimum E-value of $1e^{-10}$. Sequences with no match to a mammalian sequence in the database were removed if they were smaller than 120 amino acids. Reported hits were further filtered, only retaining a match which covered >75% of the query sequence and >50% of the subject, and >50% positive scoring matches. We assigned the human gene symbol to an annotated gene in bats if the CDS overlapped between the locus and a single TOGA-projected human gene. Genes for which we could not assign a gene symbol based on TOGA projections were assigned a symbol based on the previously computed BLAST alignments. BLAST alignments were divided into complete matches (>65% query coverage, >70% subject coverage, and >30% identity), and partial hits (>75% query coverage, >50% subject coverage, and >50% positive scoring matches). Gene symbols were retrieved for all matches with a bit-score no less than 85% the value of the top hit. Gene symbols from the majority of retained hits were assigned to a gene. Genes with no complete matches were assigned a symbol from the partial matches, with an appended L to indicate the partial match. When multiple loci were assigned the same symbol, they were distinguished by incrementing a trailing alphanumeric character.

##### 3.1.9 Computing Annotation Completeness

In order to assess the completeness of the protein coding annotation, we used BUSCO (version 3)[28] with the mammalian (odb9) protein set. Predicted peptide sequences from the six bat species along with annotated peptide sequences for seven other mammal species, including human (hg38), mouse (mm10), pig (susScr11), cow (bosTau8), cat (felCat8), horse (equCab3) and dog (canFam3), were downloaded from Ensembl (version 96) (Table S1). BUSCO was run in protein mode, and the number of complete, fragmented and missing genes were compared across assemblies.

##### 3.2 Analysis of ultraconserved elements

To assess the completeness of non-exonic regions in mammalian assemblies, we determined the number of aligning ultraconserved elements per assembly. Since the 481 ultraconserved elements (UCEs) were originally defined as genomic regions ≥200 bp that are identical between human, mouse and rat[29], we did not use the human and mouse genomes in this comparison as by definition all UCEs are present in these assemblies. As in a previous study[5], we focused on the 197 UCEs that do not overlap exons according to the human Ensembl gene annotation and that align to chicken (galGal5 assembly) and teleost fish (zebrafish danRer10, medaka oryLat2). Given their strong conservation across vertebrates, we expect that these 197 vertebrate non-exonic UCEs are present in mammalian genomes.

To align these 197 ultraconserved sequences against mammalian genomes, we used Blat (v36x2)[30] with sensitive parameters (-minIdentity=60 -minScore=30 -minMatch=1 -stepSize=8 -

mask=lower). We kept those Blat hits where the alignment had a minimum identity of 85% and at least 150 of the ≥200 bp in the ultraconserved sequence aligned. This number of aligning UCEs is shown in Fig.1d.

To investigate why 15 UCEs did not align with these criteria to individual assemblies, we inspected these UCEs in the human UCSC genome browser in the context of a multiple genome alignment of mammals and pairwise alignment chains. We used the nearest up- and downstream aligning block in the chain to determine whether the UCE maps completely or partially to a query genomic locus that includes an assembly gap, as shown in Fig. S1. These UCEs were classified as 'missing due to assembly gap'. For *M. myotis* and *P. kuhlii*, we found one and three UCEs respectively that did align but exhibited substitutions and smaller insertions/deletions that decreased the alignment identity below our 85% threshold. For these four UCEs, we used blast with default parameters for the 10x Genomics Illumina reads and daligner with "-A -k11 -w5 -h35 -e.7 -l100 -M64" parameters for PacBio reads to confirm that (i) the genomic sequence aligning to the UCE is supported by both PacBio and by Illumina reads of the respective bat species and (ii) that the human ultraconserved sequence does not have a better match in any of the read data acquired for the respective bat. To further support real sequence divergence in an otherwise ultraconserved element, we aligned the sequences of close-related bats with sequenced genomes and found that most mutations were shared among other independently-sequenced bats. The three diverged UCEs are shown in Figs. S2, S4-5. Table S5 lists the details of all these 15 UCEs.

##### 3.3 Repetitive element annotation

We annotated each genome for transposable elements following the methods in [31]. Briefly, each assembly was mined for potential novel TEs using RepeatModeler[32]. The resulting putative TE libraries were masked with RepeatMasker (v4.0.9)[32] and the results then processed using calcDivergenceFromAlign.pl in the RepeatMasker package to generate Kimura-2-parameter (K2P) distances. We presumed that younger TE families, defined as consensus sequences having hits with K2P distances less than 6.6% (approximating ~30 Myrs or less since insertion, based on a general mammalian neutral mutation rate of $2.2x10^{-9}$)[33], were lineage-specific and potentially undescribed. Consensus sequences were also filtered for size (>100 bp), subjected to iterative homology-based searches against the genome, and manually curated[31]. For each iteration, new consensus sequences were generated to match the top 50 blast hits. Bioinformatically, this was accomplished by aligning with MUSCLE (v3.8.31)[34], trimming the alignments with trimal (-gt 0.6 -cons 60) (v1.3)[35], and estimating a consensus with the EMBOSS script 'cons' (-plurality 3 -identity 3)[36]. Files with fewer than 10 blast hits were discarded. Curation of the estimated consensus by eye ensured accuracy by preventing inclusion of single indels and observing 5' and 3' TE ends to confirm the full length of each element in each alignment.

To confirm TE type, each TE was compared to three online databases: blastx to confirm the presence of known ORFs in autonomous elements, RepBase (v20181026) to identify known elements, and TEclass[37] to predict the TE type. We also used structural criteria as follows. For DNA transposons, only elements with visible terminal inverted repeats were retained. For rolling circle transposons, we required elements to have an identifiable ACTAG at one end. Putative novel SINEs were inspected for a repetitive tail and A and B boxes. LTR retrotransposons were required to have recognizable hallmarks such as TG, TGT or TGTT at their 5' and the inverse at the 3' ends. Finally, duplicates were removed via the program cd-hit-est (v4.6.6)[38,39] if they did not pass the 80-80-80 rule as described in [40].

The complete TE library for each bat was combined with a vertebrate library of known TEs in RepBase (v20181026). This library is available as Data S1. RepeatMasker was used to mask the genomes with this custom library. Postprocessing of output was performed using a custom script, RM2Bed.py (https://github.com/davidaray/bioinfo_tools), which eliminated overlapping hits and converted to Bed format. The same methods were used to analyse seven mammalian outgroups (Table S1). The resulting data is shown in Fig. 1g and Table S38.

##### 3.4 Annotation and analysis of endogenous viral elements (EVE) and endogenous retrovirus (ERV)

###### 3.4.1 EVE annotation and analysis

We analysed the bat genomes and seven additional mammalian genomes as outgroups (Table S1) for the presence of endogenous viral elements (EVEs). Mammalian genomes were converted to nucleotide BLAST databases[41]. A comprehensive library of viral proteins (Table S39) was queried against the mammalian genomes using tblastn (maximum E-value 0.001; maximum number of 100 alignments reported). The viral proteins span the viral classes and families listed in [42] and were updated to the current versions of the reference sequences for each virus. The results were manually inspected and total viral insertions under 100 amino acids in length were discarded. Reciprocal BLAST searches were run for each hit, with the best hit viral family considered the true identity. BLAST hits in regions annotated as functional mammalian genes were considered false hits. Nucleotide sequences for each identified viral family, plus extant representatives of the family and previously identified EVEs, were aligned with Aliview[43].

###### 3.4.2 ERV annotation and analysis

All 6 bat genomes and the 7 additional mammalian genomes were searched with local BLAST[41] using 14 probes of the viral proteins gag, pol and env from each genus of Retroviridae: alpha-, beta-, delta-, epsilon-, gamma-, lenti-, and spumaretroviruses (Table S40). Using the custom Python (version 3.6+) script ERVin (https://github.com/strongles/ervin), we extracted all BLAST hits with an E-value $\leqslant 0.009$ that comprised a length $\geqslant 400$ amino acids for pol regions and $\geqslant 200$ amino acids for both gag and env regions. We grouped sequences according to their taxonomic relation to the first returned hit given by reciprocal BLAST. For the pol region, we extracted the highly conserved 200 amino acid region ending with a 'Y[M/V]DD' motif for all the bats and the 7 other mammals, and aligned them using MUSCLE within the Aliview software (v1.25)[43]. We manually inspected sequences and corrected the alignment. We discarded all sequences where the highly conserved region was shorter than 50 amino acids.

To reconstruct the phylogenetic tree of the retroviral pol like sequences for all 6 bat genomes and the viral probes used in BLAST search, we first ran Prottest (v3.4.2)[44] to determine which model to use with RAxML (version 8) package[45]. The best model was the VT+G model according to the AICc scoring criteria.

Our analysis showed that *M. molossus* displayed more gamma-like sequences for all 3 viral proteins in comparison to other bat species. Apart from that, we detected integrations for the following retroviral families: delta (*M. molossus*), epsilon (*P. kuhlii*), and spuma (*M. molossus* and *R. ferrumequinum*) for the pol region; lenti and epsilon for gag region (both in *P. kuhlii*); and alpha for the env region (*P. discolor*, *R. ferrumequinum*, *R. aegyptiacus*) (Fig. 4b). Overall, the highest number of integrations was observed in *M. myotis* (with the exception of the env region which was observed most frequently in the *M. molossus* genome), while the greatest variety of genera was observed in *P. kuhlii*. We also compared the numbers of pol, env, and gag regions found in bats to the 7 mammalian reference genomes (Fig. S13). Of all analysed genomes, *M. musculus* displayed the highest number of integrations of viral protein sequences. The numbers of pol sequences found in the non-bat mammalian genomes and 5 of our 6 bat genomes were comparable to each other, with *M. myotis*, whose genome contained twice more pol and gag integrations, being an exception. Apart from mouse, all of analysed bats exhibited more env and gag integrations in comparison to the other mammalian genomes.

#### 4. Genome evolution

##### 4.1 Identification and alignment of one-to-one orthologs across Placentalia

Human transcripts were projected to 41 additional mammal species (Table S1) using TOGA as described above. To avoid aligning non-homologous exons that belong to different transcripts, we selected a single representative transcript for each gene. Selection of the representative transcript was guided by the goal of selecting a transcript with an intact reading frame in our six bats to ensure properly aligned coding regions for these bats. To this end, we considered for each human gene all Principal APPRIS isoforms that were inferred to be 1:1 orthologs in any bat species. In the case where no or multiple Principal isoforms were determined, we considered the longest annotated transcript as a candidate. If this transcript did not contain an intact open reading frame (presence of internal stop codons in all three forward frames or >20% ambiguous bases (N's)) in all six bats, we discarded this transcript as a candidate for the representative transcript and replaced it with a functional alternative isoform where possible. The coding sequences of the final representative transcripts were then extracted from human and the 41 other species using the CESAR 2.0 mapping. Individual species were ignored if the representative transcript did not contain an intact reading frame.

To align coding sequences, we used the "alignSequences" module of MACSE (v2.01)[46], trimming potential non-homologous fragments from individual sequences using its "trimNonHomologousFragments" module. Sequences which contained an in-frame stop codon after alignment were removed. Alignments were retained if they contained at least one Yinpterochiroptera and one Yangochiroptera species. This resulted in a final set of 12,931 coding alignments, having a median coverage of 44 mammals.

**4.2 Phylogenetic inference and divergence time estimation**

The best-fit model of sequence evolution for each of the 12,931 nucleotide alignment files was determined using ModelFinder[47], which is part of IQ-TREE (v.1.6.10)[48], with species trees inferred using the maximum-likelihood (ML) method of phylogenetic reconstruction. A nucleotide supermatrix was generated by concatenating all 12,931 alignments into a single file, which was used as input to infer a mammalian species tree using IQ-TREE and using model partitions for each gene. Branch-support values were determined using UFBoot (v.2.0.0)[49] with 1000 bootstrap replicates. The tree was rooted with Atlantogenata (*Trichechus manatus*, *Loxodonta africana*, *Orycteropus afer*, *Echinops telfairi*, *Dasypus novemcinctus*) as a sister group to all other clades. This topology was then used to establish a time tree using r8s (v.1.81)[50] and the Langley-Fitch (LF) ML method with Truncated Newton (TN) optimization to find objective function optima. We constrained 14 nodes with fossil calibrations [51], as shown in Fig. S26. The final divergences time estimate of the last common ancestor of bats (63.38 Mya, Table S41) is similar to previous estimates (64 Mya in [52], 66.5 Mya in [53]). This time tree was used to infer gene/miRNA family expansion and contraction (see section 4.5 and 5.2.1).

In addition to coding sequences, the position of bats within Laurasiatheria was further investigated using 10,857 orthologous conserved non-coding elements (CNEs), using the aforementioned concatenation method. To explore how different genes may impact the tree space, we carried out topology tests that compare all 15 possible Laurasiatheria topologies to each individual protein-coding gene partition or CNE alignment and their concatenated supermatrices, using approximately unbiased (AU) tests[54] as implemented in IQ-TREE. The 15 possible Laurasiatheria topologies that all have Eulipotyphla constrained as basal, and Carnivora and Pholidota constrained as sister orders, are shown in Fig. S6. The number of protein-coding genes supporting each topology as the most likely tree ranged from 477 (Tree 9) to 1,007 (Tree 1), with 2,104 genes showing more than one topology as equally likely. Only 1,173 CNE alignments supported one unique topology (Table S10). The AU-tests of the protein-coding supermatrix and the 15 topologies rejected all but Tree 1, while the CNE supermatrix rejected all but Tree 1 and Tree 2.

Model misspecification due to an inadequate fit between phylogenetic data and the model of sequence evolution used can cause biases in phylogenetic estimates[55]. To assess whether model misspecification or loss of the historical signal [56] might have been a contributing factor to our phylogenetic estimate (Fig. 2a), we examined the 12,931 alignments of protein-coding genes for evidence of violating the assumption of evolution under homogeneous conditions (assumed by the phylogenetic methods used in this paper) and for evidence that the historical signal has decayed almost completely (due to multiple substitutions at the same sites). Either of these two cases imply that the data provided by such a gene may not be fit for phylogenetic analysis. To detect model misspecification and loss of historical signal, we used Homo 2.0 (https://github.com/lsjermiin/Homo2.0) and Saturation 1.0 (https://github.com/lsjermiin/SatuRation.v1.0), respectively. For each of the 12,931 protein-coding genes and each codon site within these genes (including unlinked 1st and 2nd codon sites), we surveyed the alignment, using the match-pairs test of symmetry[57] for evidence of violating the assumption of evolution under homogeneous condition. Likewise, these datasets were analysed for evidence of saturation of substitutions at variant sites.

A majority of the datasets were found to violate the phylogenetic assumption of evolution under homogeneous conditions (1st codon sites: 29.0%; 2nd codon sites: 13.5%; 3rd codon sites: 88.8%; 1st + 2nd codon sites: 41.7%; amino acids: 5.1%), implying that many of the datasets have evolved under more complex conditions than assumed by the models of sequence evolution used (note that concatenation of alignments does not mitigate the problem identified). The problem of loss of historical signal was less pronounced (1st codon sites: 4.4%; 2nd codon sites: 10.2%; 3rd codon sites: 6.1%; 1st + 2nd codon sites: 3.1%; amino acids: 3.4%). Based on these observations, and the requirement of having a sequence from all 48 species (many of the genes did not have a complete sequence for all species), we selected 1st + 2nd codon sites from 488 genes. A concatenation of these datasets was deemed fit for phylogenetic analysis assuming evolution under homogeneous conditions, and thus was subjected to the methods above.

Additionally, given these 488 genes were considered fit for phylogenetic analyses, we further explored the position of bats in Laurasiatheria under a model of coalescence using SVDquartets[58], as implemented in PAUP* (v.4.0b10, Swofford 2003), with 500 bootstrap pseudoreplicates. SVDquartets is a single-site coalescence method that is ideally applied to unlinked sites. However, this method also performs well with multigene alignments[59]. Importantly, SVDquartets avoids problems with the recombination ratchet and gene tree reconstruction error that negatively impact sequence-based coalescence analyses with gene trees[60-62]. The tree topology inferred under a coalescence model showed the same branching pattern for laurasiatherian orders as Tree 1, with bats as sister taxa to Fereuungulata. The position of Tupaia recovered in this topology (sister to Primates) is identical to the CNE topology (Fig. 2b), but differs from the concatenation topologies based on 12,931 protein-coding genes and 488 genes that fit model assumptions where Tupaia is sister to Glires (Fig. 2a and 2d).

##### 4.3 Selection test

###### 4.3.1 Genome-wide screen for signatures of positive selection

First, the aBSREL[63] model implemented in the Hyphy package (v2.3.11) was used to identify genes that have experienced episodic selection during the evolution of bats. For each alignment, we pruned the phylogeny, estimated from our amino acid supermatrix, to include only those species present in the gene alignment. All branches in the bat subtree were labelled as test branches. For each gene, aBSREL produces a corrected P-value if multiple branches are tested. This branch-corrected P-value was extracted for each branch tested. To account for the fact that our genome-wide screen for selection considered 12,931 genes, we further corrected the branch-corrected P-values by computing a false discovery rate (FDR) using the p.adjust tool and the Benjamini–Hochberg procedure in R (v3.3.1)[64], with an FDR cut-off of 0.05. We retained genes found to be under selection only at the bat ancestor and not elsewhere in the bat subtree. Second, the branch-site test for positive selection implemented in codeml from the PAML software suite (v.9.4)[65], was used to independently verify selection ($P$-value $< 0.05$) in genes identified under aBSREL and to identify putatively selected sites. To assure correctness of our homology statement, we manually inspected the alignment of all genes with significant evidence from aBSREL and codeml to detect obvious alignment errors. In addition to manual inspection, we used T_Coffee[66] to confirm a high quality of the entire alignment. Furthermore, we carefully inspected the neighbourhood of sites reported to be under selection and used T_Coffee to confirm a high alignment quality at these selected sites. For genes, where manual inspection or T_Coffee found putative alignment ambiguities, we produced a manually-adjusted alignment and re-ran aBSREL and codeml. We only reported selection in a gene if its manually-adjusted alignment also showed significant evidence for selection (aBSREL FDR < 0.05, codeml P-value < 0.05). For example, for the gene *TJP2*, a region of potential alignment ambiguity was identified during manual inspection. The alignment produced by MACSE produced significant evidence for positive selection (aBSREL P-value = $1.3 \times 10^{-7}$, FDR = 0.002), while the manually adjusted alignment lowered significance (aBSREL P-value = 0.009, FDR = 0.813). However, the manual adjustment revealed a possible echolocator specific insertion (Fig. S9), which is not considered as all insertions/deletions in general by phylogenetic tests for positive selection. All final alignments of genes with significant evidence for positive selection after manual curation are provided in Data S2.

##### 4.3.2 Candidate genes and selection tests

A list of human ageing-related genes was collated from GenAge[67]. To augment these ageing-related genes and identify genes associated with immunity and metabolism, we queried the Gene Ontology (GO) database, AmiGO[68], with 'ageing', 'immunity' and 'metabolism' as search terms. A total of 2,453 genes were investigated across all 6 bats, using the same alignments as for the genome-wide screen.

Each of the 2,453 gene alignments was analysed for signatures of positive selection using the branch-site test. All branch-site tests were carried out using codeml, inferring the likelihood-derived dN/dS (ω) values under both the null (ω1, ω2 constrained to be less than 1) and alternative (ω2 can vary) hypotheses. As branch-site tests require a species tree, analyses were carried out using the best-supported mammal topology, displayed in Fig. 2a, with the ancestral bat lineage designated as foreground branch. A likelihood ratio test (LRT = 2*(lnL alt – lnL null)), comparing the fit of both null and alternative log-likelihood values, was carried out for each alignment. P-values were then calculated assuming a chi-squared distribution[65] and corrected for multiple testing using FDR correction (p.adjust tool and the Benjamini–Hochberg procedure in R[64]). Only significant genes at an FDR cut-off of 0.05 having ω greater than 1 were considered for further interpretation. Sequence-specific sites undergoing positive selection were identified based on significant Bayes Empirical Bates (BEB) scores obtained from codeml (P-value > 0.95), and a subsequent visual inspection of alignments to rule out false-positive results due to potentially misaligned sequences. Significant genes showing ω values greater than 1, but with no identifiable BEB sites, were also reported (Table S14). Additionally, while five of the 15 genes showing significant P-values in HyPhy were included in the candidate list of 2,453 (*AZGP1*, *CXCL13*, *GLB1*, *LRP2*, *SERPINB6*; see 4.3.1), there were 10 genes that were not (*APOBEC3H*, *C17orf78*, *HP*, *INAVA*, *KBTBD11*, *NES*, *NPSR1*, *PALB2*, *TGM2*, *TRUB2*). These extra 10 genes were independently explored for selection using codeml, to investigate consilience between different methods. P-values from these 15 genes were corrected using FDR independently of those determined for the 2,453 genes (Fig. S10a).

A total of 22 out of the 2,453 genes relating to ageing, immunity and metabolism showed evidence of positive selection in the ancestral bat lineage using codeml in PAML (Table S15). Branch-site tests showed evidence of positive selection in the ancestral bat branch for genes associated with immune system modulation including both *IL17D* and *IL-1β*: cytokines playing roles in recruitment of natural killer cells to tumours[69] and the proinflammatory response, respectively. *IL-1β* has also been shown to up-regulate *DEFB1*[70], an antimicrobial defensin also significant in our analyses. Similarly, *CXCL13* connects innate and adaptive immune systems, promoting B-cell survival and maturation[71], and elevated levels are associated with autoimmune inflammation[72]. Positive selection was found in *SEMA4D*, which plays a role in the regulation of the humoral immune response[73]. It is also involved in the interaction between T-cells and antigen-presenting cells (APCs). These APCs can be activated by *TSLP*[74], another gene showing evidence of positive selection in the ancestral bat branch. Genes involved in the recognition and response to pathogens such as *GP2*, *MRC1*, *TLR9, LCN2*[75-77] also show evidence of bat-specific positive selection. Though not showing any specific adaptive sites, *PURB* had signatures of selection. In addition to a role in cell proliferation, *PURB* also regulates *MYC*[78,79], an oncogene shown to be under divergent selection in bats[80]. Selection was found in *NR1H2*, encoding the Liver X receptor *β* receptor. This receptor is activated by lipohilic ligands, such as oxysterols, binds to DNA and can interfere with the NF-kB signalling pathway, suppressing pro-inflammatory responses[81]. Additionally, *NR1H2* also regulates cholesterol transport and metabolism in the liver, thus demonstrating both immune and metabolic activity[82].

To further investigate overlap between both the aBSREL and codeml methods of selection analysis, P-values estimated for the subset of 2,453 genes, taken from genome-wide screen of 12,931 genes with the HyPhy suite of software, were FDR corrected and compared with results from codeml in PAML. A total of 14 out of the 22 genes showing signatures of selection with codeml overlapped with those significant using aBSREL (Fig. S10b), and included the aforementioned *LRP2*, *SERPINB6*, *IL17D*, *IL-1β*, *GP2, LCN2* and *PURB*. The remaining 8 genes showing significance with codeml that did not overlap had P-values less than 0.05 before FDR correction. Combining both genome-wide and candidate gene screen approaches to selection analyses has therefore identified robust signals of adaptive selection in the ancestral bat for key genes involved in both the function and regulation of immunity and the ability to tolerate various types of pathogens.

##### 4.4 Systematic screen for gene losses

To search for gene losses that occurred in the stem Chiroptera branch, we used a previously-developed approach to detect gene-inactivating mutations[83]. This approach uses whole genome alignments between a reference (here human hg38 assembly) and the six bat genomes presented here to detect large deletions that cover exons or entire genes, insertions and deletions that shift the reading frame, mutations that disrupt donor (GT/GC) or acceptor (AG) splice site dinucleotides, and mutations that create premature stop codons. To overcome issues related to genome assembly and alignment and evolutionary changes in the exon–intron structures of conserved genes, this approach performs a series of filter steps to exclude false inactivating mutations. Specifically, the approach (i) only considers those unaligning or deleted exons or genes where the respective locus does not overlap an assembly gap in the other genome, (ii) realigns all coding exons with CESAR, a Hidden Markov Model method that considers reading frame and splice site information to produce an intact exon alignment whenever possible[17,84], (iii) excludes alignments to paralogous genes or processed pseudogenes, and (iv) considers all principal or alternative APPRIS isoforms of a gene[27] and outputs the isoform with the smallest number of inactivating mutations. Our screen for lost genes is based on the human Ensembl v96 gene annotation[85]. The maximum proportion of the reading frame that remains intact for any transcript for each gene was also calculated.

To extract genes likely lost in stem Chiroptera, we filtered for genes for which less than 80% of the ORF is intact in all six bats. We excluded genes that are classified as lost in more than 20% of the non-Chiroptera Laurasiatherian mammals contained in our multiple genome alignment (https://bds.mpi-cbg.de/hillerlab/120MammalAlignment/Human120way/). While we confirmed previous findings that *PYHIN* genes (*MNDA*, *PYHIN1*, *IFI16*, *AIM2*) are completely deleted in Chiroptera[86] (Fig. S27), we excluded these genes in our screen because they were not intact in at least 80% of non-Chiroptera laurasiatherians. For the lost genes listed in Table S16, we manually inspected the genome alignment chains to confirm that the remnants of the lost gene are located in a context of conserved gene order and to rule out that a duplicated intact copy of these genes exist in bats.

##### 4.5 Protein Family Evolution

To investigate expansions and contractions of protein families, we used CAFE (v4.0)[87]. CAFE requires annotated proteins assigned to families. To this end, we downloaded GFF3 files from Ensembl which were available for 25 of the 42 considered species. To obtain a single isoform from each gene, we used genePredSingleCover (https://github.com/ENCODE-DCC/kentUtils.git) to obtain the longest transcript from each locus, and translated this transcript to a peptide sequence. The POrthoMCL pipeline (https://github.com/etabari/PorthoMCL, accessed 12/6/2019; commit dec8e5f), a parallel implementation of the OrthoMCL algorithm, was used to cluster proteins into families. All families were assigned PANTHER Database (v.14.0) IDs, based on the human genes contained in a family. Families assigned the same ID were merged. Where POrthoMCL families were composed of multiple families, all families with overlap based on PANTHER IDs were merged. To obtain families that were already present in the Placentalia root, we retained those families where at least one member was present in all bats, at least one of human, mouse or rat (representative Euarchontoglires) and one Atlantogenata species. Families which varied in size by more than 100 members between the species with the highest and lowest count were also removed. CAFE was then used to identify families which underwent expansion or contraction at the base of bats, using the previously produced ultrametric time tree (see section 4.2 above). The caferror.py function was used to estimate an error model for the data, which was used in further analysis. A single lambda, or birth/death parameter, was inferred for the entire tree. Families were retained if estimated to have undergone a significant expansion or contraction at the ancestor of all bats with an FDR value < 0.05.

##### 4.5.1 Evolution of the APOBEC3 gene cluster

Gene family PTHR13857 showed evidence of expansion along the ancestral bat lineage. In order to identify which family members had expanded in bats, a phylogenetic tree was constructed from all proteins assigned to PTHR13857. Protein sequences were aligned using the G-INS-i algorithm in MAFFT (v7.310)[88]. A phylogenetic tree was constructed using PhyML (v20120412)[89] using the BLOSUM62 substitution matrix, with 4 gamma distributed rate categories, and invariant sites. The APOBEC3 genes were found to be expanded within this family in bats. The APOBEC3 proteins from bats were classified into three classes based on the Z-domain, Z1, Z2 and Z3 using previously published motifs[90]. The Z2B motif, previously observed in Pteropid bats, was also used in classification[91]. Manual inspection of unclassified APOBEC3 proteins revealed small changes in the length of the linker region between functional residues and adjusting this allowed classifying these previously unclassified proteins. Finally, a Z1B motif observed in *P. kuhlii* (HxEx5xxx18-19SWSPCx2Cx6Fx8Lx5xxxx5-9Lx2Lx9M) was produced by modifying the canonical Z1 (HxEx5xxx18-19SWSPCx2Cx6Fx8Lx5RIYx9Lx2Lx9M). Those which remained unclassified were manually assigned to a class. All motifs used to classify APOBEC3 proteins are given in Fig. 3c. Proteins were also designated as likely non-functional if they did not contain a deoxycytidine deaminase domain motif, $HxEx_{24-33}PCxxC$. In order to understand the duplication history of the APOBEC3 proteins, the Z domains from all bat APOBEC3 proteins were aligned using MAFFT and a phylogeny constructed based on amino acid distance using BioNJ (Fig. S28).

### 5. Evolution of non-coding genomic regions

##### 5.1 Annotation of conserved non-coding RNA genes

In brief, the conserved non-coding RNA genes were annotated using the Infernal pipeline (v1.1.2)[92]. Initially, transposable elements (TEs) and low complexity DNA regions in each genome (six bat genomes plus seven additional mammalian genomes as outgroups; see Table S1) were hard-masked using RepeatMasker (v4.0.9) (http://www.repeatmasker.org) by aligning the genomic sequences against a custom library of known repeats (section 3.3). This library contains a collection of vertebrate TEs and the most up-to-date bat-specific TEs. It is noteworthy that some regions containing certain tRNA genes, small nuclear RNA (snRNA) genes and their pseudogenes were masked by RepeatMasker due to their high similarity to SINEs. The repeat-masked genomes were queried against the Rfam database (v14.0)[93] using Infernal (v1.1.2)[92] with default parameters. The alignments with an E-value < $10^{-6}$ were considered statistically significant and their corresponding genomic regions were annotated as conserved non-coding RNA genes. Based on the Rfam database, these candidates were further categorized into ribosomal RNA (rRNA), small nuclear RNA (snRNA), small nucleolar RNA (snoRNA), microRNA (miRNA), and long non-coding RNA (LncRNA). Other RNA types or uncharacterized RNA genes were grouped into miscellaneous RNA (miscRNA). See Fig. 5a for the summary of non-coding RNAs in six bats. The number of conserved ncRNAs that are shared between bats is shown in Fig. S15.

##### 5.2 The evolution of conserved miRNA gene families

To gain an overview of the evolutionary patterns of conserved miRNA families along the bat lineages, we performed two analyses that investigate (i) expansions or contractions of members with miRNA gene families (section 5.2.1) and (i) gains or losses of miRNA families (section 5.2.2). These analyses compared 48 mammalian taxa (6 bat species plus 42 non-bat taxa; see Table S1). miRNA gene families of the 42 non-bat taxa were predicted using the same pipeline as described above, and the copy number for each miRNA family was subsequently determined. This pipeline reduced the number of false-positive miRNA predictions, which overlapped with annotated TEs, to a minimal level. We obtained a matrix containing the copy numbers of 286 conserved miRNA families across 48 mammalian species. This dataset was filtered by retaining those miRNA families present at least in one Atlantogenata species and one of the rest mammalian species investigated. As the current miRNA set is biased towards conserved and highly expressed miRNA, the lineage-specific miRNAs could not be discovered via *ab initio* genomic prediction, therefore, were not included in these analyses.

###### 5.2.1 miRNA family expansion and contraction

miRNA family expansion and contraction analysis was carried out using CAFE (v4.2.1)[87]. A random birth and death model was used to infer the evolution of miRNA gene copy number across a user-specified phylogenetic tree. We used the supermatrix tree that was inferred on the basis of the alignments of 12,931 single-copy orthologous genes across 48 taxa (section 4.1) and calibrated as described in section 4.2. An error model was estimated to correct for genome assembly error. The global parameters lambda and lambdamu, which indicated miRNA birth and death rates across all branches for all gene families, were separately estimated using maximum likelihood. A P-value was calculated for each family. The miRNA families with an FDR value $< 0.05$ were regarded to have a significantly accelerated expansion and contraction rate. The genomic loci of the miRNA families exhibiting expansions and contractions in bat lineages were manually checked and confirmed.

###### 5.2.2 miRNA gene gain and loss

The gain and loss of miRNA families was inferred by using Dollop from the Phylip software (v3.696) (http://evolution.genetics.washington.edu/phylip/doc/dollop.html). Dollop is based on the Dollo parsimony principle, which assumes an independent evolution in each lineage and irreversibility of gene loss. In the context of this study, it implies that once a miRNA gene family is predicted to be lost in certain lineages, it cannot be regained during evolution. For Dollo inference, the supermatrix tree (section 4.2) and a binary matrix derived from the matrix used for CAFE analysis (section 5.2.1) were employed. In this binary matrix, '1' and '0' indicate the presence and absence of each miRNA family in each of the 48 taxa, respectively. The number of miRNA family gain and loss in each branch and node was further extracted using in-house Perl scripts.

To assess the performance of the Dollo parsimony principle, we generated a random matrix of phylogenetic profiles where both miRNA family presence in each species and the phylogenetic tree were shuffled. Based on this matrix, the number of losses required to explain the random profiles was determined by Dollop. We observed a major difference in the number of inferred miRNA losses between real and random data. In particular, the random data resulted in multiple losses, while the real data could be generally explained by a limited number of losses (Fig. S17). This result supports the Dollo assumption that the evolutionary patterns of most miRNA gene families can be inferred by a single acquisition event.

###### 5.2.3 Single-copy miRNA alignments across 48 mammals

To ascertain the sequence conservation of these predicted miRNA families between bats and other mammals, we focused on the single-copy miRNA genes across 48 taxa. We only considered the single-copy miRNA genes that were present in at least 80% of all taxa and at least 3 bat species. Based on these criteria, 98 single-copy miRNA genes were investigated and their precursor sequences in each genome were retrieved using Bedtools (v2.25.0)[94], respectively. For each miRNA gene, the precursor sequences were aligned using ClustalW (v2.1)[95] and the alignments were visualized in Geneious (v7.1.9) (https://www.geneious.com). For each miRNA, conservation of the mature 5' and 3' sequences and the hairpin loop was further investigated and manually curated (Table S42).

**5.3 Novel microRNAs that evolved in bats**

**5.3.1 Small RNA Illumina sequencing from brain, kidney and liver**

To identify novel microRNA evolved in bats, we Illumina sequenced small RNA libraries from brain, kidney and liver for all 6 bat species (Table S23). Briefly, total RNA was extracted from respective tissue types using TRIzol reagents (No. 15596-018, Carlsbad, CA) following the manufacturer's instructions. The quality and quantity of RNA were measured using a Bioanalyzer 2100 (Aligent Technologies). The samples with total RNA > 1 µg and an RNA integrity score (RIN) > 7 were prioritized for Illumina small RNA library preparation. RNA libraries were prepared using the Illumina TruSeq small RNA library preparation kit and were further sequenced on Illumina HiSeq 4000 platforms at the BGI (Hong Kong). Each sample was sequenced to a minimum depth of 30 million 50 bp single-end reads. The information of small RNA sequencing is summarized in Table S23.

**5.3.2 miRNA profiling pipeline**

Our approach to identify novel miRNAs was based largely on miRDeep2 (v2.0.0.8)[96]. Prior to analyses, the 3' adaptor sequence (5'-TGGAATTCTCGGGTGCCAAGGAACTCCAA-3') and low-quality bases (< Q25) were trimmed from the raw reads using Cutadapt (v1.14). We further filtered the reads with low complexity and only retained the reads ranging from 16 bp to 25 bp in length. Subsequently, identical reads were compressed to single entries with the headers indicating their read counts using Mapper[96]. These unique sequence tags were then mapped to the respective genome and were analysed by miRDeep2 to predict mature miRNAs and their precursor sequences. The prediction was based on the stability of their secondary structures and their sequence similarity to the known miRNA curated in miRBase (release 22)[97]. We considered miRNA candidates, which had the read counts < 5 and the true positive probability < 60%, as unreliable and excluded them from downstream analyses. miRDeep2 categorized miRNA into known and novel groups. We further manually inspected the novel groups by comparing them against the miRBase (release 22). For each bat species, any miRNA in the novel category, which shared the same seed region (nucleotides 2-7 of the mature miRNA sequence) with a known miRNA, were moved to the respective known groups. This filter ensures that the miRNAs in novel groups have a novel target specificity and potentially a novel repertoire of gene targets.

**5.3.3 Identification of known and novel miRNAs in each bat species**

To identify known miRNA in each bat species, the raw reads from brain, kidney and liver were first pooled together and the pipeline described above was employed. This pipeline resulted in two categories of predicted miRNAs: known and novel miRNAs. The miRNAs in the known group also exist in other mammalian species in miRBase. In general, novel miRNAs are usually expressed at a lower level than known miRNAs[98], which makes it more likely that authentic novel miRNA are falsely regarded as sequencing noise. To resolve this, we also predicted novel miRNAs for each bat using small RNA-seq data from each individual tissue (brain, kidney and liver). In this second screen, we considered miRNA candidates assigned to the novel group, if their expression was detected in at least two of the three profiled tissues. To achieve this, for each species all novel precursor miRNA predicted from brain, kidney and liver were pooled, and the sequence similarity was calculated using CD-HIT (v4.6.7)[38]. Only miRNAs, whose precursor sequences showed > 95% identity between tissues, were considered as reliable novel miRNA. The number of known and novel miRNAs identified in each species is listed in Table S23. For each species we further analysed the genomic coordinates of both known and novel miRNA based on gene annotation using Bedtools (v2.27.0)[94]. The distribution of miRNA locations in exons, introns, 3'UTRs and intergenic regions is given in Fig. S29.

##### 5.3.4 miRNA evolution in bats

To better understand miRNA evolution in bats, we investigated and compared novel miRNAs that evolved in our 6 bat species. miRNAs are often expressed in a time- and tissue-specific manner, which implies that small RNA-seq data may not capture all novel miRNA that are shared among all 6 bats. Therefore, to identify novel miRNAs that are likely shared among all 6 bats but are not present in any other of the 42 mammals, we integrated our small RNA-seq data and sequence similarity searches. Briefly, we first merged all novel miRNA precursors predicted from small RNA-seq data of our bats, and removed redundancy using CD-HIT (v4.6.7)[38]. Next, to identify shared miRNAs by sequence similarity, we mapped these nonredundant novel miRNA precursors to the 6 bat genomes and the other 42 mammalian genomes using bowtie (v2.2.5)[99] with the -N 1 parameter to allow at most one mismatch in the alignment seed. We allowed a maximum of 2 mismatches or indels outside the seed region and required an identical seed sequence. The number of novel miRNA and novel seeds shared across 6 bats was plotted in Fig. S20. Next, we only kept miRNA precursors that were successfully mapped to all 6 bat genomes, but did not map to any of the other 42 mammalian genomes. Subsequently, the filtered miRNA precursors were compared against the NCBI nucleotide database[41] and miRBase (release 22). We excluded any miRNA that exhibited homology to non-bat genomic sequences (NCBI nt database) and or that exhibited homology to non-bat miRNAs. This approach resulted in 12 novel miRNAs identified in all 6 bats. Details of these 12 novel miRNAs, including precursor and mature sequences, seed regions, expression values in different tissues, and hairpin structures, are listed in Table S24.

##### 5.3.5 3'UTR and miRNA target prediction

Our analysis of 3'UTRs inferred from Iso-seq data showed that many genes in each bat species had alternative 3'UTRs (Table S21). In order to obtain a comprehensive set of 3'UTRs that maximizes the potential target space for miRNA target prediction, we generated a "pseudo 3'UTR" for each gene per species, defining the pseudo 3'UTR of a gene as the union of all its annotated 3' UTRs. To do this, we used Bedtools (v2.27.0) to merge the coordinates of alternative 3'UTRs for genes with more than one annotated 3'UTR where possible or concatenated different 3'UTRs with 20 'X's if they did not share overlapping coordinates using in-house scripts. For the few cases, where the 3'UTRs of neighbouring genes overlapped in their genomic coordinates, we first separated the 3'UTRs of each gene and processed them separately. The statistics of pseudo 3'UTRs is summarized in Table S21.

We found that miR-337-3p has a unique seed region in bats compared to all other 42 taxa (Fig. 4c). To investigate whether this unique seed alters the predicted target genes, we developed a pipeline to extrapolate different targets of miR-337-3p between bats and human. As miR-337-3p mature sequences are conserved among all bats, we created a 'master list' of 3'UTRs by merging the pseudo 3'UTRs from the 6 bats to predict target genes. This procedure produced a set of 13,083 genes that could be potentially regulated by miR-337-3p in all 6 bats. We used the mature miR-337-3p sequence to predict targets in the master list of 3'UTRs using both miranda (v3.3a)[100] and RNAhybrid (v2.2.1)[101]. For miranda, we determined the optimal minimum free energy (MFE) cut-off by employing empirical data (real miRNA – target gene pairs predicted by miranda)[100] and plotting their distribution. As shown in Fig. S30, we observed a wide range of MFE values and chose -10 kcal/mol as the cut-off. For RNAhybrid, no empirical data is available, therefore we used the default cut-off of -20 kcal/mol. To increase the reliability of target prediction, we only kept target genes that were predicted by both methods or target genes that were predicted in multiple species ($n > 1$) by only by one method. To predict miR-337-3p targets in human, we extracted 3'UTR sequences from the same 13,083 genes in the human genome (hg38) that had corresponding 'pseudo' 3'UTRs found in 6 bats (the 'master' list). This allowed us to compare predicted targets in 3'UTRs of the same set of genes in bats and humans, which is a requirement to test whether the differences in the miR-337-3p seed region alters the set of predicted target genes. Targets of human miR-337-3p were predicted using the same procedure as described above.

GO enrichment analysis was performed using DAVID[102]. The non-redundant list of target genes predicted above was used as a query list while the list of the non-redundant genes that had 3'UTR data supported by Iso-seq was used as the background list. Enrichment analysis was performed on the first sublevel GO terms for Biological Process (BP), Cellular Component (CC) and Molecular Function (MF), using Fisher's exact tests. Enriched GO terms with a P-value $< 0.05$ after correcting for multiple testing using the Benjamini-Hochberg method were considered statistically significant.

##### 5.4 Functional validation of novel miRNAs and their regulatory gene targets

Luciferase assays to test the functionality of miRNAs were designed and performed as described previously[103,104]. The precursor sequences predicted by miRDeep2 were cloned in the pLKO.1 vector (Invitrogen) carrying the flanking sequences representing the primary transcript of the hsa-miR-342, which allowed optimal transcription; we did this in order to ensure transcription from a known and reliably expressed miRNA. All insertions were confirmed by Sanger sequencing. To maximize the sensitivity of the assay, we designed the miRNA sensors to contain two copies of the ideal targets of the to-be-tested miRNA. To this end, we inserted two repetitions of a fully complementary sequence to the cognate miRNA within the 3'UTR of the firefly luciferase gene in the pmiR-GLO vector (Promega). All cloning oligonucleotides are listed in Table S20. Luciferase assays were performed in HEK293 cells as described in [103,104]. Briefly, 50ng of miRNA expression vector and 70ng of sensor vector were combined and transfected in HEK293 cells (18K cells per well in 96well plate format, density $5.625 \times 10^5$ cells per $cm^2$) using GeneJuice (Merck Millipore) transfection reagent. 48 h post-transfection, firefly luciferase and renilla luciferase activities were measured as per manufacturer's instructions (Dual Luciferase reporter assay system, Promega), using a fully automated plate reader (TECAN, F200PRO or TECAN MPLEX, both equipped with fully automated injectors). Ratios between the firefly/renilla luciferase activity were normalized to account for technical variability.

**Fig. S1: Missing UCE.157 in some of mammalian genomes.** UCE.157 is not fully present in the assemblies of cow, cat and dog because of assembly gaps. UCSC genome browser screenshot shows a multiple genome alignment of mammals of the locus around UCE.157 (highlighted) and pairwise chains of co-linear alignments (blocks represent local alignments; double lines represent unaligned sequence and single lines represent deletions). The top-level pairwise alignment chains between human (reference species) and cow, cat, and dog show that UCE.157 only partially aligns (cow, dog) or does not align at all (cat). The unaligned region overlaps an assembly gap in all three cases, suggesting that the UCE sequence is not present because of assembly incompleteness. In support that the UCE is present in other mammals, the alignment chains of close-related sister species (zebu cattle, leopard, dingo) show that these assemblies completely contain UCE.157.

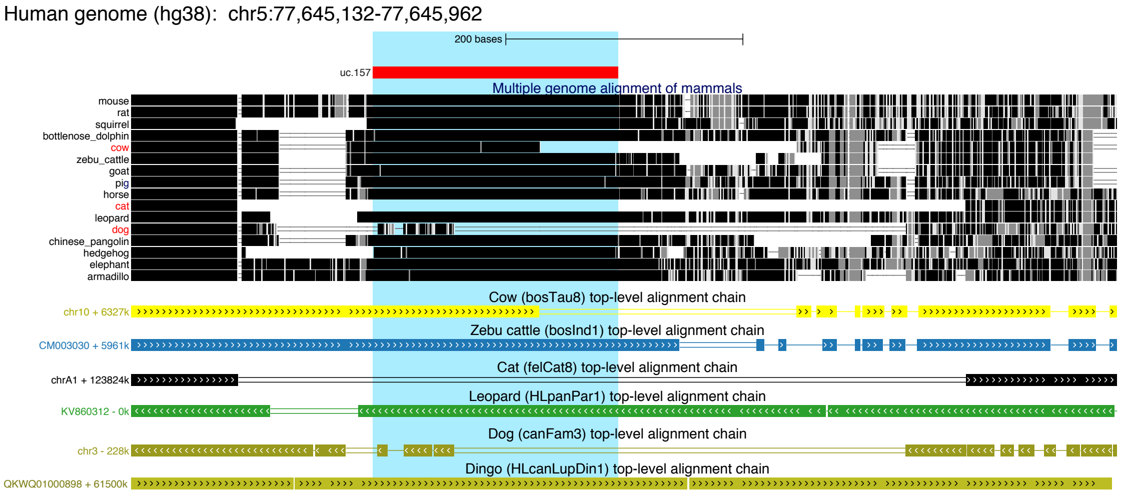

**Fig. S2**: **Alignment of bat UCE.47 sequences showing sequence divergence in *Myotis* and *Pipistrellus* bats**. Dots in the alignment represent nucleotides that are identical to the human sequence shown at the top. While *R. aegyptiacus*, *R. ferrumequinum*, *P. discolor*, and *M. molossus* have few sequence changes compared to human, *M. myotis* and *P. kuhlii* show numerous mutations. Importantly, many of these mutations are shared between *M. myotis* and *P. kuhlii*, indicating that these mutations already arose early in Vespertilionid lineage. Supporting this, mutations are also shared with related Vespertilionid species. Shared mutations also show that the sequence divergence is real and not attributed to base errors in the assemblies.

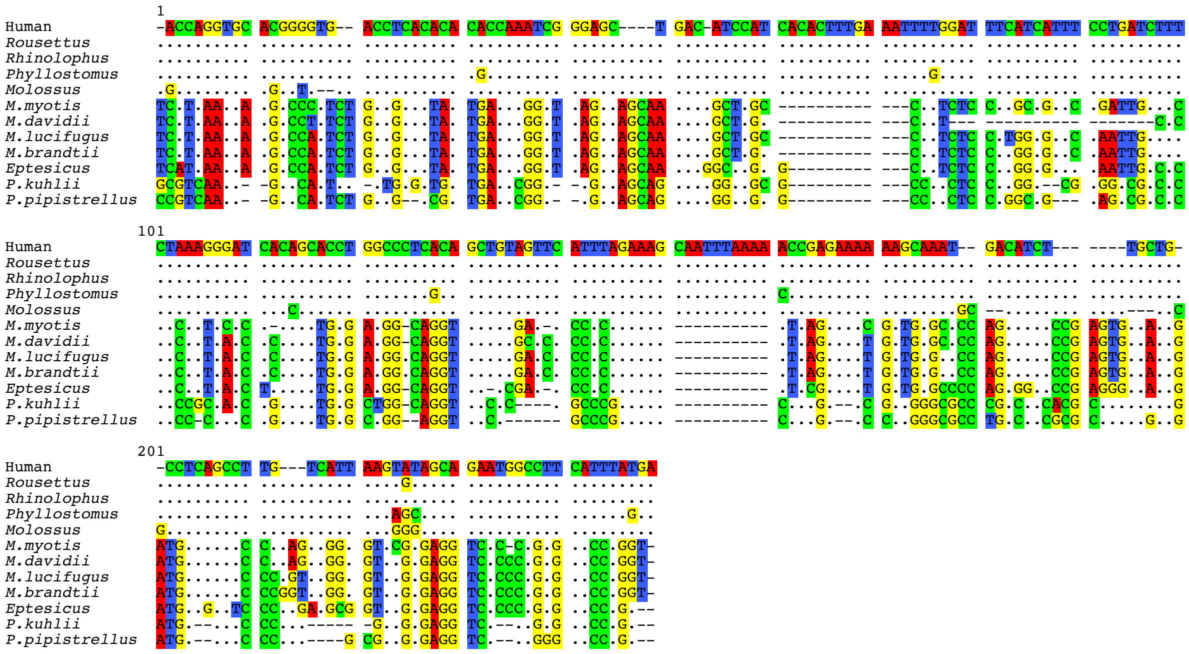

**Fig. S3: Genetic distance of Vespertilionid UCE.47 to other bats and canonical human sequence.** The heatmap shows that the UCE.47 sequences of *Rousettus*, *Rhinolophus*, *Phyllostomus*, *Molossus* and human are highly conserved. In contrast, the sequence of Vespertilionid bats, represented by *Myotis*, *Eptesicus* and *Pipistrellus* species, is substantially diverged from the conserved UCE sequence but relatively similar within the clade.

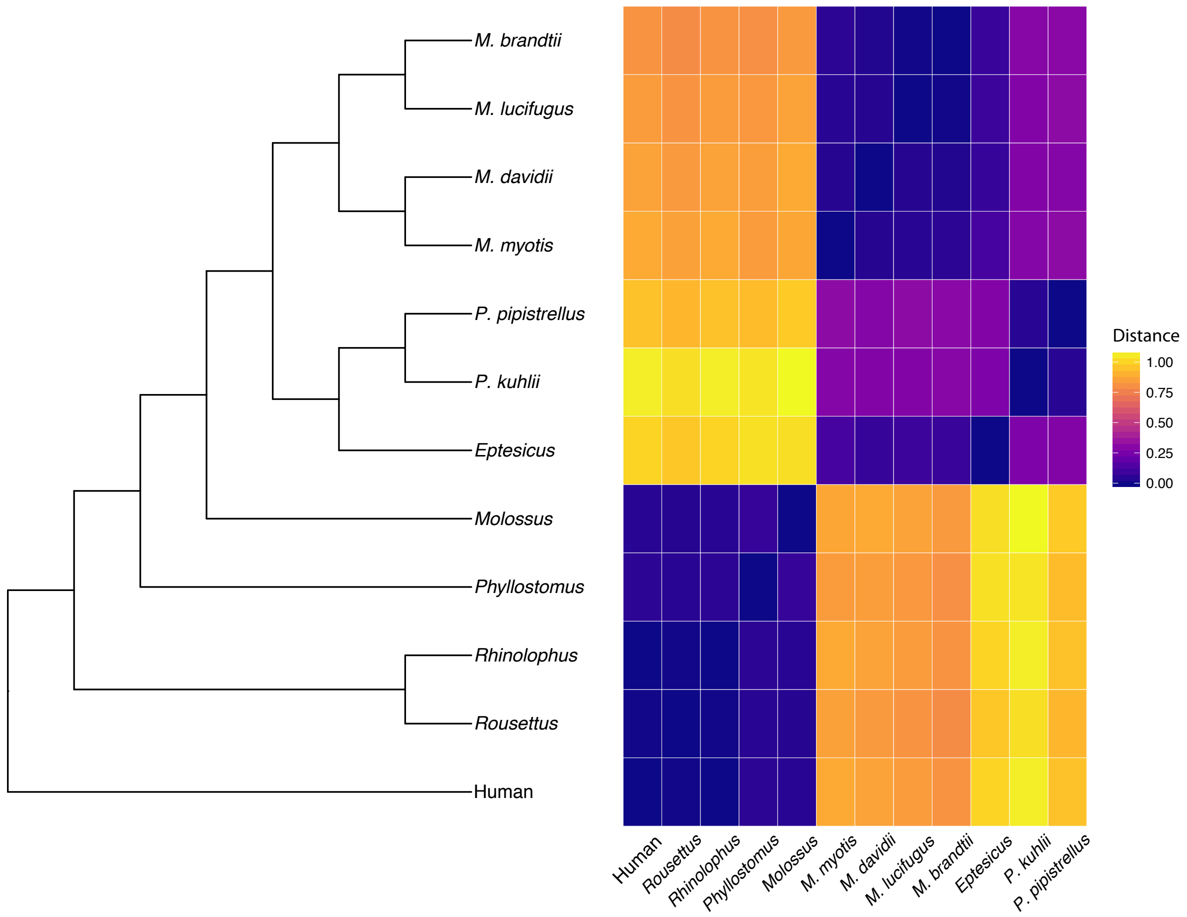

**Fig. S4: Alignment of bat UCE.394 sequences showing sequence divergence in *Pipistrellus* bats.** Dots in the alignment represent nucleotides that are identical to the human sequence shown at the top. Compared to other bats, *P. kuhlii* shows an increased number of mutations in this UCE sequence. These mutations supported by Illumina reads (note that reads do not cover the entire UCE locus). Furthermore, most mutations are shared with *P. pipistrellus* (which has two nearly identical loci in the current genome assembly), indicating that the sequence divergence is real and not attributed to base errors in the assembly.

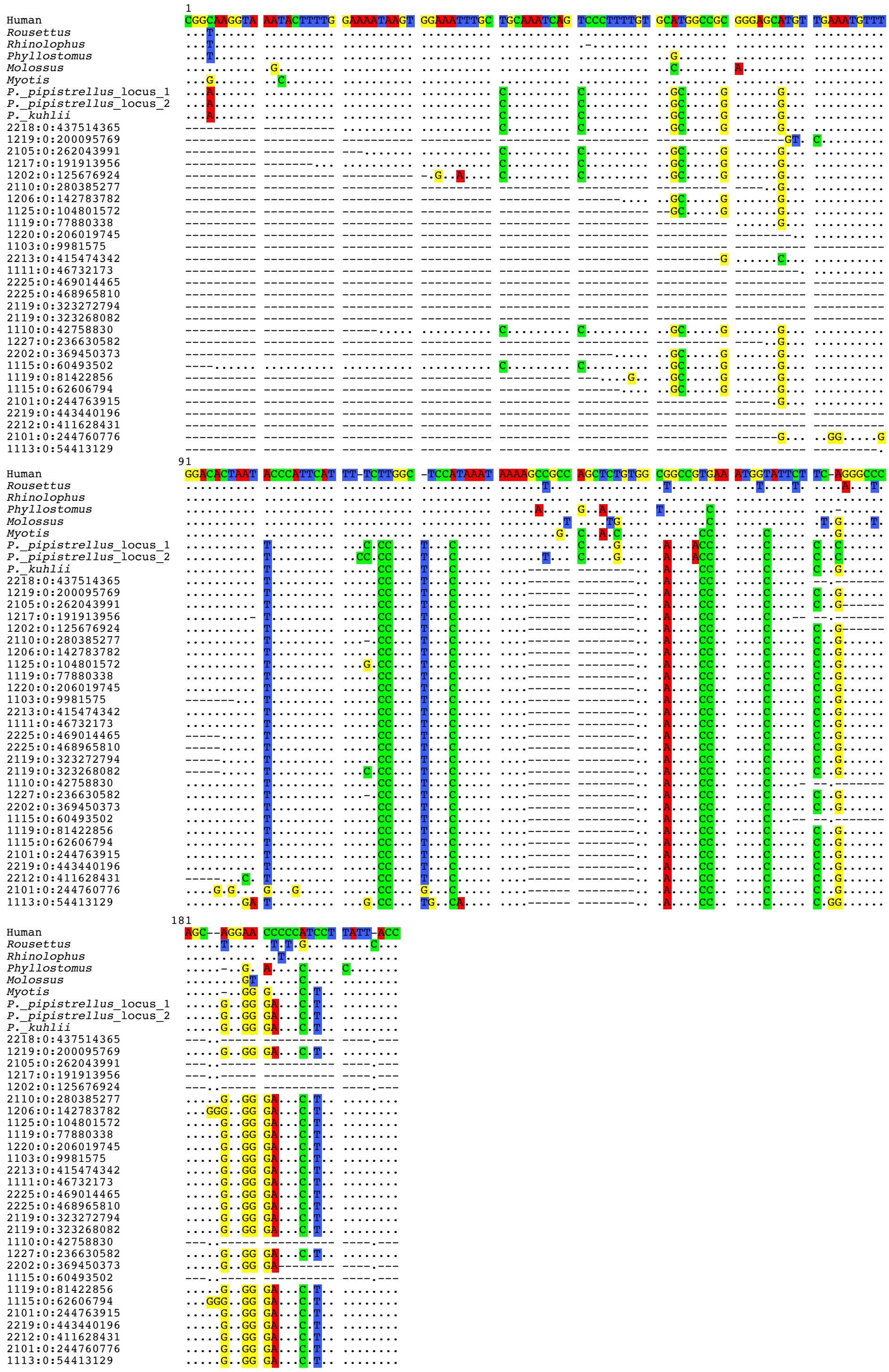

**Fig. S5: Alignment of bat UCE.446 sequences showing sequence divergence in *Pipistrellus* bats.** Dots in the alignment represent nucleotides that are identical to the human sequence shown at the top. Compared to other bats, *P. kuhlii* shows an increased number of mutations in this UCE sequence; however, it should be noted that *M. myotis* also shows an increased number of mutations. Since most mutations are shared between *P. kuhlii* and *P. pipistrellus*, base errors in the assembly are highly unlikely to account for the increased sequence divergence.

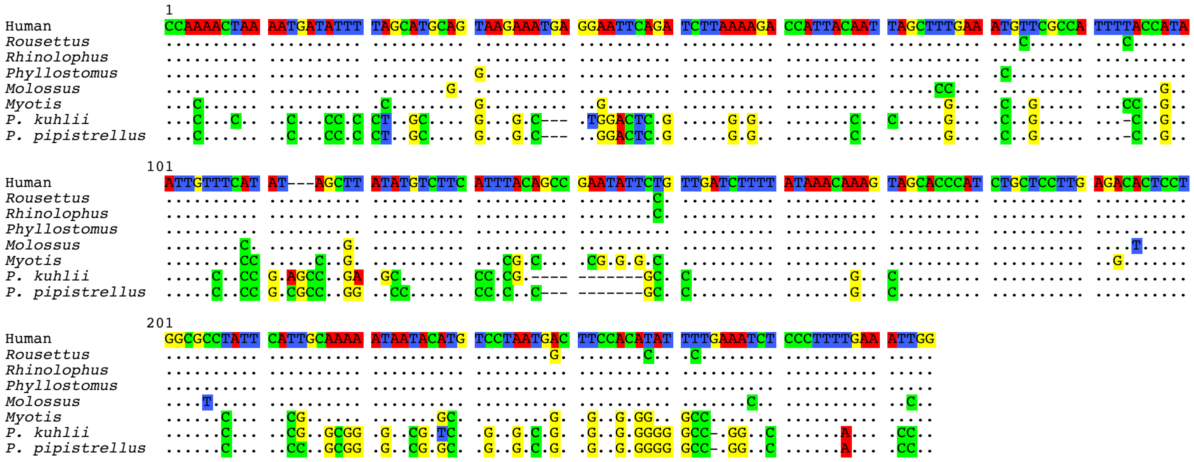

**Fig. S6**: **The 15 topologies showing different arrangements of Laurasiatheria and the number of gene trees supporting them with the highest likelihood.**

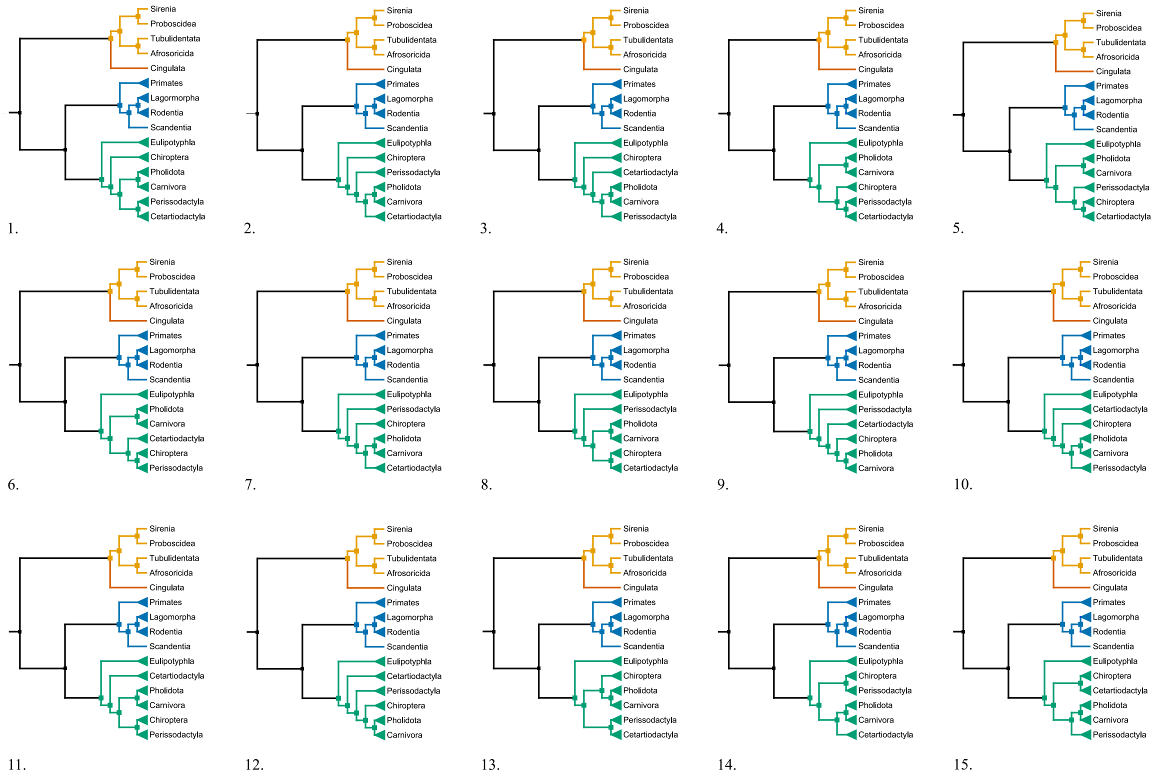

**Fig. S7: Tree topology using coalescence methodologies.** Using the 488 genes considered fit for phylogenetic analyses, the position of baths within Laurasiatheria under a model of coalescence using SVDquartets. The resulting phylogeny is displayed. The tree is rooted on Atlantogenata, with support values from bootstrap pseudoreplicates. Only nodes with support less than 100 have their values displayed.

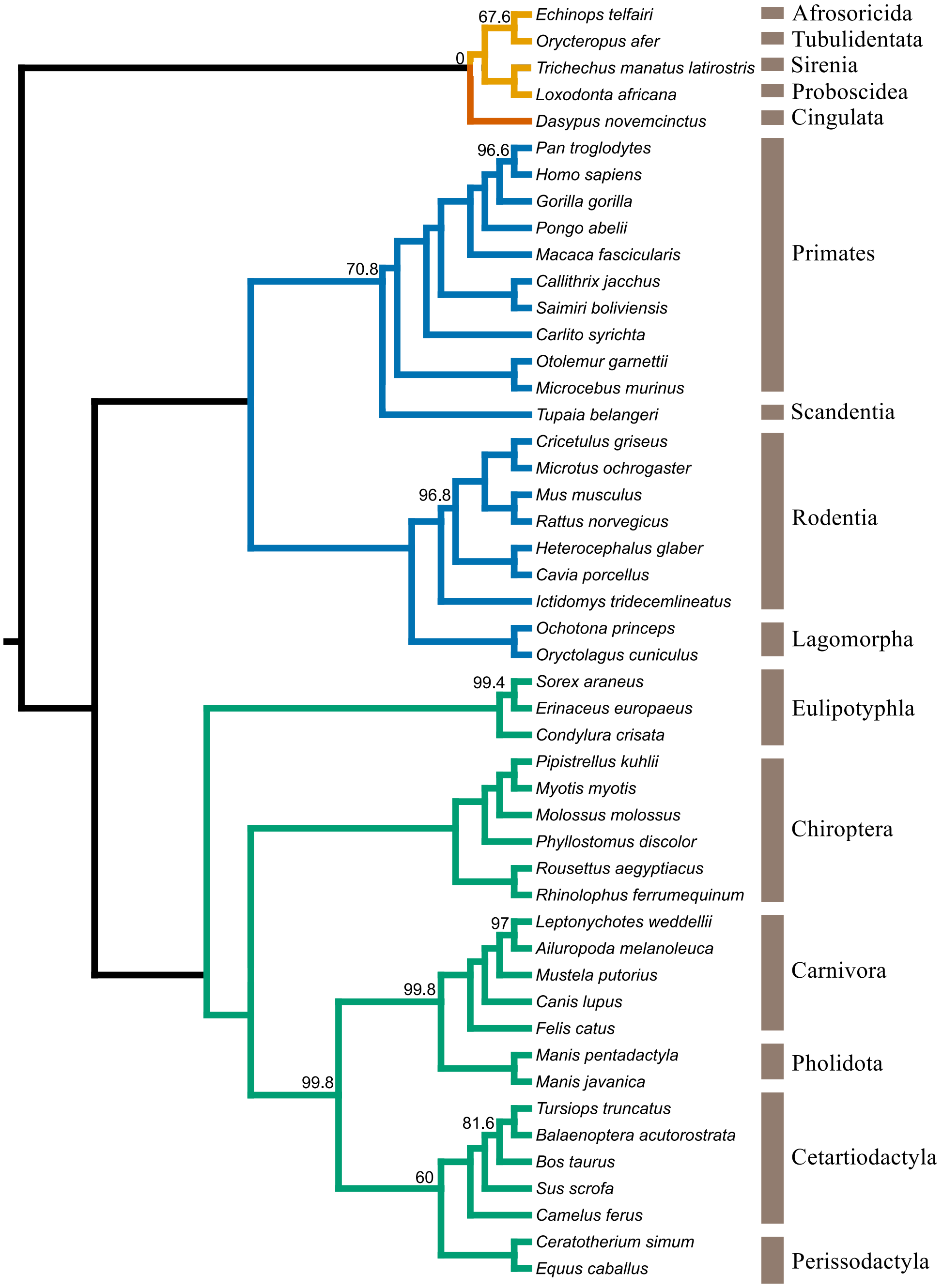

**Fig. S8: LRP2 sites under positive selection in bats.** Multiple sequence alignments of local regions surrounding 2 bat specific mutations which were found to under positive selection (BEB > 0.95) using codeml (PAML). Site 1564 shows bat specific changes at a conserved residue. The paraphyletic echolocating bats (indicated by a red dot) all share a methionine at this site, while the pteropodid, which do not use laryngeal echolocation, have a threonine at this site. Site 2540 shows a bat specific change, shared by all bats. The presence and patterns of mutations found in our six bats were confirmed in 6 previously published bat genomes, to increase taxonomic representation. Human (*Homo*), cow (*Bos*) and dog (*Canis*) are also shown.

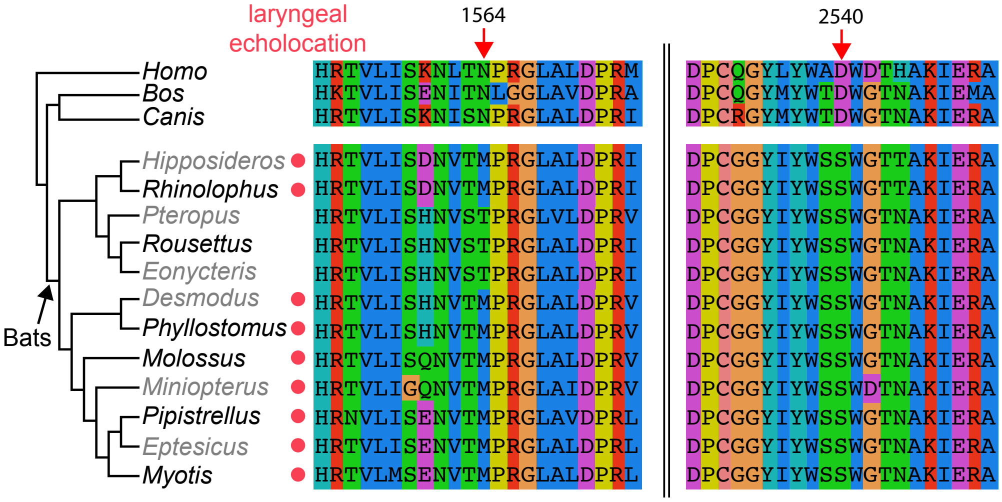

**Fig. S9: Alignment ambiguity in TJP2 gene.** Multiple sequence alignments of local regions surrounding bat specific mutations which were found to under positive selection (BEB > 0.95) using codeml (PAML), indicated by red arrows. The paraphyletic echolocating bats are indicated by a red dot. The alignment at left depicts two bat specific indels, produced by manual adjustment. Echolocating bats share an insertion not present in the pteropodid bats, which lack laryngeal echolocation. This insertion is adjacent to a bat specific deletion. An alternative alignment, produced by MACSE, shows a single bat deletion with two putatively selected sites. The presence and patterns of mutations found in our six bats were confirmed in 6 previously published bat genomes, to increase taxonomic representation. Human (*Homo*), cow (*Bos*) and dog (*Canis*) are also shown.

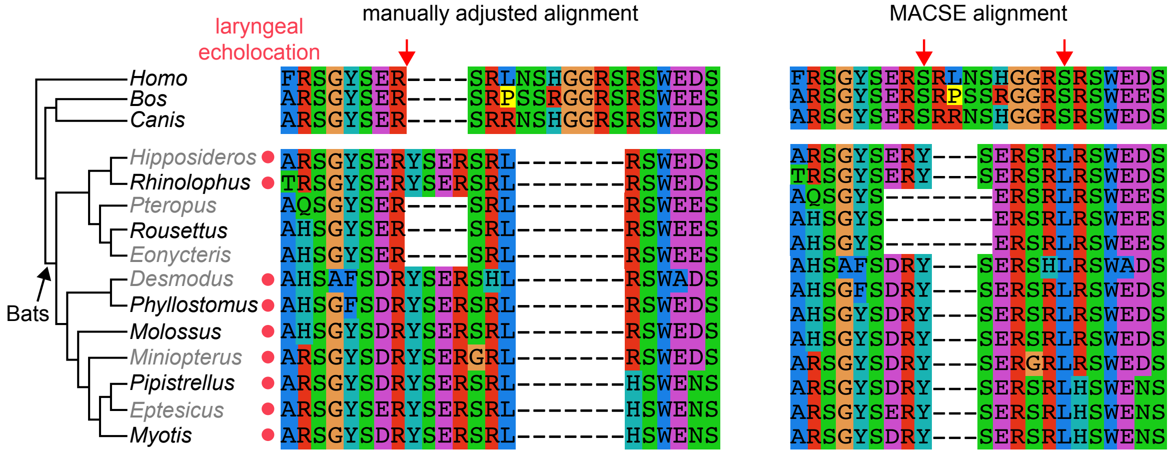

**Fig. S10: Overlapping genes between aBSREL and codeml. (a)** Genes showing evidence of significant positive selection across both aBSREL (12,931 genes) and codeml (2,453 genes) are shown. Genes not in the candidate 2,453 genes that were significant in HyPhy and subsequently validated with codeml are designated by '*'. **(b)** Genes showing evidence of significant positive selection in both aBSREL (HyPhy) and codeml (PAML) are displayed. Results are based on comparing corrected P-values from the 2,453 candidate genes with their respective results from the genome-wide scan.

(a)

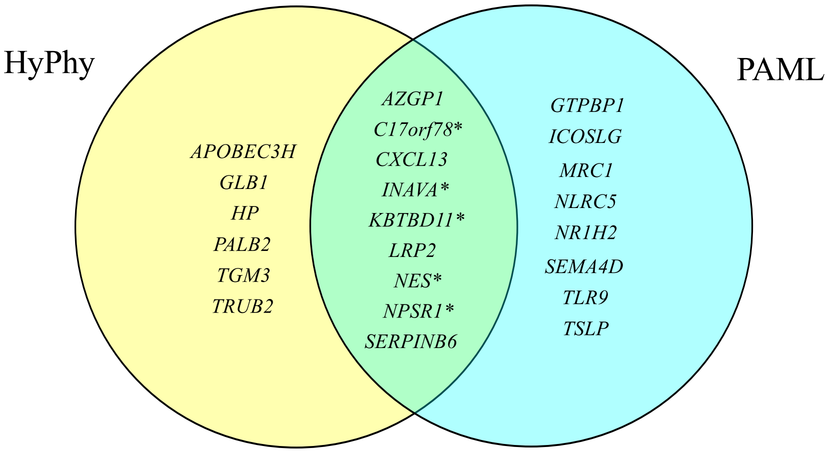


(b)

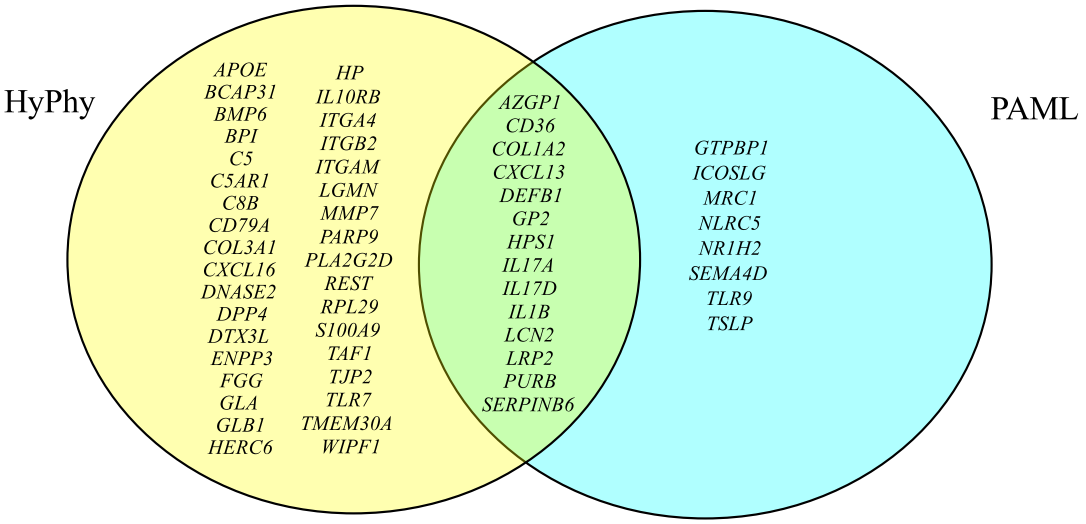

**Fig. S11: Inactivating mutations in LRRC70 in bats.** The single coding exon of *LRRC70* in each of the six bats is represented as a box. All detected inactivating mutations are visualized. Vertical red lines show frameshifting deletions, arrow heads indicate frameshifting insertions. The size of deletions or insertions is given on top of the mutation. Premature stop codons are indicated by black vertical lines and the corresponding triplet. Mutated ATG start codons are indicated as 'noATG'. One representative mutation for each bat is shown in detail in an alignment between human and bats (red font indicates the inactivating mutation). Genome assemblies produced in this study are in black, publicly available assemblies of sister species are in grey font. It should be noted that the position of the -4 bp frameshifting deletion in *Pipistrellus* and *Myotis* is ambiguous and can be shifted by up to 3 bp to the right without affecting alignment identity. Importantly, the presence of the exact same mutation in independently sequenced and assembled genomes of sister species excludes the possibility that the representative mutations are erroneous.

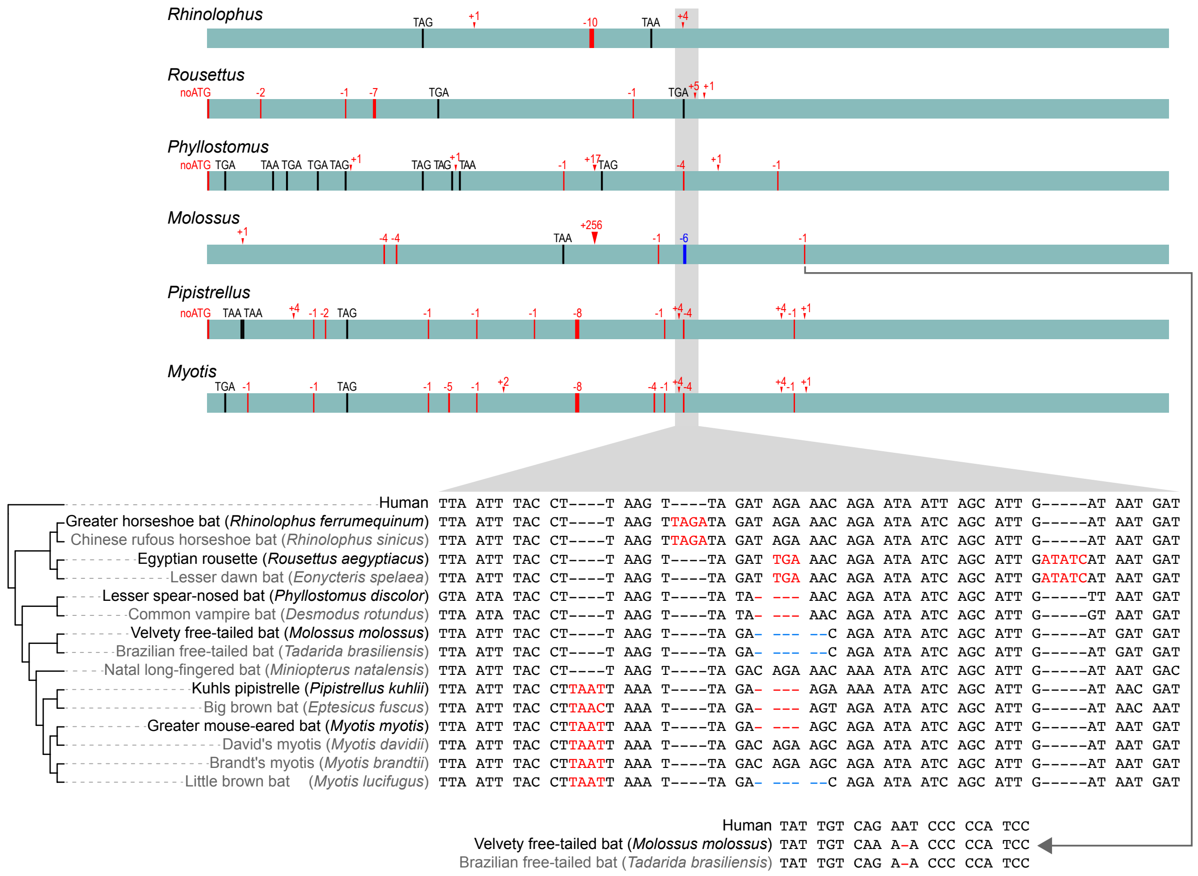

**Fig. S12: Inactivating mutations in IL36G in bats.** The four coding exons of *IL36G* are represented as boxes. All detected inactivating mutations are visualized. Vertical red lines show frameshifting deletions, arrow heads indicate frameshifting insertions. The size of deletions or insertions is given on top of the mutation. Premature stop codons are indicated by black vertical lines and the corresponding triplet. Mutated ATG start codons are indicated as 'noATG'. Splice site mutations are shown by red letters at the end of an exon (donor mutation) or the beginning of an exon (acceptor mutation). One representative mutation for each bat is shown in detail in the alignment between human and bats (red font indicates the inactivating mutation). Genome assemblies produced in this study are in black, publicly available assemblies of sister species are in grey font. As for *LRRC70*, the presence of the exact same mutation in sister species genomes excludes the possibility that such mutations are erroneous.

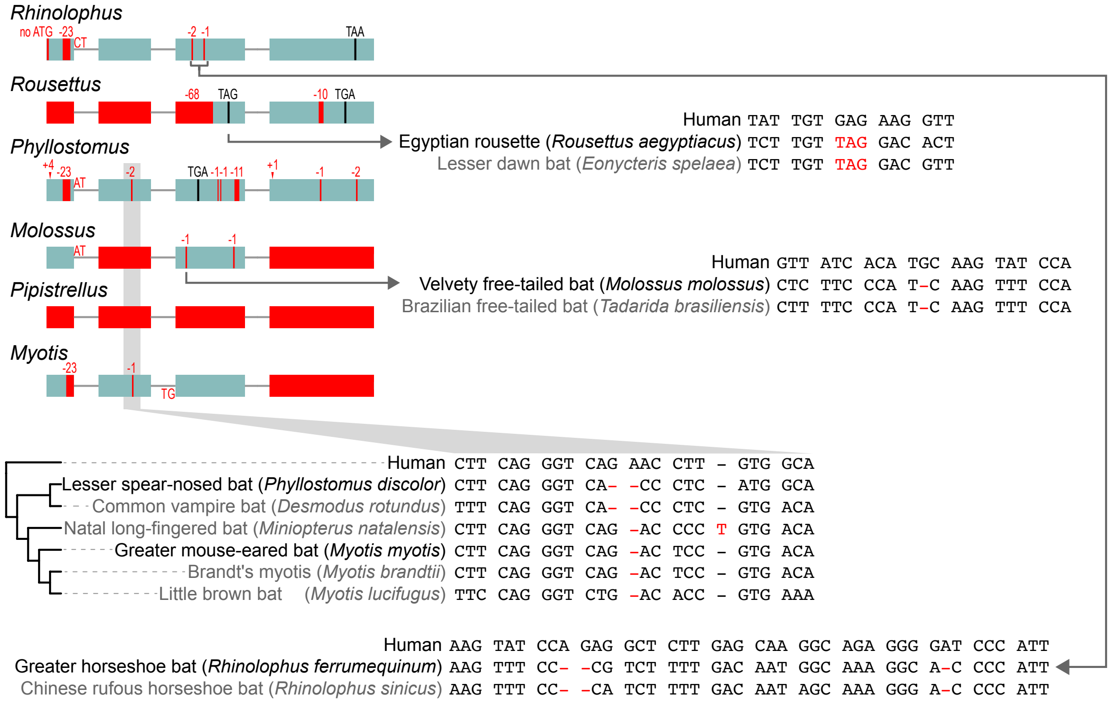

**Fig. S13: Mosaic plot showing the relative numbers of viral gene sequences found in six bat genomes compared to seven mammalian reference genomes.**

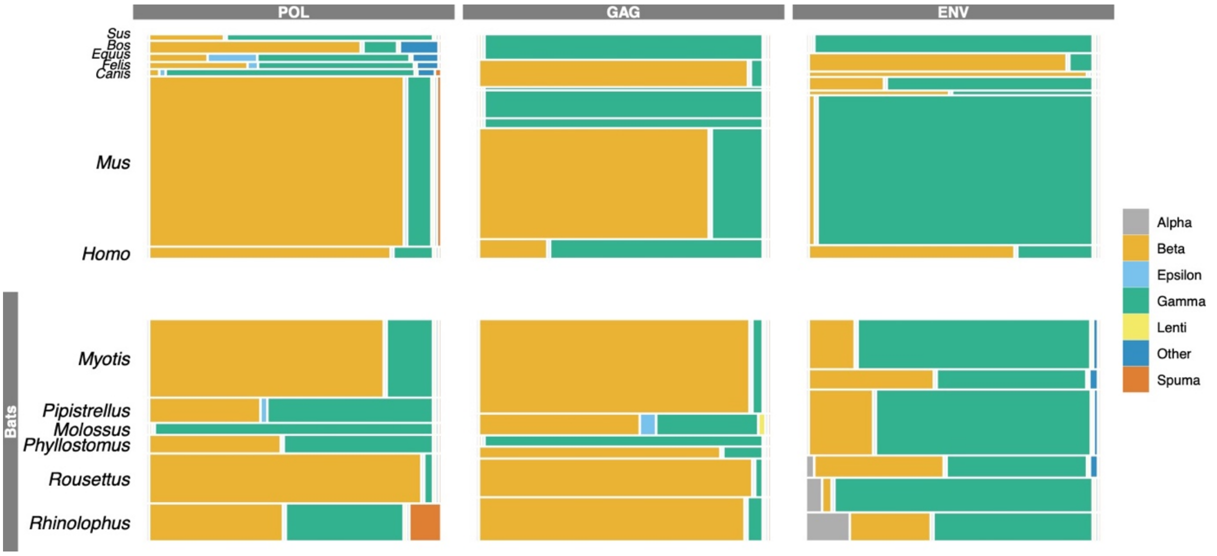

**Fig. S14: Reconstructed phylogenetic tree of viral pol sequences found in six species of bats.** Each species used in the ERV search is marked with a colour: *Phyllostomus* (navy blue), *Myotis* (green), *Pipistrellus* (orange), *Rhinolophus* (yellow), *Molossus* (light blue), *Rousettus* (pink) and reference sequences (black). Bootstrap values are shown where the values are ≥ 70%. The tips of the phylogeny are labelled with the species name, position in the reference genomes and the direction of the sequences (N- negative/P- positive).
**(See Separate File)**

**Fig. S15: The number of conserved noncoding RNA genes shared across 6 bat species.** The figure was generated using UpSetR[105]. Each black dot indicates each species. The connected black dots indicate the number of noncoding RNA genes shared between them.

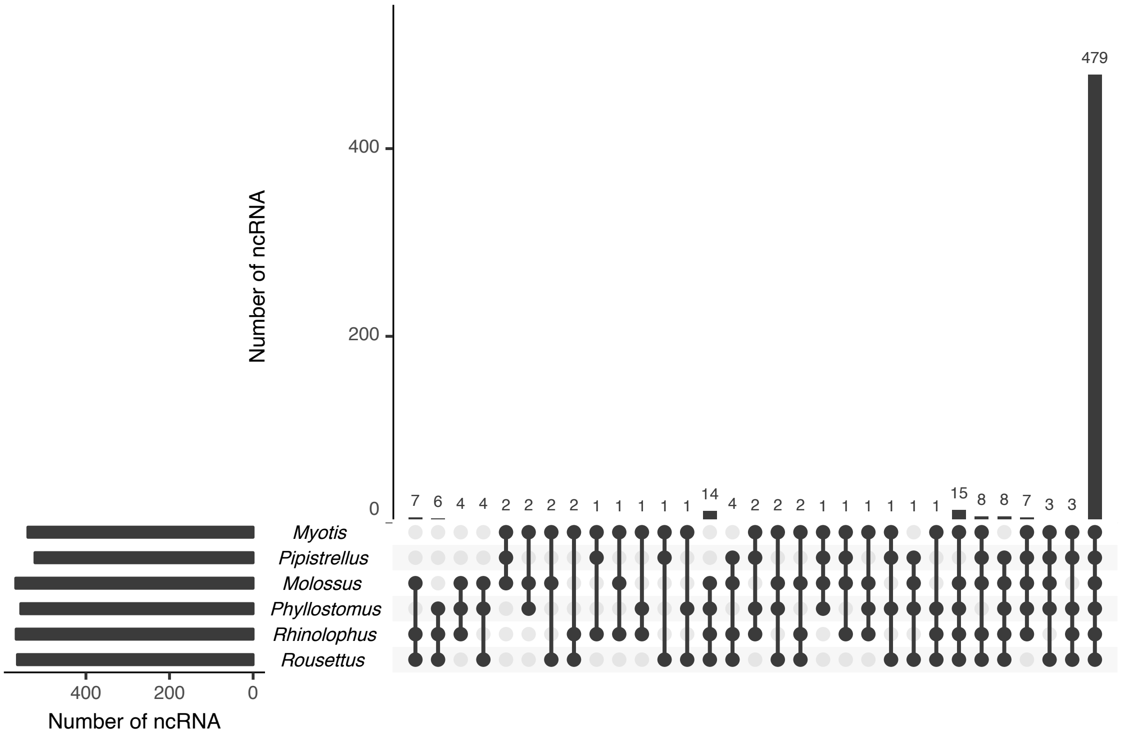

**Fig. S16: The number of miRNA gene families under contraction and expansion along the lineages.** The phylogenetic tree was inferred based on the alignment of 12,931 protein-coding genes. Based on 286 conserved miRNA gene families, the number of miRNA families under a significant rate of contraction and expansion was inferred by CAFE. The red values indicate the number of expanded miRNA families while the blue values indicate the number of contracted miRNA families. The bat species were highlighted in purple.

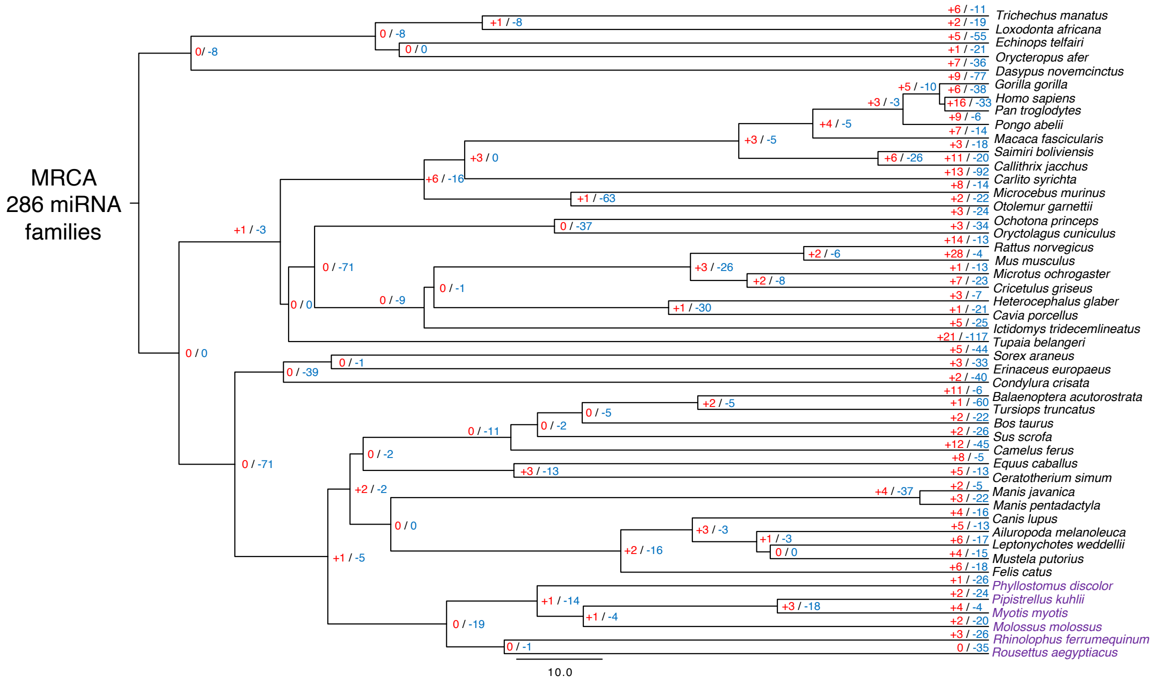

**Fig. S17: The performance of the Dollo parsimony principle on random and real data.** The real data refer to the observed phylogenetic tree inferred from the alignment of 12,931 protein-coding genes and the matrix containing the number of miRNA copies across 48 mammalian species; while the random data refer to the relationship of species in the phylogenetic tree and the number of miRNA copies that have been randomly shuffled. The figure indicates the number of losses required to explain the observed phylogenetic pattern, both on real and random data.

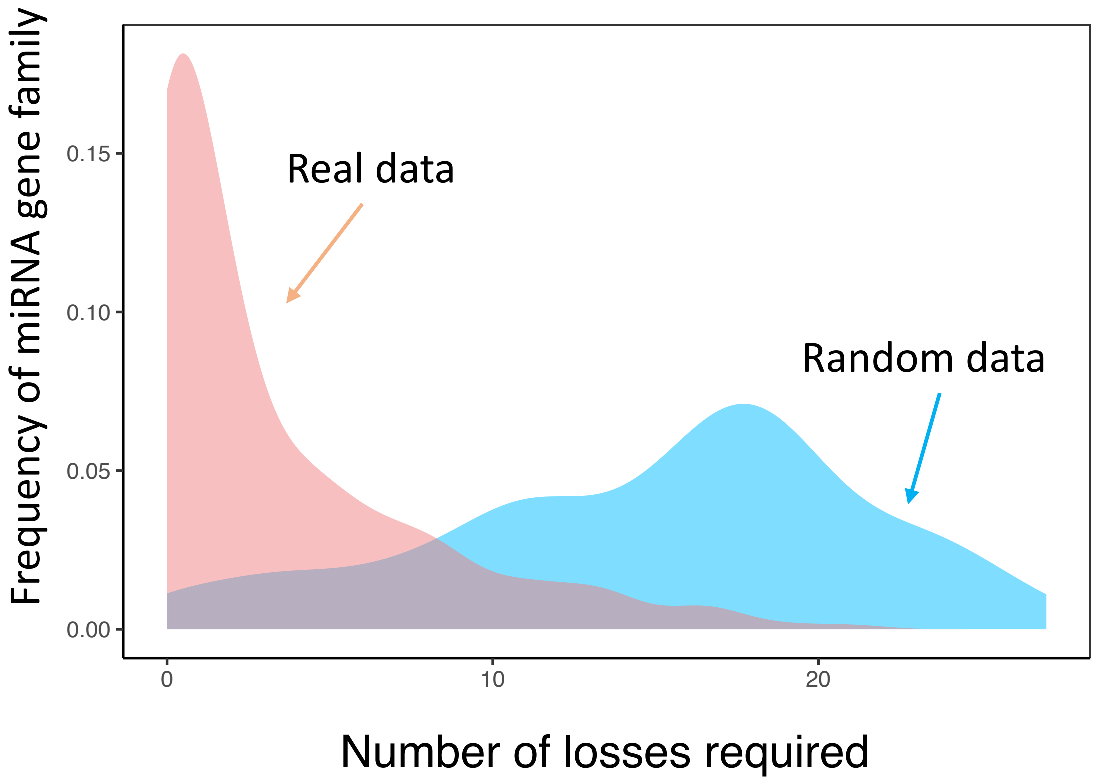

**Fig. S18: The miRNA gene loss and acquisition based on the Dollo parsimony principle**. The red numbers indicate miRNA gain while the blue numbers indicate miRNA loss. The bat species were highlighted in purple.

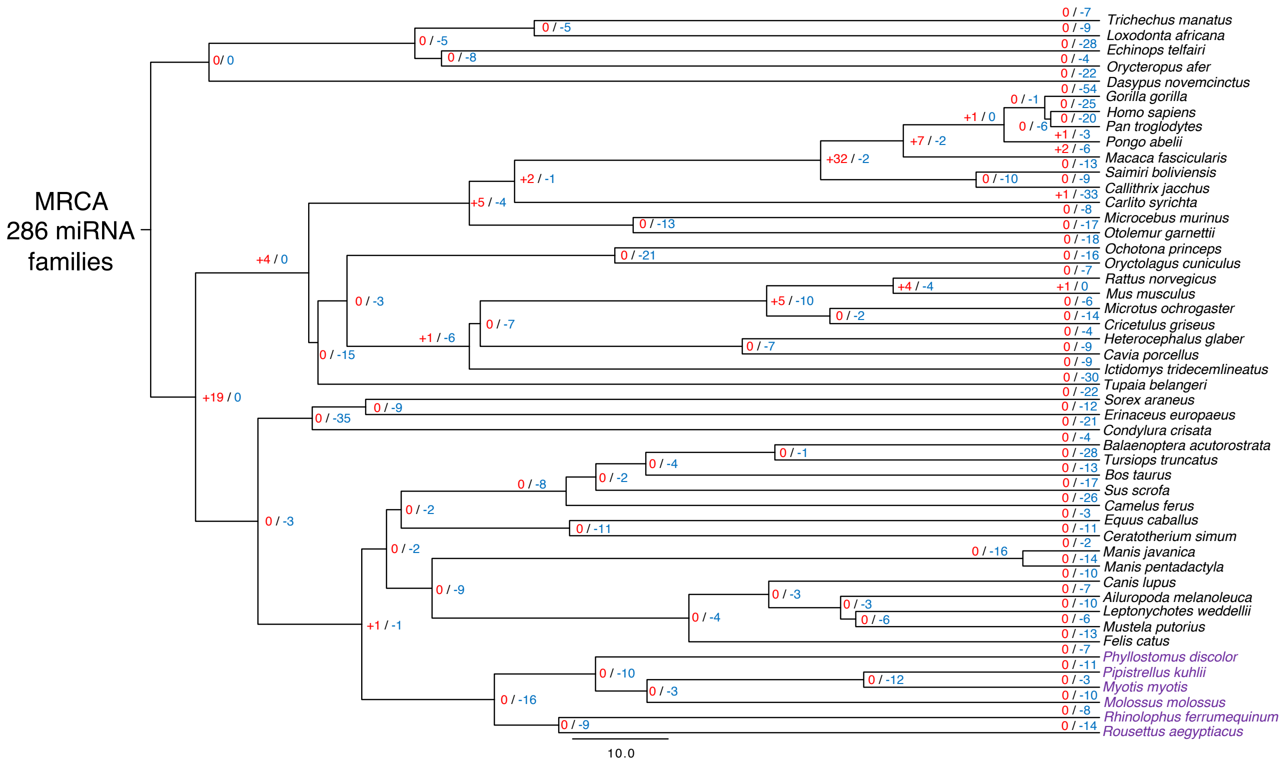

**Fig. S19: The expression (RPM) of miR-337 in brain, liver and kidney in 6 bat species based on the miRNA-Seq data.**

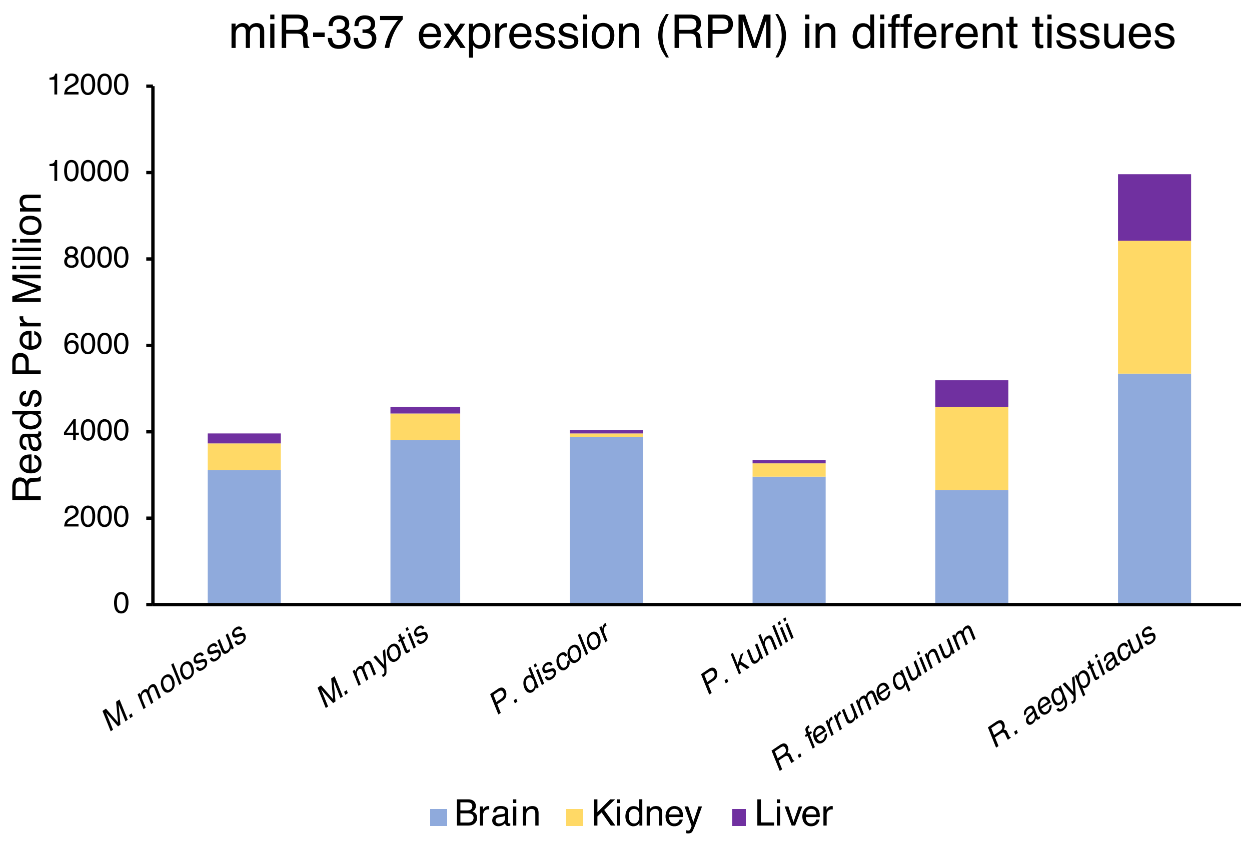

**Fig. S20: The evolution of novel miRNA predicted in 6 bat species. a)** The number of novel mature miRNA shared across 6 bat species. **b)** The number of novel seeds shared across 6 bat species.

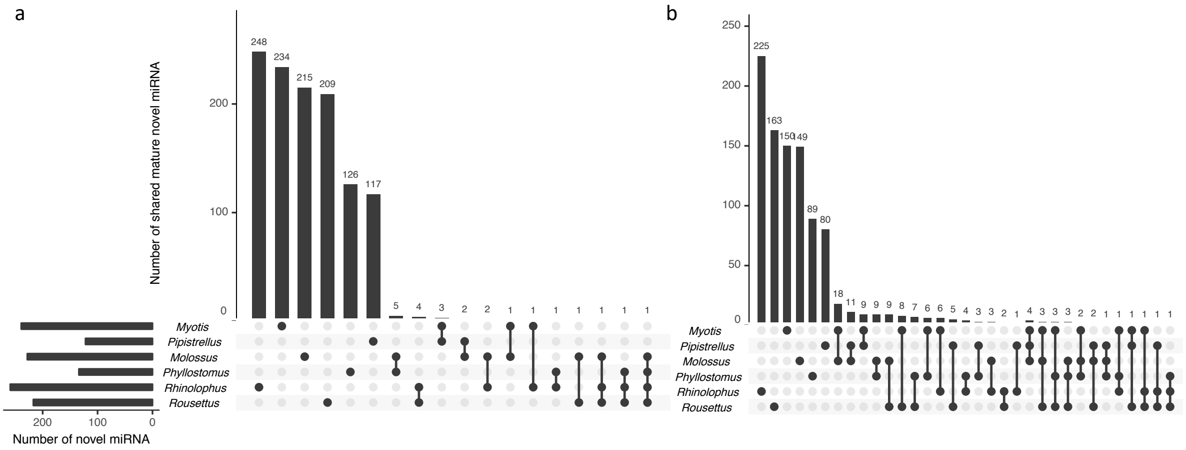

**Fig. S21: Distribution of PacBio read length (N1 to N100).** The dashed line at 4 kb marks the minimum read length that was used in the assemblies.

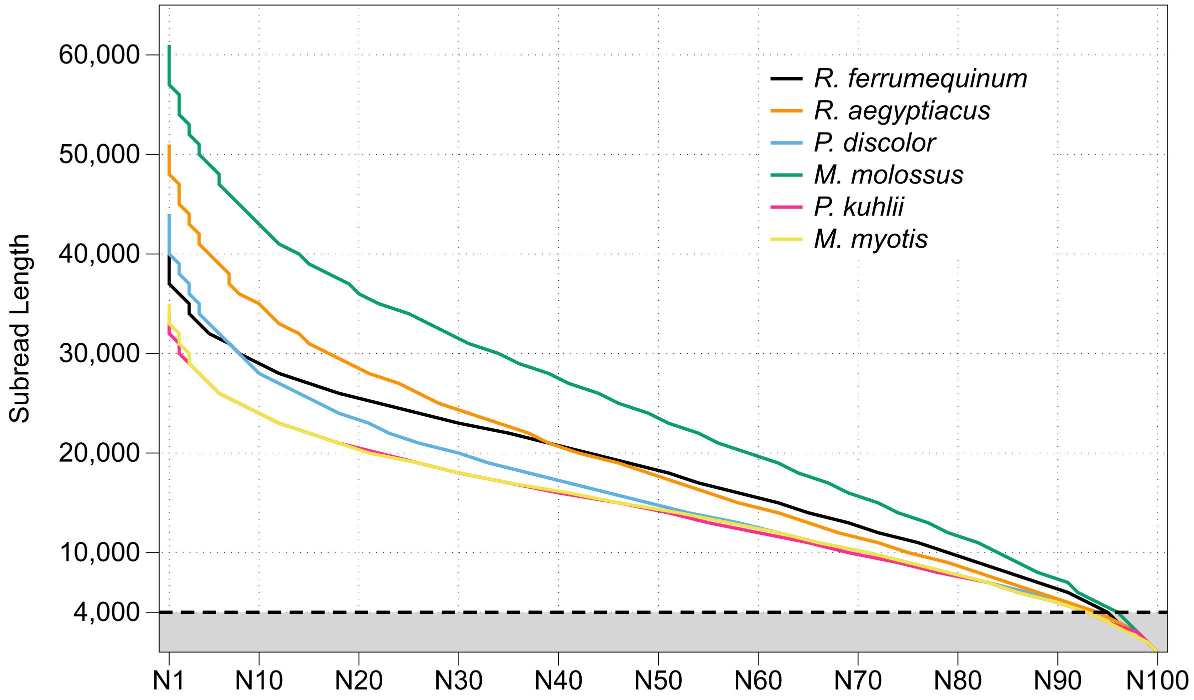

**Fig. S22: Distribution of putative 10x molecule lengths (N1 to N100).** Based on the final assemblies, 10x reads were mapped with longranger align and the molecule lengths were calculated using the tool bxcheck (https://github.com/pd3/bxcheck).

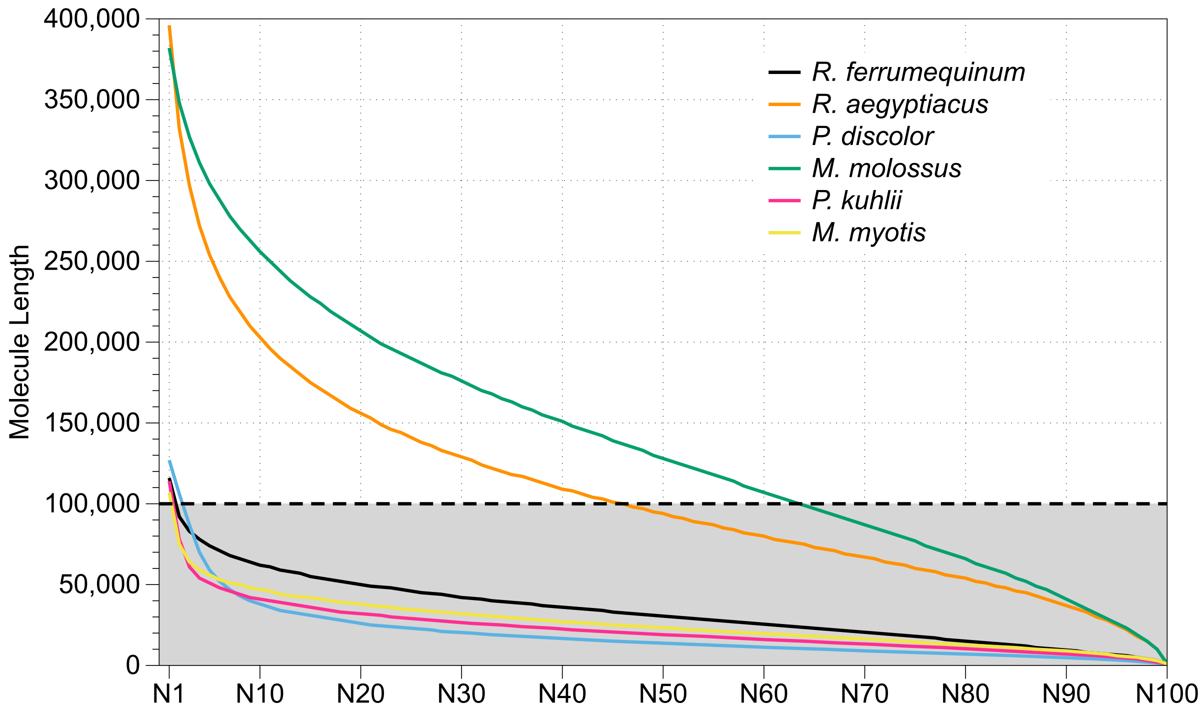

**Fig. S23: Bat1K assembly pipeline.**

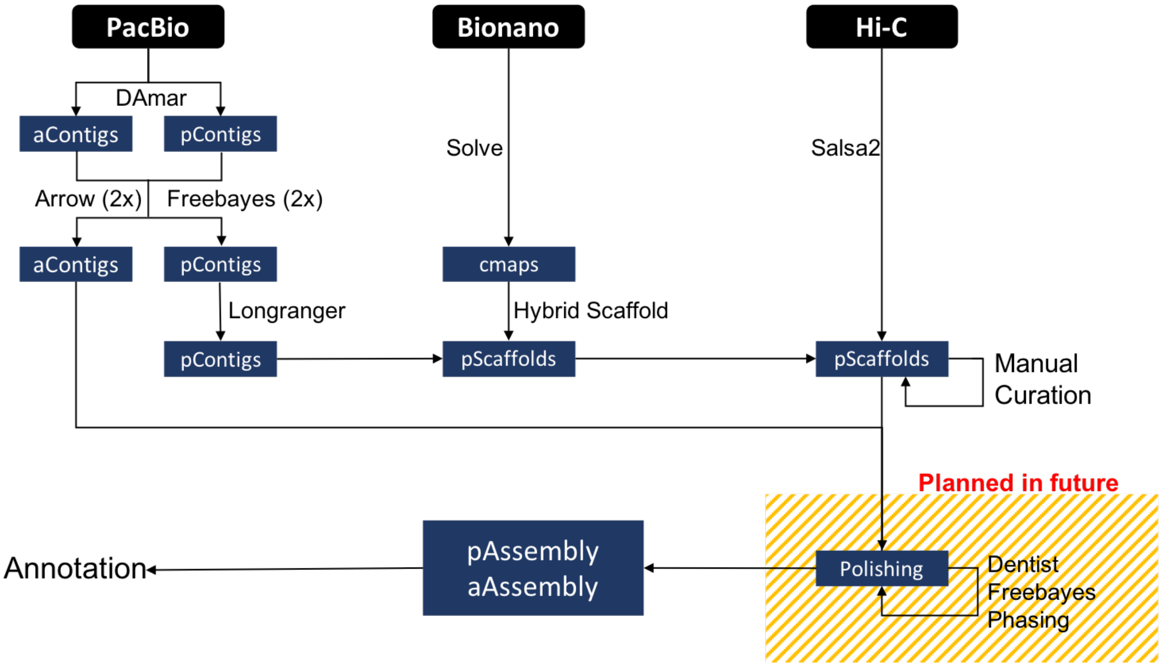

**Fig. S24: Hi-C maps for *M. myotis* prior (left figure) and post manual curation (right figure).** Hi-C maps were created by mapping and filtering the Hi-C read pairs by using the tools bwa, pairsamtools, pairix and cooler following the Hi-C data processing pipeline on https://hms-dbmi.github.io/hic-data-analysis-bootcamp. Left figure: Ellipse 1 shows that scaffold 2 contains a false join. It was split in the manual curation step. Ellipse 2 highlights two scaffolds, which were not joined in automated scaffolding steps but were manually integrated into scaffold 3.

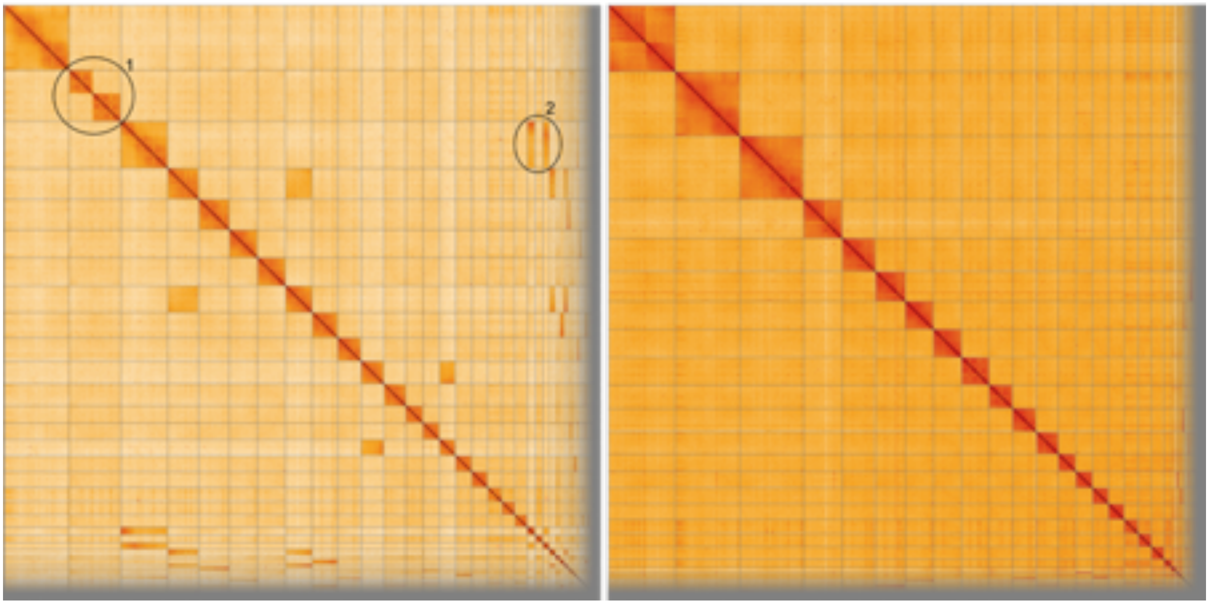

**Fig. S25: Comparison of scaffold lengths and chromosome lengths that were estimated from published karyotype images.** (A) *M. molossus*; (B) *M. myotis*; (C) *R. aegyptiacus*.

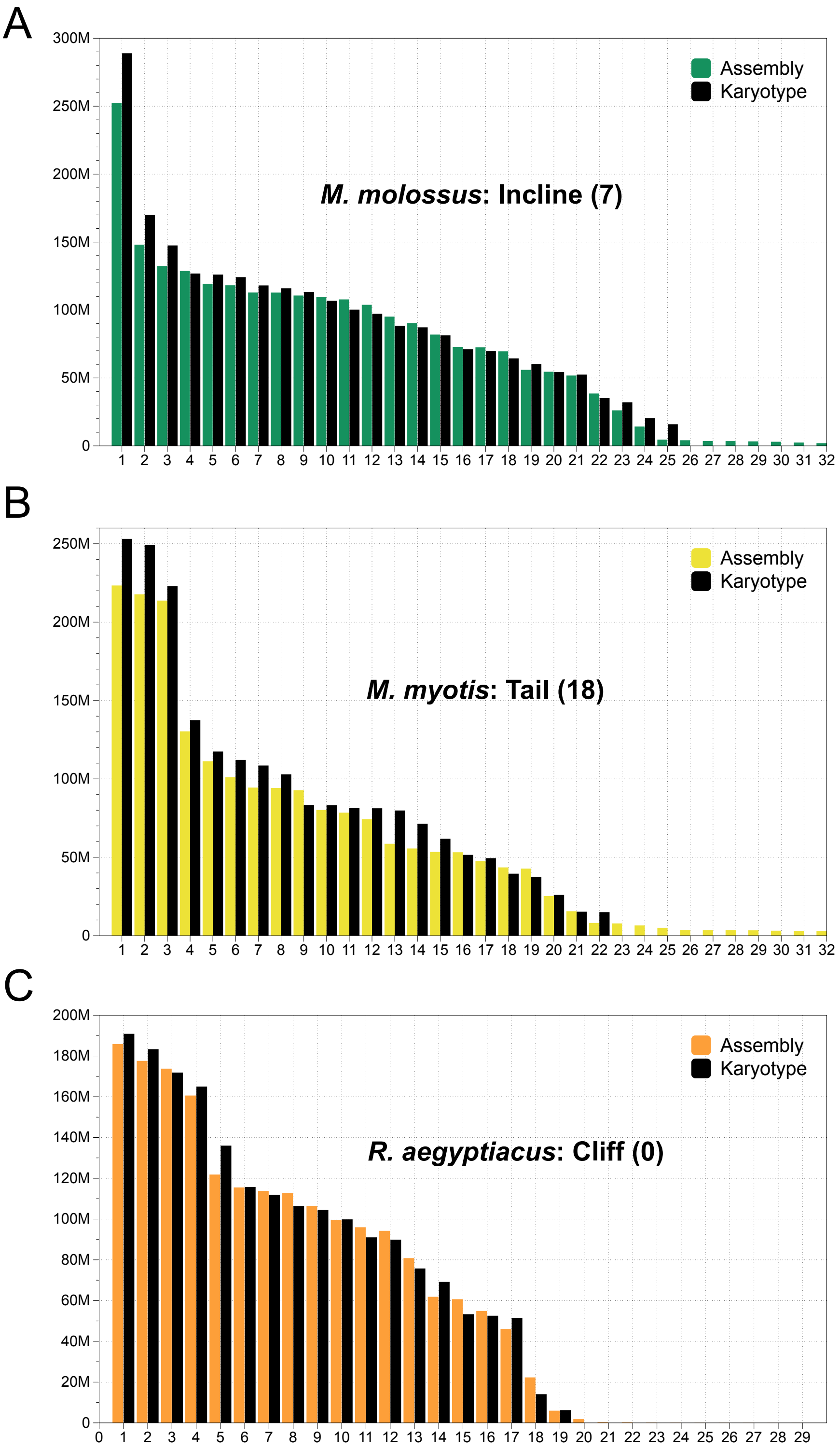

**Fig. S26: The tree topology inferred with a supermatrix of 12,931 genes.** Divergence times are calculated using the tree topology inferred with a supermatrix of 12,931 genes. Node calibrated with fossils are highlighted in red.

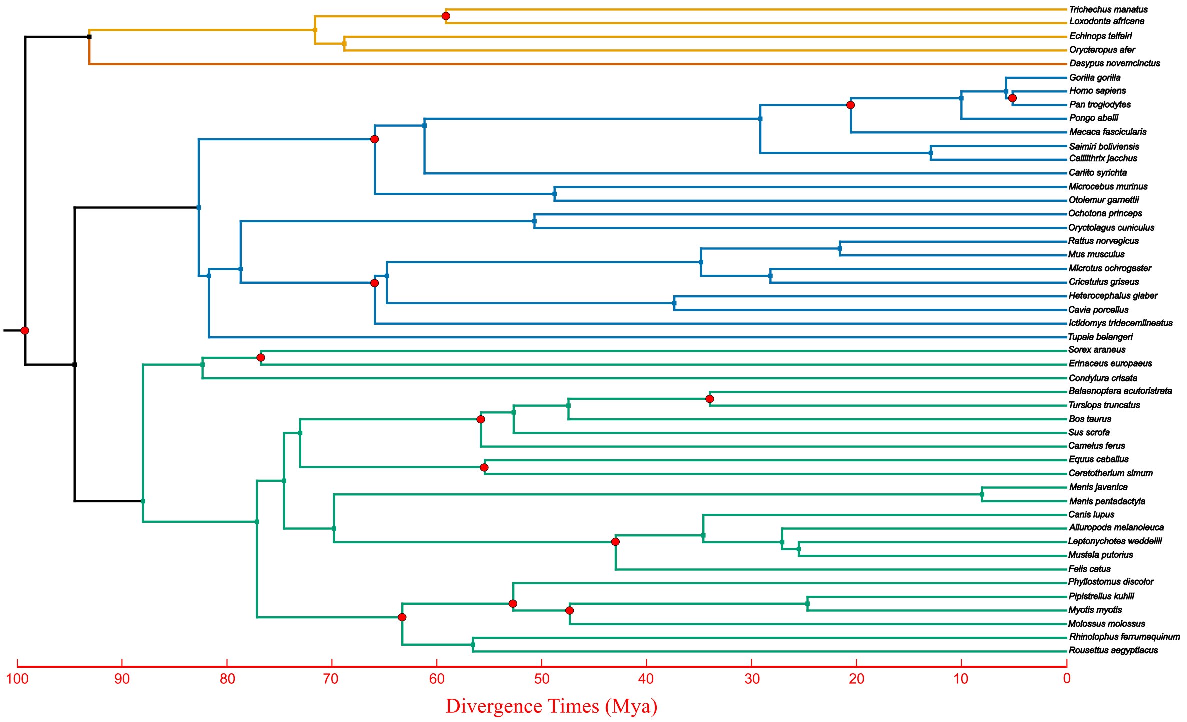

**Fig. S27: Loss of *PYHIN* genes in bats.** UCSC genome browser screenshot shows the *PYHIN* genes in the human genome (blue highlight). The alignment nets show that all six bats have large deletions that remove these genes, which is consistent with previous results that all bats have lost these genes[86]. Because these genes are also lost in several other mammals, we filtered them out in our genome-wide screen for gene losses.

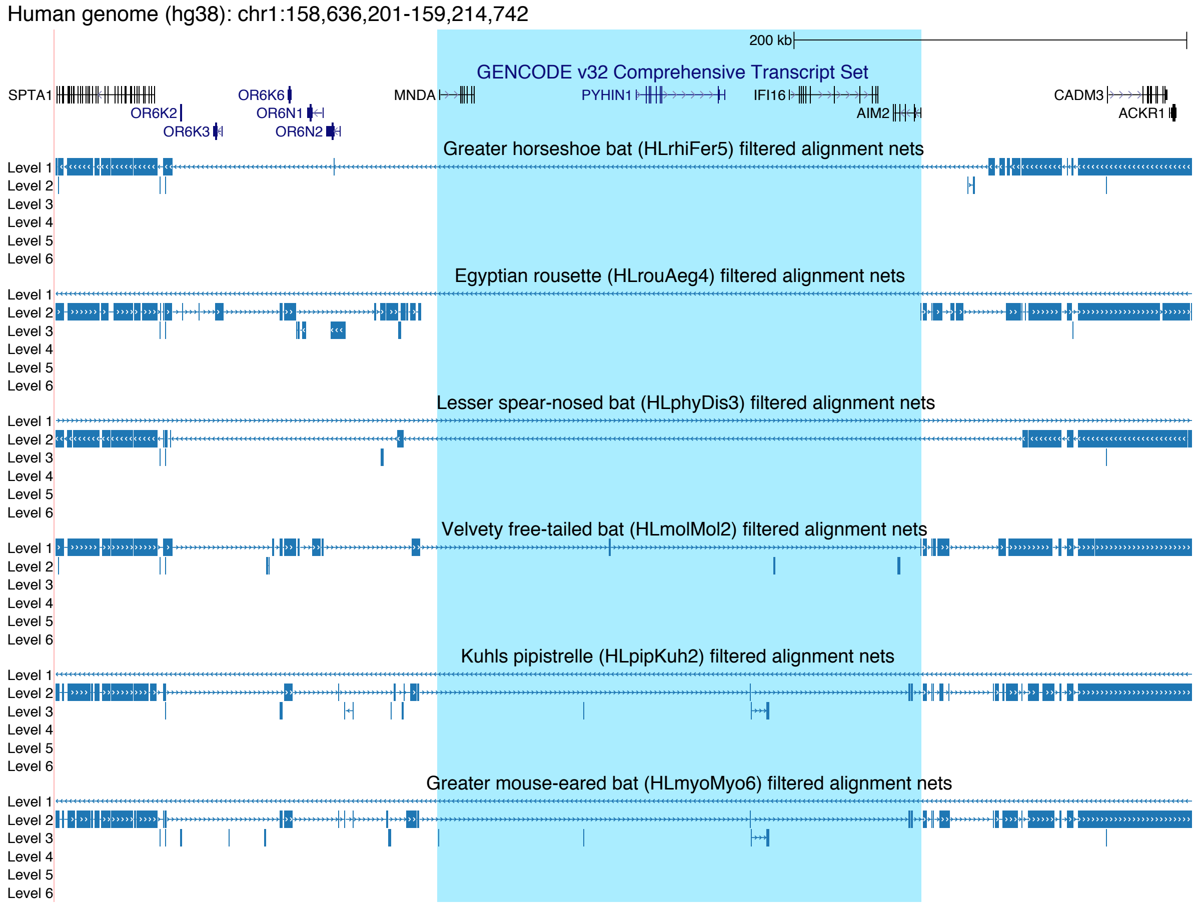

**Fig. S28: Gene tree showing the evolution of *APOBEC3* genes in mammals.** A phylogenetic tree was reconstructed using PhyML. We used the Z domain of the members of the panther gene family PTHR13857, added by *APOBEC3* genes we annotated in our six bat genomes. In addition to a possible small expansion in the ancestral bat lineage, the tree supports a scenario of several additional *APOBEC3* expansions in independent bat lineages.

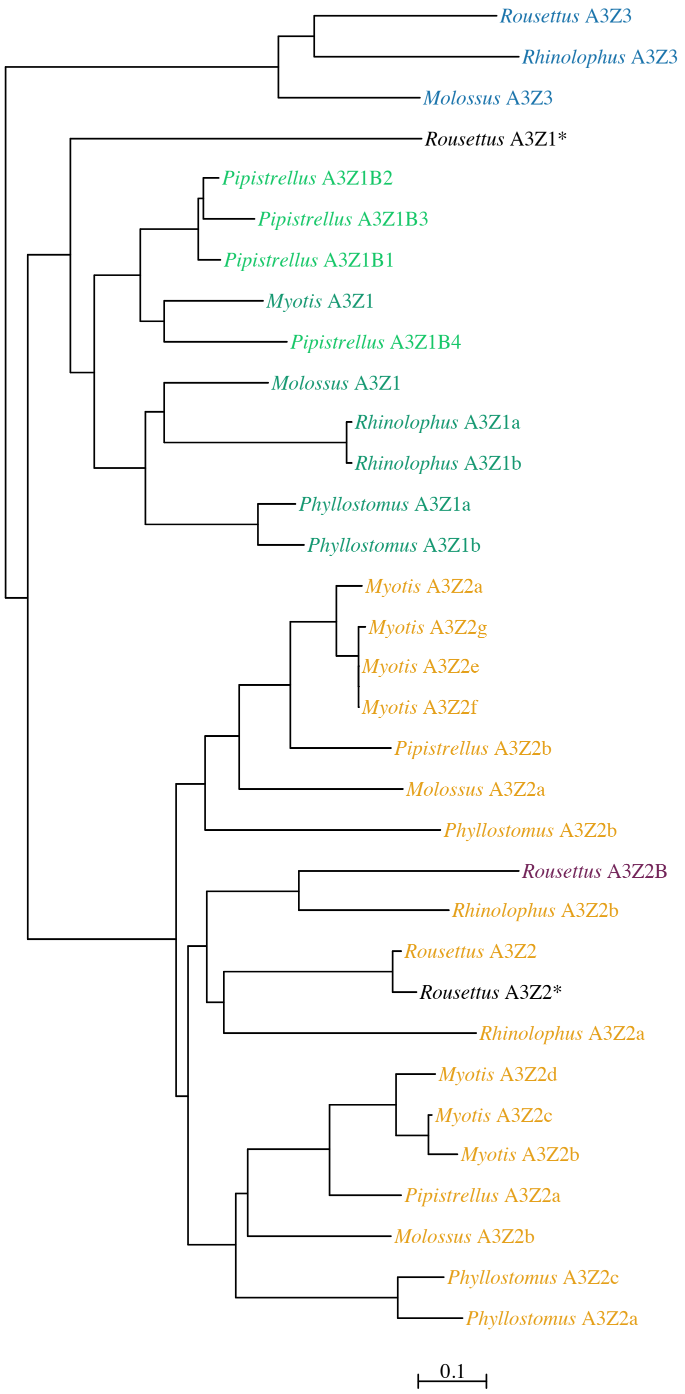

**Fig. S29: Genomic annotation of predicted miRNA loci in 6 bat species. A)** The number of known and novel miRNA located in exonic, intronic and intergenic regions; **B)** The percentage of known and novel miRNA located in exonic, intronic and intergenic regions.

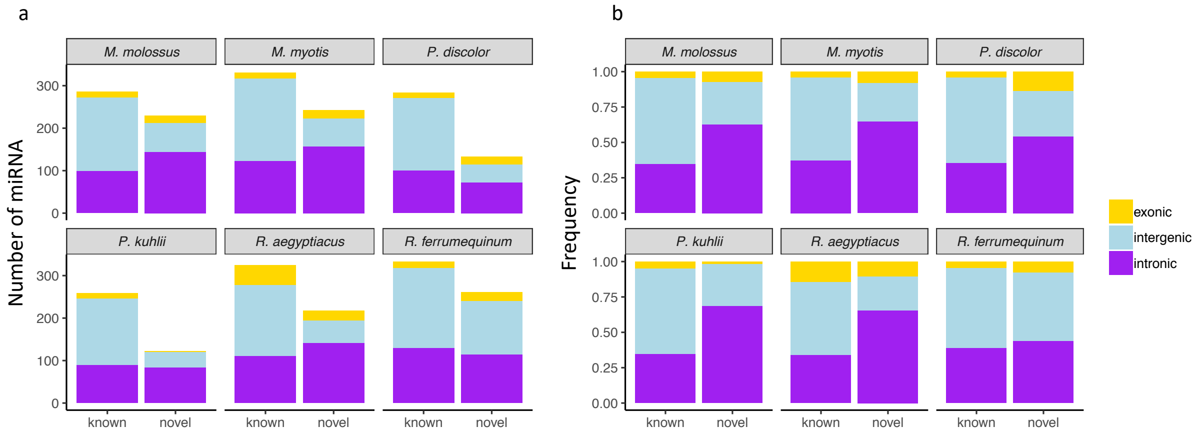

**Fig. S30: The empirical minimum free energy (NFE) distribution based on 1,000,000 human miRNA and target predictions by miranda.**

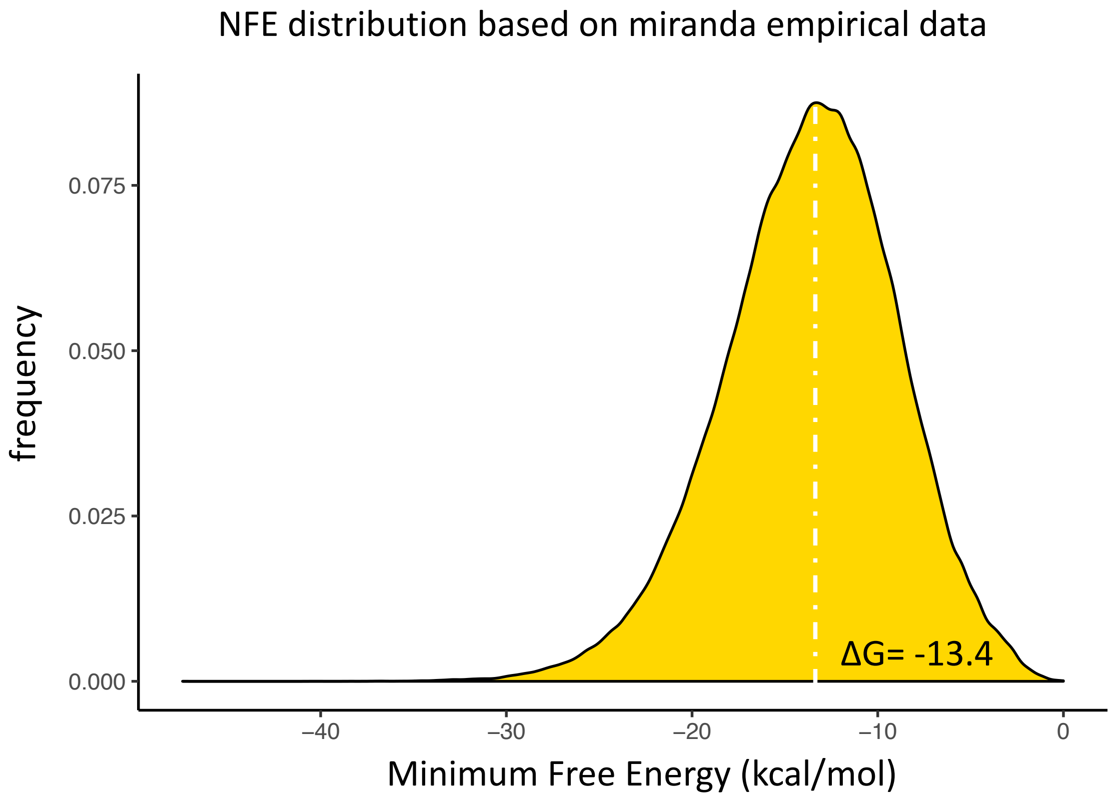

**Table S1: Information of 48 mammalian genomes used for comparative and phylogenetic analyses in this study.** The species highlighted in blue indicate 6 bat species whose genomes were sequenced in this study while the species in red indicate 7 species used as references for comparative analyses.

| Species name | Common name | Order | Genome version |
|---|---|---|---|
| *Trichechus manatus* | American manatee | Afrotheria | triMan1 |
| *Loxodonta africana* | African bush elephant | Afrotheria | loxAfr3 |
| *Orycteropus afer* | Aardvark | Afrotheria | oryAfe1 |
| *Echinops telfairi* | Lesser hedgehog | Afrotheria | echTel2 |
| *Dasypus novemcinctus* | Armardillo | Xenarthra | dasNov3 |
| *Gorilla gorilla* | Gorilla | Primates | gorGor5 |
| *Homo sapiens* | Human | Primates | hg38 |
| *Pan troglodytes* | Chimpanzee | Primates | panTro5 |
| *Pongo abelii* | Orangutan | Primates | ponAbe2 |
| *Macaca fascicularis* | Long-tailed macaque | Primates | macFas5 |
| *Saimiri boliviensis* | Squirrel monkey | Primates | saiBol1 |
| *Callithrix jacchus* | Common marmoset | Primates | calJac3 |
| *Carlito syrichta* | Tarsier | Primates | tarSyr2 |
| *Microcebus murinus* | Grey mouse lemur | Primates | micMur3 |
| *Otolemur garnettii* | Northern greater galago | Primates | otoGar3 |
| *Ochotona princeps* | American pika | Largomorpha | ochPri3 |
| *Oryctolagus cuniculus* | European rabbit | Largomorpha | oryCun2 |
| *Rattus norvegius* | Rat | Rodentia | Rnor6 |
| *Mus musculus* | Mouse | Rodentia | mm10 |
| *Microtus ochrogaster* | Vole | Rodentia | micOch1 |
| *Cricetulus griseus* | Chinese hamster | Rodentia | criGri1 |
| *Heterocephalus glaber* | Naked mole rat | Rodentia | hetGla2 |
| *Cavia porcellus* | Guinea pig | Rodentia | cavPor3 |
| *Ictidomys tridecemlineatus* | Thirteen-lined ground squirrel | Rodentia | speTri2 |
| *Tupaia belangeri* | Northern tree shrew | Scandentia | tupBel1 |
| *Sorex araneus* | Common shrew | Eulipothphla | sorAra2 |
| *Erinaceus europaeus* | European hedgehog | Eulipothphla | eriEur2 |
| *Condylura cristata* | Star-nosed mole | Eulipothphla | conCri1 |
| *Balaenoptera acutorostrata* | Minke whale | Cetartiodactyla | balAcu1 |
| *Tursiops truncatus* | Bottlenose dolphin | Cetartiodactyla | turTru3 |
| *Bos taurus* | Cattle | Cetartiodactyla | bosTau8 |
| *Sus scrofa* | Pig | Cetartiodactyla | susScr11 |
| *Camelus ferus* | Camel | Cetartiodactyla | camFer2 |
| *Equus caballus* | Horse | Perissodactyla | equCab3 |
| *Ceratotherium simum* | White rhinoceros | Perissodactyla | cerSim1 |
| *Manis javanica* | Sunda pangolin | Pholidota | manJav1 |
| *Manis pentadactyla* | Chinese pangolin | Pholidota | manPen1 |
| *Canis lupus familiaris* | Dog | Carnivora | canFam3 |
| *Ailuropoda melanoleuca* | Giant panda | Carnivora | ailMel1 |
| *Leptonychotes weddellii* | Seal | Carnivora | lepWed1 |
| *Mustela putorius* | Polecat | Carnivora | musPut1 |
| *Felis catus* | Cat | Carnivora | felCat8 |
| *Phyllostomus discolour* | Pale spear-nosed bat | Chiroptera | phyDis3 |
| *Pipistrellus kuhlii* | Kuhl's pipistrelle | Chiroptera | pipKuh2 |
| *Myotis myotis* | Greater mouse-eared bat | Chiroptera | myoMyo6 |
| *Molossus molossus* | Velvety free-tailed bat | Chiroptera | molMol2 |
| *Rhinolophus ferrumequinum* | Greater horseshoe bat | Chiroptera | rhiFer5 |
| *Rousettus aegyptiacus* | Egyptian fruit bat | Chiroptera | rouAeg4 |

**Table S2: The final genomes (all lengths in Mbp).** N50 values correspond to *post hoc* genome size of the final assemblies (sum of the length of all scaffolds).

| Species | Scaffolds | | | | Primary Contigs | | | | |
|---|---|---|---|---|---|---|---|---|---|
| | Number | Total Length | Maximum Length | N50 | Number | Total Length | Maximum Length | N50 | Avg. QV |
| *M. myotis* | 92 | 2,003 | 223 | 94.45 | 630 | 1,974 | 52 | 12.51 | 42.7 |
| *P. kuhlii* | 202 | 1,776 | 197 | 80.24 | 597 | 1,763 | 51 | 10.59 | 40.8 |
| *R. ferrumequinum* | 50 | 2,075 | 126 | 92.00 | 347 | 2,056 | 81 | 21.75 | 46.2 |
| *P. discolor* | 41 | 2,095 | 215 | 171.08 | 451 | 2,059 | 72 | 15.51 | 42.9 |
| *R. aegyptiacus* | 29 | 1,894 | 186 | 113.81 | 271 | 1,867 | 81 | 22.00 | 43.9 |
| *M. molossus* | 60 | 2,319 | 252 | 110.67 | 412 | 2,268 | 77 | 22.17 | 42.2 |

**Table S3: Manual curation and consistency with karyotypes.**

| Species | Splits / Joins | # of Chromosomes in Karyotype = N | Correlation with top N Scaffolds | Tail Status | % of Data in N Largest Scaffolds |
|---|---|---|---|---|---|
| ***M. myotis*** | 2 / 6 | 22 | 0.99 | Tail (18) | 95.6% |
| ***P. kuhlii*** | not curated | 22 | 0.96 | Tail (33) | 86.6% |
| ***R. ferrumequinum*** | 3 / 1 | 29 | 0.98 | Incline (5) | 98.9% |
| ***P. discolor*** | 10 / 13 | 16 | 0.98 | Cliff (-1) | 99.6% |
| ***R. aegyptiacus*** | 0 / 5 | 18 | 0.99 | Cliff (0) | 99.5% |
| ***M. molossus*** | 2 / 6 | 24 | 0.98 | Incline (7) | 98.3% |

**Table S4: The presence of highly-conserved BUSCO genes in the genome and in the gene annotations.**

| | species | BUSCO applied to genome assembly | | | BUSCO applied to gene annotation | | |
|---|---|---|---|---|---|---|---|
| | | Complete | Fragmented | Missing | Complete | Fragmented | Missing |
| | *Homo* | 94.59% | 2.46% | 2.95% | 99.95% | 0.00% | 0.05% |
| | *Mus* | 95.30% | 2.36% | 2.34% | 99.83% | 0.02% | 0.15% |
| | *Canis* | 95.30% | 2.36% | 2.34% | 98.56% | 1.00% | 0.44% |
| | *Felis* | 94.76% | 2.61% | 2.63% | 98.39% | 0.97% | 0.63% |
| | *Equus* | 96.22% | 2.07% | 1.71% | 97.78% | 0.85% | 1.36% |
| | *Bos* | 93.71% | 3.05% | 3.24% | 98.98% | 0.68% | 0.34% |
| | *Sus* | 94.08% | 3.51% | 2.41% | 98.85% | 0.68% | 0.46% |
| Bat1K assemblies | *Rhinolophus* | 95.42% | 2.36% | 2.22% | 99.66% | 0.17% | 0.17% |
| | *Rousettus* | 95.83% | 1.92% | 2.24% | 99.34% | 0.29% | 0.37% |
| | *Phyllostomus* | 94.91% | 1.80% | 3.29% | 99.66% | 0.15% | 0.19% |
| | *Molossus* | 92.93% | 3.17% | 3.90% | 99.49% | 0.19% | 0.32% |
| | *Pipistrellus* | 95.35% | 2.36% | 2.29% | 99.56% | 0.17% | 0.27% |
| | *Myotis* | 94.44% | 2.88% | 2.68% | 99.63% | 0.24% | 0.12% |

**Table S5: Analysis of ultraconserved elements (UCEs) that do not align with ≥ 85% identity and at least 150 bp.**

| UCE not found | species in which the UCE was not found | assembly | Reason | Reason |
|---|---|---|---|---|
| uc.157 | *Miniopterus* | minNat1 | Assembly artifact | UCE overlaps a 1189 bp assembly gap |
| uc.158 | *Miniopterus* | minNat1 | Assembly artifact | UCE overlaps a 1833 bp assembly gap |
| uc.159 | *Miniopterus* | minNat1 | Assembly artifact | UCE overlaps a 1786 bp assembly gap |
| uc.157 | Cow | bosTau8 | Assembly artifact | assembly) |
| uc.170 | Cow | bosTau8 | Assembly artifact | mutus) assemblies |
| uc.10 | Dog | canFam3 | Assembly artifact | UCE overlaps a 300 bp assembly gap contained in a 4.3 kb locus in canFam3. However gap size is likely underestimated as the corresponding locus in human is 100 kb and several chrUn parts align to the human locus (but not the UCE). UCE is fully present in the Dingo (Canis lupus dingo) assembly |
| uc.157 | Dog | canFam3 | Assembly artifact | UCE overlaps a 476 bp region in canFam2 (chr3:31,944,063-31,944,538) with low quality scores (Phred <30), this low-quality region has remained identical in canFam3. Searching the NCBI traces finds a single Sanger Read (ti:294708094) that aligns well but was apparently not incorporated in the assembly. UCE is fully present in the Dingo (Canis lupus dingo) assembly |
| uc.296 | Dog | canFam3 | Assembly artifact | UCE overlaps a 340 bp assembly gap. UCE is fully present in the Dingo (Canis lupus dingo) assembly |
| uc.3 | Dog | canFam3 | Assembly artifact | UCE overlaps a 4474 bp assembly gap. UCE is fully present in the Dingo (Canis lupus dingo) assembly |
| uc.157 | Cat | felCat8 | Assembly artifact | UCE overlaps a 643 bp assembly gap. UCE is fully present in the Leopard (Panthera pardus) assembly |
| uc.398 | Cat | felCat8 | Assembly artifact | assembly |
| uc.47 | *Myotis* | this study | real divergence | real divergence |
| uc.394 | *Pipistrellus* | this study | real divergence | real divergence |
| uc.446 | *Pipistrellus* | this study | real divergence | real divergence |
| uc.47 | *Pipistrellus* | this study | real divergence | real divergence |

**Table S6: Related species used for Genome Threader alignments.** Details of publicly available cDNA and protein data aligned to 6 genome assemblies for annotation from a related species.

| Reference species | Query species | Query source | No. of query cDNA | No. of query peptides | Related at taxonomic rank |
|---|---|---|---|---|---|
| *Molossus molossus* | *Miniopterus natalensis* | Refseq | 25,266 | 25,266 | Superfamily: Vespertilionoidea |
| *Myotis myotis* | *Myotis lucifugus* | Ensembl | 22,432 | 20,719 | Genus: Myotis |
| *Phyllostomus discolor* | *Desmodus rotundus* | Refseq | 28,829 | 28,829 | Family: Phyllostomidae |
| *Pipistrellus kuhlli* | *Eptesicus fuscus* | Refseq | 18,724 | 18,263 | Subfamily: Vespertilioninae |
| *Rhinolophus ferrumequinum* | *Rhinolophus sinicus* | Refseq | 29,785 | 29,785 | Genus: Rhinolophus |
| *Rousettus aegyptiacus* | *Pteropus vampyrus* | Ensembl | 18,086 | 17,053 | Subamily: Pteropodinae |

**Table S7: Sources of RNA-Seq and Iso-seq transcriptomic data that we used for annotating genes.**

| Species | Tissue | Accession Number/ Bioproject |
|---|---|---|
| *Myotis myotis* | Brain | This study |
| | Heart | This study |
| | Liver | This study |
| | Kidney | This study |
| *Molossus molossus* | Blood | This study |
| *Phyllostomus discolor* | Brain | PRJNA291690 |
| *Pipistrellus kuhlii* | Fibroblast | PRJNA565655 |
| *Rhinolophus ferrumequinum* | Brain | SRR1048140 |
| | Brain | SRR1048142 |
| | Liver | SRR2754983 |
| | Liver | SRR2757329 |
| | Intestine | SRR6749599 |
| | Intestine | SRR6749600 |
| | Intestine | SRR6749601 |
| | Intestine | SRR6749602 |
| *Rousettus aegyptiacus* | Testes | SRR2914372 |
| | Liver | SRR2914369 |
| | Kidney | SRR2914360 |
| | Heart | SRR2914359 |
| | Brain | SRR2914295 |
| | Liver | SRR2914059 |
| | Kidney | SRR2913355 |
| | Heart | SRR2913354 |
| | Brain | SRR2913353 |

**Table S8: Overview of samples used for Iso-seq.**

| Species | Year of collection | Location | Tissue | Storage | Extraction | RIN |
|---|---|---|---|---|---|---|
| *Molossus molossus* | 2017 | Panama | brain | snap frozen | RNAeasy | 9.1 |
| | | | testes | snap frozen | RNAeasy | 9.0 |
| *Myotis myotis* | 2015 | Limerzel, France | brain | snap frozen | RNAeasy | 8.4 |
| | | | liver | snap frozen | RNAeasy | 8.9 |
| | | | kidney | snap frozen | RNAeasy | 9.1 |
| *Pipistrellus kuhlii* | 2015 | Italy | brain | snap frozen | Chloroform-Isopropanol | 8.1 |
| *Phyllostomus discolor* | 2016 | Munich, Germany, (LMU captive colony) | brain | snap frozen | Relia prep | 7.4 |
| | | | testes | snap frozen | RNAeasy | 9.7 |
| *Rhinolophus ferrumequinum* | 2018 | United Kingdom | brain | snap frozen | RNAeasy | 9.1 |
| *Rousettus aegyptiacus* | 2018 | Berkeley, USA, (captive colony) | brain | Qiazol frozen | RNAeasy | 8.6 |
| | | | testes | Qiazol frozen | RNAeasy | 8.7 |

**Table S9: Best-fit models of sequence evolution for all coding genes, CNEs and 1st +2nd codon site alignments were determined using IQTREE.**
**(See separate file)**

**Table S10: The name, alignment length and number of taxa present for 10,857 conserved non-coding elements (CNEs).**
**(See separate file)**

**Table S11: The 15 topologies showing different arrangements of Laurasiatheria and the number of gene trees supporting them with the highest likelihood.** Positions of other mammals are fixed relative to the coding-gene supermatrix topology (topology 1).
**(See separate file)**

**Table S12: The metrics of the alignments of 12,931 genes.**
**(See separate file)**

**Table S13: Results of the genome-wide screen for positive selection in coding genes.**
**(See separate file)**

**Table S14: All 2,453 genes relating to ageing/immunity/metabolism used in selection analysis with PAML and their taxonomic representation.**
**(See separate file)**

**Table S15: The significant genes under positive selection along the bat ancestral branch using PAML.**
**(See separate file)**

**Table S16: Genes that were inferred to be lost in all 6 bats analysed in this study.**

| Gene Symbol | Ensembl Gene ID |
|---|---|
| *AS3MT* | ENSG00000214435 |
| *HIST1H4K* | ENSG00000273542 |
| *IL36G* | ENSG00000136688 |
| *KLK4* | ENSG00000167749 |
| *KRBA2* | ENSG00000184619 |
| *LRRC70* | ENSG00000186105 |
| *MS4A3* | ENSG00000149516 |
| *U2AF1L4* | ENSG00000161265 |
| *ZBED9* | ENSG00000232040 |
| *ZFP30* | ENSG00000120784 |

**Table S17: Gene families that were estimated to have undergone a contraction or expansion in the ancestral bat lineage.** Values shown are P-values for evidence in shift in the rate of birth/death of a gene family along the given branch. "Corrected Chiroptera" gives the P-value after FDR correction along the Chiroptera ancestral branch. The Family ID is the PANTHER family ID. Where multiple PANTHER families were collapsed, IDs were concatenated. If no human protein was present in a family, no PANTHER ID was not assigned, and an internal identifier used. Expansion/Contraction was determined by comparing the Chiroptera ancestor to the inferred scrotiferan ancestor.
**(See separate file)**

**Table S18: The Number of viral integrations in analysed genomes.** The number of all retroviral integrations found in six bat genomes for each viral class and protein (Pol, Gag, Env). Pol sequences were additionally searched in seven reference genomes and their numbers were compared.
**(See separate file)**

**Table S19: The number of 286 conserved miRNA gene copies across 48 mammalian taxa based on the *de novo* genomic prediction using the Infernal pipeline.**
**(See separate file)**

**Table S20: Oligonucleotide sequences used for cloning**. Restriction site tags and spacers between individual miRNA binding sites are highlighted in bold, mature miRNAs are italic, and miRNA seed sequences are underlined.

| Insert for cloning | SENSE OLIGO (5'→3') | ANTISENSE OLIGO (5'→3') |
|---|---|---|
| Bat-miR-19125 | **CTAGA**TCCCTTGGAAAGAGCCTTGTTTTGGAAGGGAAGGGGGAAGAGGCTCTGCCCTTGACC*TCTACCCTTTGTTTCCTTCCAGC*CTTTGTCCAGGAGTTGAGGAAGAGG**G** | **TCGAC**CCTCTTCCTCAACTCCTGGACAAAGGCTGGAAGGAAACAAAGGGTAGAGGTCAAGGGCAGAGCCTCTTCCCCCTTCCCTTCCAAAACAAGGCTCTTTCCAAGGGA**T** |
| Bat-miR-4665 | **CTAGA**CCCCTACTTGCAGTTGGTCCGACGG*TTGTGGGTTATTGTTAAG*CTGATTAACATTGTCTCCCTCCACACAACCACATTTGACTGACTTTGTATTTTGCCCTAGTC**G** | **TCGAC**GACTAGGGCAAAATACAAAGTCAGTCAAATGTGGTTGTGTGGAGGGAGACAATGTTAATCAGCTTAACAATAACCCACAACCGTCGGACCAACTGCAAGTAGGGG**T** |
| Bat-miR-6665 | **CTAGA**ACAAAGTAGGTTAGATCTTGCCAGA*TTAGGTGGAGATTCTCGCAGGG*GGAGTTCAACTTCATATACCCTTGCAAGATACTCCTCTGTCTGGAAAGGTCTTCCTCT**G** | **TCGAC**AGAGGAAGACCTTTCCAGACAGAGGAGTATCTTGCAAGGGTATATGAAGTTGAACTCCCCCTGCGAGAATCTCCACCTAATCTGGCAAGATCTAACCTACTTTGT**T** |
| Bat-miR-19125_sensor | **TCGAG**GCTGGAAGGAAACAAAGGGTAGAGAATATGCTGGAAGGAAACAAAGGGTAGA**T** | **CTAGA**TCTACCCTTTGTTTCCTTCCAGCATATTCTCTACCCTTTGTTTCCTTCCAGC**C** |
| Bat-miR-4665_sensor | **TCGAG**CTTAACAATAACCCACAAGAATATCTTAACAATAACCCACAA**T** | **CTAGA**TTGTGGGTTATTGTTAAGATATTCTTGTGGGTTATTGTTAAG**C** |
| Bat-miR-6665_sensor | **TCGAG**CCCTGCGAGAATCTCCACCTAAGAATATCCCTGCGAGAATCTCCACCTAA**T** | **CTAGA**TTAGGTGGAGATTCTCGCAGGGATATTCTTAGGTGGAGATTCTCGCAGGG**C** |
| Bat-miR-337 | **CTAGA**ACAGTCAGTAAGTGGGGGGTGAGAACGGCTTCATCCAGGAGTTGATGCCCAGTTATCCAGC*GCCTAGA*TGATGCCTTTCTTCATCCCCTTCAA**G** | **TCGAC**TTGAAGGGGATGAAGAAAGGCATCATCTAGGCGCTGGATAACTGGGCATCAACTCCTGGATGAAGCCGTTCTCACCCCCCACTTACTGACTGT**T** |
| hsa-miR-337 | **CTAGA**GTAGTCAGTAGTTGGGGGGTGGGAACGGCTTCATACAGGAGTTGATGCACAGTTATCCAGC*TCCTATA*TGATGCCTTTCTTCATCCCCTTCAA**G** | **TCGAC**TTGAAGGGGATGAAGAAAGGCATCATATAGGAGCTGGATAACTGTGCATCAACTCCTGTATGAAGCCGTTCCCACCCCCCAACTACTGACTAC**T** |
| Bat-miR-337_sensor | **TCGAG**GAAGAAAGGCATCATCTAGGCGGAATATGAAGAAAGGCATCATCTAGGCG**T** | **CTAGA**CGCCTAGATGATGCCTTTCTTCATATTCCGCCTAGATGATGCCTTTCTTC**C** |
| hsa-miR-337_sensor | **TCGAG**GAAGAAAGGCATCATATAGGAGGAATATGAAGAAAGGCATCATATAGGAG**T** | **CTAGA**CTCCTATATGATGCCTTTCTTCATATTCCTCCTATATGATGCCTTTCTTC**C** |

**Table S21: The statistics of 3'UTR analysis for 6 bat genomes.**

| | Total 3UTR | Different genes with overlapped 3UTR loci | After merging overlapped coordinates | Pseudo 3UTR |
|---|---|---|---|---|
| *M. molossus* | 13,671 | 290 | 11,912 | 8,613 |
| *M. myotis* | 13,263 | 406 | 11,024 | 8,811 |
| *P. kuhlii* | 6,891 | 182 | 6,372 | 5,612 |
| *P. discolor* | 15,122 | 476 | 12,196 | 9,030 |
| *R. ferrumequinum* | 7,913 | 226 | 7,194 | 6,346 |
| *R. aegyptiacus* | 16,115 | 327 | 13,394 | 9,519 |

**Table S22: The gene targets of human and bat miR-337 predicted by RNAhybrid and miranda.** The gene targets, which were specific to bat and human and were shared between bat and human, were listed respectively.

**(See separate file)**

**Table S23: Summary of miRNA sequencing and analysis in 6 bat genomes.**

| Species | Tissue | Raw reads | Mapping rate | Known miRNA | Novel miRNA |
|---|---|---|---|---|---|
| *Myotis myotis* | Brain | 44,256,216 | 90.3% | 329 | 242 |
| | Kidney | 41,823,612 | 82.7% | | |
| | Liver | 42,013,686 | 81.8% | | |
| *Pipistrellus kuhlii* | Brain | 46,857,126 | 92.6% | 258 | 122 |
| | Kidney | 35,303,058 | 73.2% | | |
| | Liver | 39,045,247 | 75.0% | | |
| *Molossus molossus* | Brain | 35,168,361 | 93.8% | 286 | 229 |
| | Kidney | 44,074,092 | 91.6% | | |
| | Liver | 36,055,759 | 86.5% | | |
| *Phyllostomus discolor* | Brain | 107,798,649 | 94.7% | 284 | 133 |
| | Kidney | 39,805,199 | 91.7% | | |
| | Liver | 27,597,872 | 89.9% | | |
| *Rhinolophus ferrumequinum* | Brain | 52,426,146 | 91.8% | 332 | 261 |
| | Kidney | 50,908,913 | 91.1% | | |
| | Liver | 48,450,654 | 86.2% | | |
| *Rousettus aegyptiacus* | Brain | 25,458,780 | 91.6% | 325 | 217 |
| | Kidney | 45,673,617 | 91.6% | | |
| | Liver | 33,831,764 | 91.0% | | |

**Table S24: The summary of 12 novel miRNA at the ancestral bat lineage.** These newly-evolved miRNA were not found in any other species.
**(See separate file)**

**Table S25: Overview species and tissues for high molecular weight genomic DNA (HMW gDNA) extraction.**

| Species | Sex | Provided by | Year of collection | Location |
|---|---|---|---|---|
| *Molossus molossus* | male | Dina Deichman | 2018 | Gamboa, Panama (9.1165° N, 79.6965° W) |
| *Myotis myotis* | female | Emma Teeling & Sébastien Puechmaille | 2015 | Limerzel, France (47.6333° N, 2.3500° W) |
| *Pipistrellus kuhlii* | male | Emma Teeling & Andrea Locatelli | 2017 | Bergamo, Italy (45.7430° N, 9.5831° E) |
| *Phyllostomus discolor* | male | Sonja Vernes | 2016 | Munich, Germany (Captive colony) |
| *Rhinolophus ferrumequinum* | female | Gareth Jones | 2016 | United Kingdom (51.7108° N, 2.2776° W) |
| *Rousettus aegyptiacus* | male | Sonja Vernes | 2017 | Berkeley, USA (Captive colony) |

**Table S26: The information of gDNA extraction for 6 bat species.** The table includes gDNA extraction protocol, size range of extracted gDNA determined by PFGE, and applied technologies (CLR: continuous long reads, PCE: phenol-chloroform-extraction). MA= MagAttract used for Pacbio CLR and 10X (40-60kb); Plug= 50-500kb only used for Bionano.

| Species | Tissue | Extraction | Size range (kb) | PacBio CRL | Bionano | 10x linked read |
|---|---|---|---|---|---|---|
| ***Molossus molossus*** | muscle | PCE | 50 - 150 | X | | |
| | liver | plug | 50 - > 500 | | X | X |
| ***Myotis myotis*** | muscle | PCE | 50 - 300 | X | | X |
| | muscle | Plug | 50 - > 500 | | X | |
| ***Pipistrellus kuhlii*** | muscle | PCE | 50 - 250 | X | | X |
| | heart | Plug | 50 - 400 | | X | |
| ***Phyllostomus discolor*** | muscle | Plug<br>MA | 40-60 | | X | |
| ***Rhinolophus ferrumequinum*** | lung | PCE | 50 - 250 | X | | X |
| | lung | Plug | VGL | | X | |
| ***Rousettus aegyptiacus*** | muscle | PCE | 50 - 250 | X | | |
| | liver | plug | 50 - > 500 | | X | X |

**Table S27: The information of Pacbio CLR library preparation and sequencing.**

| Species | Shearing size (kb) | Size selection (kb) | PacBio polymerase | No. of SMRT cells | Average yield per SMRT cell (Gb) | Average insert N50 per SMRT cell (kb) |
|---|---|---|---|---|---|---|
| *Molossus molossus* | 75 | 25 | 2.1 | 26 | 4.8 | 23.7 |
| *Myotis myotis* | 35 | 12 - 15 | 2.0 | 39 | 3.31 | 14.3 |
| *Pipistrellus kuhlii* | 35 - 40 | 15 - 20 | 2.0 | 48 | 2.55 | 13.76 |
| *Phyllostomus discolor* | 35 - 40 | 12 - 15 | 2.0 | 43 | 4.10 | 15.05 |
| *Rhinolophus ferrumequinum* | 40 | 18 | 2.0 | 25 | 5.09 | 18.25 |
| *Rousettus aegyptiacus* | 60 | 18 | 2.0 | 35 | 3.92 | 18.08 |

**Table S28: Sequencing depth, effective genome coverage and mean molecular length calculated by the 10x Supernova tool.**

| Species | Long gDNA | megasize gDNA | Sequenced fragments (Mi reads) | Genome coverage (linked reads) | Mean molecular length (kb) |
|---|---|---|---|---|---|
| *Molossus molossus* | - | X | 354 | 49x | 131.9 |
| *Myotis myotis* | X | - | 319 | 44x | 29.4 |
| *Pipistrellus kuhlii* | X | - | 327 | 46x | 19.7 |
| *Phyllostomus discolor* | X | - | 789 | 109x | 16.0 |
| *Rhinolophus ferrumequinum* | X | - | 345 | 48x | 28.8 |
| *Rousettus aegyptiacus* | - | X | 365 | 51 | 97.9 |

**Table S29: Statistics of PacBio dataset.** The Raw data set contains all PacBio subreads longer then 500 b. The Filtered_1 data set contains only statistics for the longest read of each Zero-mode waveguide (ZMW). The Filtered_2 data set, that was used for the assembly, contains only the longest subread per ZMW with a minimum length of 4 kb.

| | | Number of SMRT Cells | Number of Reads (M) | Total Base Pairs (Gbp) | Estimated Coverage | Average Read Length (Kbp) | Longest Read (Kbp) | Finish Date |
|---|---|---|---|---|---|---|---|---|
| **M. myotis** | *Raw* | 48 | 21.4 | 182.1 | 90.9 | 8.5 | 150.6 | May 2017 |
| | *Filtered_1* | | 16.4 | 150.4 | 75.1 | 9.2 | | |
| | *Filtered_2* | | 11.4 | 140.2 | 70.0 | 13.8 | | |
| **P. kuhlii** | *Raw* | 48 | 16.0 | 143.6 | 80.8 | 9.0 | 150.4 | Jun 2017 |
| | *Filtered_1* | | 12.8 | 122.4 | 68.9 | 9.6 | | |
| | *Filtered_2* | | 9.2 | 115.0 | 64.7 | 12.6 | | |
| **R. ferrumequinum** | *Raw* | 25 | 14.5 | 152.0 | 73.3 | 10.5 | 161.2 | Aug 2017 |
| | *Filtered_1* | | 11.3 | 127.1 | 61.2 | 11.2 | | |
| | *Filtered_2* | | 8.2 | 121.0 | 58.3 | 14.8 | | |
| **P. discolor** | *Raw* | 43 | 18.2 | 163.5 | 78.0 | 9.0 | 148.0 | Mar 2018 |
| | *Filtered_1* | | 15.6 | 148.7 | 71.0 | 9.5 | | |
| | *Filtered_2* | | 10.8 | 138.9 | 66.3 | 12.8 | | |
| **R. aegyptiacus** | *Raw* | 33 | 12.0 | 122.3 | 64.6 | 10.2 | 174.1 | Feb 2018 |
| | *Filtered_1* | | 11.1 | 117.5 | 62.0 | 10.6 | | |
| | *Filtered_2* | | 7.7 | 110.5 | 58.3 | 14.4 | | |
| **M. molossus** | *Raw* | 26 | 11.1 | 135.1 | 58.3 | 12.1 | 101.0 | Sep 2018 |
| | *Filtered_1* | | 9.0 | 124.9 | 53.8 | 13.8 | | |
| | *Filtered_2* | | 6.7 | 120.2 | 51.8 | 18.0 | | |

**Table S30: Statistics of 10x datasets and yields over 100 Kbp.** Raw: raw Illumina sequencing statistics; barcode removed: Illumina read pair statistics after trimming off the 16 bp 10X and 7 bp Illumina barcodes from the R1 reads; Molecule length >100K: Statistics for Molecule lengths that were calculated from the final assemblies and the tool bxcheck (https://github.com/pd3/bxcheck)

| | | Number of Lanes | Number of Reads (M) | Total Base Pairs (Gbp) | Estimated Coverage | Read Cloud N50 (Kbp) | Finish Date |
|---|---|---|---|---|---|---|---|
| ***M. myotis*** | *Raw* | 4 | 319.4 | 95.8 | 47.8 | 23 | Aug 2017 |
| | *barcode removed* | | 319.4 | 88.5 | 44.2 | | |
| | *Molecule length >=100K* | | 3.8 | 1.1 | 0.5 | | |
| ***P. kuhlii*** | *Raw* | 4 | 327.4 | 98.9 | 55.7 | 19 | Nov 2017 |
| | *barcode removed* | | 327.4 | 91.3 | 51.4 | | |
| | *Molecule length >=100K* | | 4.6 | 1.4 | 0.8 | | |
| ***R. ferrumequinum*** | *Raw* | 4 | 345.0 | 104.2 | 50.2 | 30 | Nov 2017 |
| | *barcode removed* | | 345.0 | 96.2 | 46.4 | | |
| | *Molecule length >=100K* | | 5.4 | 1.6 | 0.8 | | |
| ***P. discolor*** | *Raw* | 8 | 788.9 | 236.7 | 113.0 | 13 | Mar 2018 |
| | *barcode removed* | | 788.9 | 218.5 | 104.3 | | |
| | *Molecule length >=100K* | | 18.5 | 5.6 | 2.7 | | |
| ***R. aegyptiacus*** | *Raw* | 8 | 365.1 | 110.2 | 58.2 | 94 | Oct 2018 |
| | *barcode removed* | | 365.1 | 101.8 | 53.8 | | |
| | *Molecule length >=100K* | | 167.9 | 50.7 | 26.8 | | |
| ***M. molossus*** | *Raw* | 8 | 354.8 | 107.2 | 46.2 | 128 | Oct 2018 |
| | *barcode removed* | | 354.8 | 99.0 | 42.7 | | |
| | *Molecule length >=100K* | | 226.4 | 68.4 | 29.5 | | |

**Table S31: Statistics of Bionano dataset.** The filtered data set were used for the *de novo* Bionano assembly and consists of molecules with a minimum length of 150 kb and a minimum of 9 label sites.

| | | Technology | Number of Molecules (M) | Total Length (Gbp) | Estimated Coverage | Average Molecule Length (Kbp) | Finish Date Where |
|---|---|---|---|---|---|---|---|
| ***M. myotis*** | *Raw* | *DLE1* | 27.7 | 1960 | 979 | 71 | Nov 2018 |
| | *Filtered* | | 2.0 | 396 | 198 | 195 | Ploen |
| ***P. kuhlii*** | *Raw* | *DLE1* | 17.9 | 1419 | 799 | 79 | Nov 2018 |
| | *Filtered* | | 1.9 | 460 | 259 | 242 | Ploen |
| ***R. ferrumequinum*** | *Raw* | *BSPQI* | 6.4 | 547 | 263 | 85 | Dec 2017 |
| | *Filtered* | | 1.0 | 313 | 151 | 308 | Rockefeller |
| | *Raw* | *BSSSI* | 12.0 | 1046 | 504 | 89 | Dec 2017 |
| | *Filtered* | | 2.1 | 597 | 288 | 286 | Rockefeller |
| ***P. discolor*** | *Raw* | *BSPQI* | 6.7 | 764 | 365 | 114 | Jun 2017 |
| | *Filtered* | | 1.8 | 480 | 229 | 265 | Rockefeller |
| | *Raw* | *BSSSI* | 2.7 | 315 | 150 | 118 | Jun 2017 |
| | *Filtered* | | 0.7 | 186 | 89 | 151 | Rockefeller |
| ***R. aegyptiacus*** | *Raw* | *DLE1* | 2.4 | 410 | 216 | 169 | Feb 2019 |
| | *Filtered* | | 1.0 | 320 | 169 | 309 | Dresden |
| ***M. molossus*** | *Raw* | *DLE1* | 9.2 | 948 | 409 | 102 | Oct 2017 |
| | *Filtered* | | 2.0 | 462 | 199 | 234 | Ploen |

**Table S32: Statistics of Bionano restriction map assemblies.**

| | Technology | Number of Maps | Total Length (Gbp) | Average Map Length (Mbp) | N50 Map Length (Mbp) |
|---|---|---|---|---|---|
| ***M. myotis*** | *DLE1* | 342 | 2.22 | 6.5 | 44.8 |
| ***P. kuhlii*** | *DLE1* | 474 | 1.78 | 3.7 | 13.4 |
| ***R. ferrumequinum*** | *BSPQI* | 1390 | 2.12 | 1.5 | 2.3 |
| | *BSSSI* | 812 | 2.36 | 2.9 | 6.3 |
| ***P. discolor*** | *BSPQI* | 1193 | 2.42 | 2.0 | 3.1 |
| | *BSSSI* | 1345 | 2.07 | 1.5 | 2.4 |
| ***R. aegyptiacus*** | *DLE1* | 58 | 1.96 | 33.8 | 88.5 |
| ***M. molossus*** | *DLE1* | 222 | 2.56 | 11.6 | 80.4 |

**Table S33: Statistics of Hi-C dataset.** The estimated coverage was computed by using the final assembly sizes (including gap size).

| | Number of Cycles | Number of Reads (M) | Total Base Pairs (Gbp) | Estimated Coverage | Finish Date |
|---|---|---|---|---|---|
| ***M. myotis*** | 80 PE | 376.4 | 30.1 | 15.0 | Jun 2017 |
| ***P. kuhlii*** | 80 PE | 389.8 | 31.2 | 17.6 | May 2017 |
| ***R. ferrumequinum*** | 150 PE | 306.5 | 46.3 | 22.3 | Sep 2017 |
| ***P. discolor*** | 150 PE | 1331.4 | 199.7 | 95.3 | Feb 2018 |
| ***R. aegyptiacus*** | 150 PE | 924.0 | 139.5 | 73.7 | Dec 2018 |
| ***M. molossus*** | 150 PE | 975.1 | 147.2 | 63.5 | Feb 2019 |

**Table S34: The genomes after read assembly.** (all lengths in Mbp, NG50 correspond to *post hoc* genome size of the final assemblies.)

| Species | Primary Contigs | | | Alternate Contigs | | | Discarded Contigs | | |
|---|---|---|---|---|---|---|---|---|---|
| | Number | Total Length | NG50 | Number | Total Length | N50 | Number | Total Length | N50 |
| ***M. myotis*** | 598 | 1,976 | 11.80 | 671 | 55 | 0.10 | 406 | 22 | 0.06 |
| ***P. kuhlii*** | 527 | 1,765 | 10.24 | 624 | 53 | 0.10 | 270 | 13 | 0.05 |
| ***R. ferrumequinum*** | 324 | 2,056 | 21.74 | 67 | 58 | 0.10 | 120 | 7 | 0.07 |
| ***P. discolor*** | 421 | 2,056 | 16.15 | 234 | 20 | 0.10 | 245 | 14 | 0.06 |
| ***R. aegyptiacus*** | 260 | 1,866 | 21.74 | 108 | 10 | 0.11 | 141 | 9 | 0.07 |
| ***M. molossus*** | 396 | 2,261 | 21.58 | 565 | 89 | 0.18 | 205 | 15 | 0.08 |

**Table S35: The genomes after Bionano scaffolding.** (all lengths in Mbp; NG50 correspond to *post hoc* genome size of the final assemblies. Δ Number refers to the number of contig breaks. Δ NG50 refers to difference between the NG50 of Bionano contigs and NG50 of locally-phased contigs.)

| Species | Scaffolds | | | | Δ Primary Contigs | |
|---|---|---|---|---|---|---|
| | Number | Total length | Maximum length | NG50 length | Δ Number | Δ NG50 |
| ***M. myotis*** | 119 | 2,003 | 113 | 62.10 | +9 | +0.00 |
| ***P. kuhlii*** | 223 | 1,776 | 92 | 48.91 | +8 | +0.00 |
| ***R. ferrumequinum*** | 49 | 2,076 | 201 | 96.92 | +16 | +0.00 |
| ***P. discolor*** | 99 | 2,095 | 121 | 47.75 | +27 | -1.00 |
| ***R. aegyptiacus*** | 49 | 1,951 | 178 | 93.67 | +10 | +0.00 |
| ***M. molossus*** | 76 | 2,319 | 132 | 84.84 | +11 | +0.00 |

**Table S36: Statistics of 6 bat genomes after Hi-C scaffolding.** (all lengths in Mbp, NG50 correspond to *post hoc* genome size of the final assemblies. Δ Number refers to the number of contig breaks. Δ NG50 refers to difference between the NG50 of Hi-C contigs and NG50 of Bionano contigs.)

| Species | Scaffolds | | | | Δ Primary Contigs | |
|---|---|---|---|---|---|---|
| | Number | Total Length | Maximum Length | NG50 | Δ Number | Δ NG50 |
| ***M. myotis*** | 100 | 2,003 | 218 | 89.76 | +22 | +0.00 |
| ***P. kuhlii*** | 202 | 1,776 | 197 | 80.24 | +62 | +0.00 |
| ***R. ferrumequinum*** | 48 | 2,076 | 128 | 90.45 | +7 | +0.00 |
| ***P. discolor*** | 64 | 2,095 | 215 | 104.13 | +3 | +0.00 |
| ***R. aegyptiacus*** | 40 | 1,951 | 186 | 121.83 | +2 | +0.00 |
| ***M. molossus*** | 67 | 2,319 | 230 | 100.25 | +5 | +0.00 |

**Table S37: Number of gene evidence separated by type of evidence that were used to annotate coding genes in the genomes of the six bats.** * the reference species in this projection was *Myotis lucifugus* (Ensembl gene annotation), while for all other projections, we used our *Myotis myotis* gene annotation

| Evidence | Molossus | Myotis | Phyllostomus | Pipistrellus | Rhinolophus | Rousettus |
|---|---|---|---|---|---|---|
| Human Projections | 76,605 | 76,781 | 76,363 | 74,627 | 77,238 | 76,670 |
| Mouse Projections | 49,206 | 50,555 | 49,750 | 48,964 | 49,916 | 49,319 |
| Bat Projections | 51,471 | 20241 * | 50,790 | 53,358 | 49,862 | 49,222 |
| TAMA filtered transcripts | 25,046 | 29,148 | 16,326 | 19,236 | 25,099 | 43,004 |
| FLNC, ANGEL Positive transcript | 62,303 | 61,638 | 107,272 | 28,398 | 25,866 | 87,623 |
| GenomeThreader Alignments | 8,449 (stringent parameters), 18,556 (lax parameters) | 23,787 | 14,372 | 14,132 | 25,795 | 13,192 |
| Augustsus single genome mode | 64,664 | 57,976 | 51,137 | 39,673 | 42,882 | 44,162 |
| Augustus CGP | 24,729 | 23,800 | 24,067 | 21,100 | 22,001 | 23,373 |

**Table S38: Base pair counts and genome proportion estimates of transposable element classes in each examined taxon.**
**(See separate file)**

**Table S39: The library of viral protein sequences used for endogenous viral elements (EVE) analysis.**
**(See separate file)**

**Table S40: The probes of viral proteins gag, pol and env that were used to identify the endogenous retrovirus sequences in 6 bat genomes.**

| Family | Abbreviation | Name | POL | ENV | GAG |
|---|---|---|---|---|---|
| Alpharetroviruses | ALV | Avian Leukosis Virus | AJG42161.1 | AJG42162.1 | AJG42160.1 |
| | RSV | Rous Sarcoma Virus | CAA48535.1 | CAA48536.1 | CAA48534.1 |
| Betaretroviruses | SRV | Simian retrovirus 2 | ATN28189.1 | ATN28190.1 | ATN28187.1 |
| | JSRV | Jaagsiekte sheep retrovirus | AAD45226.1 | NP_041188.1 | AAA89180.1 |
| Gammaretroviruses | DrERV | Desmodus rotundus endogenous retrovirus | AJR27940.1 | AJR27937.1 | AJR27933.1 |
| | PoERV | Porcine endogenous retrovirus | AAL38193.1 | CAA76583.1 | ADG27335.1 |
| | KoRV | Koala Retrovirus | YP_009513211.1 | YP_009513212.1 | AAF15097.1 |
| Deltaretrovirus | BLV | Bovine leukemia virus | BAA00544.1 | ALB75304.1 | AAC82585.1 |
| | HTLV | Human T-lymphotropic virus 2 | AAD34842.1 | AAD34843.1 | AAB59884.1 |
| Epsilonretroviruses | WEHV | Walleye epidermal hyperplasia virus 1 | AAD30048.1 | AAD30049.1 | AAD30047.1 |
| Lentiviruses | HIV-1 | Human immunodeficiency virus 1 | NP_789740.1 | AAC82596.1 | AAC82593.1 |
| | FIV | Feline immunodeficiency virus | CAA40318.1 | AAB59940.1 | AAB59936.1 |
| Spumaretroviruses | BFV | Bovine foamy virus | AFR79239.1 | AAB68771.1 | AAB68769.1 |
| | FFV | Feline foamy virus | CAA11581.1 | CAA70076.1 | CAA70074.1 |

**Table S41: Divergence time estimates (in millions of years) using r8s.** Their respective nodes are relative to tree topology 1.

| Node | Divergence Time (Mya) |
|---|---|
| Eutherian Root | 99.37 |
| Atlantogenata | 93.26 |
| Afrotheria | 71.7 |
| Paenungulata | 59.2 |
| Tubulidentata+Afrosoricida | 68.91 |
| Boreoeutheria | 94.67 |
| Euarchontoglires | 82.82 |
| Primates | 66 |
| Haplorhini | 61.26 |
| Simiiformes | 29.21 |
| Catarrhini | 20.55 |
| Hominidae | 10 |
| Homininae | 5.73 |
| Pan+Homo | 5.11 |
| Saimiri+Callithrix | 12.93 |
| Strepsirrhini | 48.84 |
| Glires | 81.85 |
| Rodentia+Lagomorpha | 78.81 |
| Lagomorpha | 50.76 |
| Rodentia | 66 |
| Hystricomorpha | 64.86 |
| Muroidea | 34.86 |
| Murinae | 21.61 |
| Microtus+Cricetulus | 28.24 |
| Heterocephalidae+Caviidae | 37.42 |
| Laurasiatheria | 88.14 |
| Eulipotyphla | 82.45 |
| Erinaceidae+Soricidae | 76.85 |
| Scrotifera | 77.26 |
| Carnivora+Pholidota+Cetartiodactyla+Perissodactyla | 74.67 |
| Cetartiodactyla+Perissodactyla | 73.14 |
| Cetartiodactyla | 55.86 |
| Sus+Bos+Cetacea | 52.75 |
| Bos+Cetacea | 47.52 |
| Cetacea | 34 |
| Perissodactyla | 55.5 |
| Carnivora+Pholidota | 69.9 |
| Pholidota | 8.03 |
| Carnivora | 43 |
| Caniformia | 34.64 |
| Arctoidea | 27.12 |
| Leptonychotes+Mustela | 25.53 |
| Chiroptera | 63.38 |
| Yangochiroptera | 52.78 |
| Molossus+Vespertilionidae | 47.39 |
| Vespertilionidae | 24.7 |
| Yinpterochiroptera | 56.63 |

**Table S42: Investigation of conserved 1-to-1 single-copy miRNA genes in 6 bat species compared to other 42 mammalian taxa.** The conservativeness of mature miRNA and their seed regions was manually curated. 5p and 3p indicate the coordinates of 5p & 3p mature miRNA in multiple alignments. The empirical expression data were based on the miRBase (release 22). The missing values indicate that the miRNA does not have either 5p or 3p mature sequences.
**(See separate file)**
