## Supplementary figures and images for "Six new reference-quality bat genomes illuminate the molecular basis and evolution of bat adaptations"

### Supplementary Figure S14

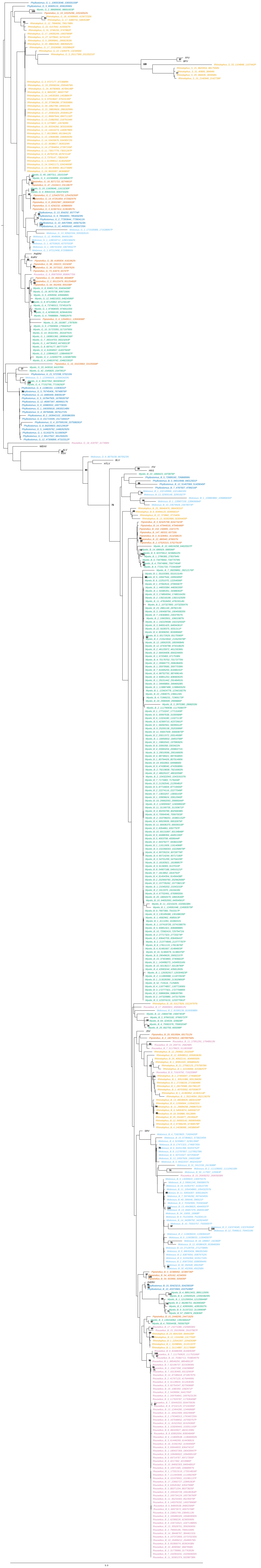
